# NFIB influences progenitor competence in maturation of GABAergic neurons in mice

**DOI:** 10.1101/2024.03.18.585524

**Authors:** Ann Rose Bright, Yana Kotlyarenko, Florian Neuhaus, Diana Rodrigues, Chao Feng, Christian Peters, Ilaria Vitali, Elif Dönmez, Michael H. Myoga, Elena Dvoretskova, Christian Mayer

## Abstract

Diverse types of GABAergic projection neurons and interneurons of the telencephalon derive from progenitors in a ventral germinal zone, called the ganglionic eminence. Using single-cell transcriptomics, chromatin accessibility profiling, lineage tracing, birthdating, heterochronic transplantation, and perturbation sequencing in mouse embryos, we investigated how progenitor competence influences the maturation and differentiation of these neurons. We found that the progression of neurogenesis over developmental time shapes maturation competence in ganglionic eminence progenitors, influencing how they progress into mature states. In contrast, differentiation competence, which defines the ability to produce diverse transcriptomic identities, remains largely unaffected by the stages of neurogenesis. Chromatin remodeling alongside a NFIB-driven regulatory gene module influences maturation competence in late-born neurons. These findings provide key insights into how transcriptional programs and chromatin accessibility govern neuronal maturation and the diversification of GABAergic neuron subtypes during neurodevelopment.

## Introduction

The development of diverse neuronal types is orchestrated with temporal, spatial, and numerical precision (Bandler and Mayer, 2023). It depends on the competence of neuronal progenitor cells, regulated by gene regulatory programs that govern cell differentiation and maturation (Bonnefont and Vanderhaeghen, 2021; Farnsworth and Doe, 2017). This study dissects two key aspects of progenitor competence: maturation competence and differentiation competence (Fig. 1a). Maturation competence, as defined here, refers to the potential of progenitors to generate postmitotic progeny with different maturation states. Differentiation competence describes the potential of progenitors to produce a diverse array of neuronal types, each characterized by different gene expression profiles. How these facets of progenitor competence are regulated and coordinated during neurogenesis remains an important question in neuronal development.

**Figure 1:**
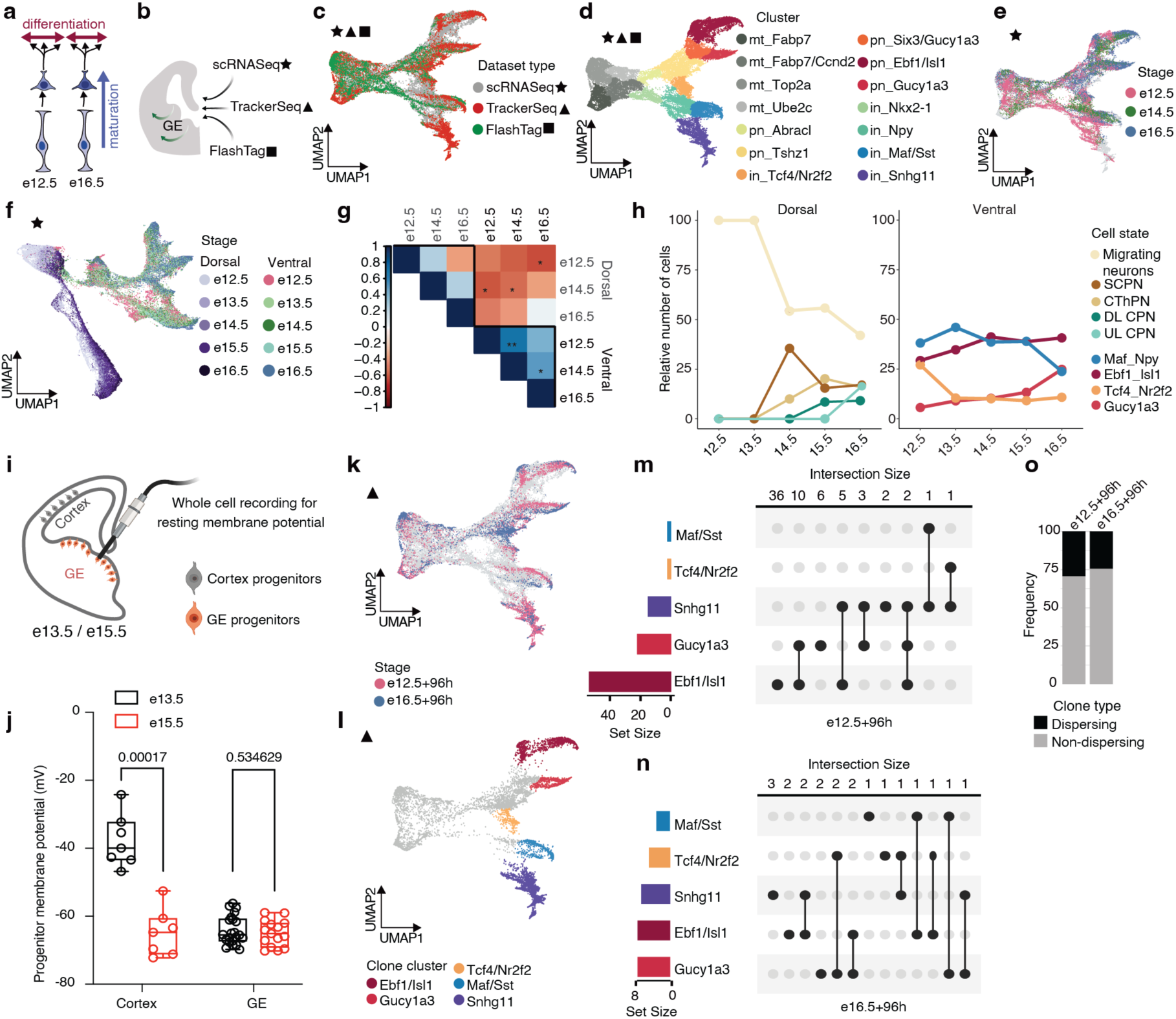
Stable differentiation competence in GABAergic progenitors. **a**, Schematic illustrating the difference between maturation and differentiation. **b**, Summary of methods used to investigate the competence of progenitors located in the GE. **c**, UMAP plot showing single cells derived from scRNA-seq, TrackerSeq, and FlashTag datasets aligned in Monocle3; Different symbols and colors corresponding to datasets. **d**, UMAP plot showing GABAergic cells; each color representing a different cluster; mt: mitotic; pn: projection neuron; in: interneuron. **e**, UMAP plot showing scRNA-seq datasets, with colors indicating various collection stages. **f**, UMAP plot showing single cells from ventral (GABAergic lineage) and dorsal (glutamatergic lineage) telencephalon, with colors indicating various collection stages. **g**, Pearson’s correlation plot between dorsal and ventral progenitors at different developmental stages; * *P*<0.05, ** *P*<0.01. **h**, Line plot showing relative cell number of dorsal (left) and ventral (right) postmitotic neuronal states across stages. The annotation of dorsal cell states is derived from the original publication. SCPN: subcerebral projection neuron; CThPN: corticothalamic projection neuron; DL CPN: deep layer callosal projection neuron; UL CPN: upper layer callosal projection neuron. **i**, Schematic illustrating whole-cell recording for resting membrane potential. **j**, Box plots showing the membrane potential in cortical and GE progenitors at e13.5 and e15.5 (two-sided t-test). **k**, UMAP plot showing TrackerSeq barcoded cells, each color representing a stage of IUE; IUE at e12.5 and e16.5, scRNA-seq after 96 hours. **l**, UMAP plot showing cell states at the branches used for clone grouping. **m**, Upset plot showing clonal intersections in TrackerSeq_e12.5 + 96h_. **n**, Upset plot showing clonal intersections in TrackerSeq_e16.5 + 96h_. **o**, Barplot showing the frequency of dispersing and non-dispersing clones in TrackerSeq_e12.5 + 96h_ and TrackerSeq_e16.5 + 96h_.

Excitatory neurons of the mammalian cerebral cortex are developmentally derived from proliferative zones in the dorsal telencephalon (Puelles et al., 2000). During neurogenesis, the competence of progenitors that give rise to excitatory neurons changes over time, influencing the sequence of neuronal differentiation events and guiding these neurons toward their final fate (Di Bella et al., 2021; Telley et al., 2019; Vitali et al., 2018). Less is known about how mitotic progenitor competence regulates the development of inhibitory neurons, which originate from the ganglionic eminences (GE) in the ventral telencephalon (Wonders and Anderson, 2006; Gelman et al., 2011; Anderson et al., 2001). During neurogenesis, mitotic progenitors in the ventricular zone (VZ) of the GE divide to produce postmitotic precursors. These precursors begin to mature and differentiate in the GE, and these processes continue as they migrate to different regions of the telencephalon and integrate into neuronal circuits. While there has been progress in correlating gene expression dynamics with chromatin accessibility (Allaway et al., 2021; Fleck et al., 2021; Janssens et al., 2021; Gonzalez-Blas et al., 2023), the impact of mitotic progenitor competence on the differentiation and maturation of inhibitory neurons remains poorly understood.

Here, we explored the role of progenitor competence in forebrain inhibitory neuron development using a range of techniques, including FlashTag (FT) birth labeling (Govindan et al., 2018), perturbation sequencing (Dvoretskova et al., 2024), and single-cell lineage analysis (Bandler et al., 2022). We show that, in contrast to progenitors in the cortex, progenitors in the GE maintain their competence to generate a consistent set of postmitotic cell states, as demonstrated by lineage tracing and the comparisons of isochronic cohorts of early and late-born inhibitory neurons. However, early and late born cohorts differ in the progression rate at which they move through maturation. These stage-specific differences originated from variations in chromatin accessibility profiles, as demonstrated by enhancer-driven gene regulatory networks (eGRNs). NFIB emerged as a key transcription factor (TF) in regulatory gene modules active in late-born progenitors, likely driving the observed changes in competence, as confirmed by perturbation sequencing and cleavage under target and release under nuclease (CUT&RUN) experiments. Finally, heterochronic transplantations revealed that maturation competence is influenced by the extrinsic environment. An interactive web-based resource is available for exploring scRNA-seq, scATAC-seq, and eGRN datasets, including comparisons between the GE and dorsal cortical neurogenesis (http://141.5.108.55:3838/mind_shiny/). Our findings demonstrate that both maturation and differentiation competence of progenitors are key determinants of neuronal development, with distinct roles in shaping dorsal and ventral neuronal lineages.

## Results

To investigate how progenitor competence influences neuronal maturation and differentiation in GABAergic lineages, we analyzed neuronal populations generated at different stages of neurogenesis (embryonic day (e) 12.5–e16.5) using distinct approaches, including scRNA-seq (★), barcode lineage tracing (▲) (Bandler et al., 2022), and fluorescent birthdating (■; Fig. 1a,b, Extended Data Fig. 1a,b, Supplementary Fig. 1a,b) (Govindan et al., 2018).

For scRNA-seq (★), we collected embryos from Dlx5/6-Cre::tdTomato mice at e12.5, e14.5, and e16.5, in which GABAergic neurons are labeled with a fluorescent reporter (Monory et al., 2006). From the same brains, cortical and striatal regions were manually dissected, dissociated and tdTomato-positive (tdTomato^+^) cells were enriched by fluorescence-activated cell sorting (FACS). Cells from the GE (without FACS enrichment) and tdTomato^+^ cells from the cortex and striatum (with FACS enrichment) were pooled to capture developmental states ranging from mitotic progenitors to postmitotic precursors and subjected to scRNA-seq (Extended Data Fig. 1c).

For barcode lineage tracing (▲), we devised a published method called TrackerSeq, which uses heritable DNA barcodes to label individual progenitors and their progeny followed by multiplexed scRNA-seq (Bandler et al., 2022; Dvoretskova et al., 2024). We targeted progenitors in the GE at e16.5 with TrackerSeq plasmids via *in utero* electroporation (IUE), FACS-enriched electroporated cells 96 h later, and performed scRNA-seq (TrackerSeq_e16.5 + 96h_). In our analysis, we also included a published TrackerSeq dataset, in which TrackerSeq plasmids were electroporated at e12.5, and the targeted cells were collected 96 h later (TrackerSeq_e12.5 + 96h_; Extended Data Fig. 1d)(Bandler et al., 2022). For birthdating (■), we used a technique called FlashTag (FT), which labels isochronic cohorts of cells with the fluorescent dye carboxyfluorescein succinimidyl ester (CFSE) (Govindan et al., 2018). In this method, mitotic cells layering the ventricle are labelled during the M phase of the cell cycle and maintain high fluorescence when leaving the cell cycle.

We injected CFSE into the ventricles of e12.5 and e16.5 wild-type embryos. Six hours later, we anatomically dissected the GE, FACS enriched FT labelled (FT^+^) cells and performed scRNA-seq (FT_e12.5 + 6h_ and FT_e16.5 + 6h_ respectively) (Extended Data Fig. 1e). At the same time point, coronal sections revealed CFSE+ cells in the VZ and subventricular zone (SVZ) (Govindan et al., 2018), representing isochronic cohorts transitioning through mitotic progenitor, intermediate progenitor (*Ascl1*), and postmitotic precursor stages (*Gad2*), as shown by RNAscope (Extended Data Fig. 1f-h). We also injected CFSE into the ventricles of e12.5 Dlx5/6-Cre::tdTomato mouse embryos, allowing for the collection of inhibitory neurons 96h post-injection from anatomically dissected cortical and striatal tissue following their migration. TdTomato^+^ and FT^+^ cells were enriched by FACS and scRNA-seq was performed (FT_e12.5 + 96h_; Extended Data Fig. 1e-h).

We pre-processed and merged datasets from all three methods (★, ▲, ■), using Seurat (Stuart et al., 2019), aligned the batches using Monocle3 (Trapnell et al., 2014; Haghverdi et al., 2018), and projected the data into a low-dimensional UMAP space (Fig. 1c). We then performed clustering (Fig. 1d, Extended Data Fig. 2a,b) and trajectory analyses using Monocle3 (Extended Data Fig. 2c), which learns the sequence of gene expression changes and uses a diffusion pseudotime algorithm to identify developmental trajectories. Consistent with previous work, clusters, and trajectories represented a continuum of cell state transitions during cellular maturation and differentiation (Extended Data Fig. 2d) (Mayer et al., 2018; Bandler et al., 2022; Lee et al., 2022; Rhodes et al., 2022; Lim et al., 2018). We manually annotated clusters based on marker gene expression, identifying them as mitotic apical progenitors (APs; *Fabp7*), mitotic basal progenitors (BPs; *Fabp7*, *Ccnd1*, *Top2a*, and *Ube2c*), GABAergic projection neuron precursors (PNs: *Abracl*, *Tshz1*, *Six3*, *Gucy1a3*, *Ebf1*, and *Isl1*), and GABAergic interneuron precursors (INs: *Nkx2-1*, *Npy*, *Maf*, *Sst*, and *Snhg11*; Fig. 1d; Extended Data Fig. 2e). After cell cycle exit, a common trajectory diverged, giving rise to distinct precursor states of PNs and INs. Each of these trajectories underwent subsequent divisions, resulting in multiple postmitotic precursor states that have been shown in previous studies to be linked to adult cell types (Mayer et al., 2018; Bandler et al., 2022) (see Methods) (Extended Data Fig. 2e,f; Supplementary Fig. 2a,b).

To dissect neuronal maturation and differentiation during early stages of development, we first explored the scRNA-seq (★) data in our combined single cell trajectory (Fig. 1e) and calculated the sequential patterns of gene expression along the Monocle3 pseudotime trajectory. Surprisingly, the dynamic expression of TFs along the pseudotime trajectory was highly conserved across different stages of neurogenesis (e12.5, e14.5 and e16.5; Supplementary Fig. 2c).

### The differentiation competence differs between dorsal and ventral lineages

The identified developmental progression differs from dorsal lineages (Di Bella et al., 2021; Telley et al., 2019). To investigate this, we focused on differences in neurogenesis between dorsal and ventral lineages. We merged and aligned the scRNA-seq (★) datasets and published data of GABAergic neurons from e13.5 and e15.5 (Bandler et al., 2022) and glutamatergic neurons from e12.5 to e16.5 (Di Bella et al., 2021) using Monocle3 (Fig. 1f, Supplementary Fig. 3a). In the UMAP representation, dorsal and ventral lineages overlapped at the level of APs but separated into distinct trajectories at the stage of BPs and postmitotic precursors (Supplementary Fig. 3b) (Moreau et al., 2021). When comparing successive developmental stages, cells of the dorsal lineage showed a sequential shift in the UMAP positioning, consistent with findings from previous studies (Telley et al., 2016; Di Bella et al., 2021). In contrast, cells of the ventral lineage largely overlapped across developmental stages (Fig. 1f). We identified genes associated with the emergence of inhibitory and excitatory neurons by selecting dynamic genes across pseudotime in the two lineages (see Methods). Only few genes overlapped between inhibitory and excitatory lineages, primarily at the initial pseudotime scores (Supplementary Fig. 3c,d; Supplementary Fig. 4a–d).

To quantify the temporal progression of dorsal and ventral progenitors, we calculated Pearson correlation coefficients between APs from each group, using highly variable genes (see Methods). Ventral progenitors showed higher correlation coefficients between successive stages of neurogenesis than dorsal progenitors, indicating less change in their gene expression profiles (Fig. 1g; Supplementary Fig. 3e). Furthermore, differential gene expression analysis of ventral progenitors across stages revealed that only a few genes were upregulated at later stages of neurogenesis (Supplementary Fig. 5a). Genes that were downregulated at later stages were primarily related to self-renewal (Supplementary Fig. 5a,b), in line with a change in balance between cell proliferation and differentiation during neurogenesis (Götz and Huttner, 2005). Next, we annotated postmitotic cells based on marker gene expression (ventral lineage) or published data (dorsal lineage) and quantified the proportion of cells in postmitotic precursor states across different developmental stages (Supplementary Fig. 5c). While the relative distribution of precursor states was similar across stages in ventral cells, it sequentially shifted in dorsal cells (subcerebral PN (SCPN) → corticothalamic PN (CThPN) → deep-layer callosal PN (DL CPN) → upper-layer callosal PN (UP CPN); Fig. 1h). Furthermore, we observed a similar trend when using fine-grained cluster annotation, that was inferred from the integrated dataset (Supplementary Fig. 5d,e).

In dorsal progenitors of the cortical VZ, bioelectrical processes have been shown to coordinate the temporal progression of developmental competence, despite these cells being nonexcitable (Vitali et al., 2018). Using whole-cell patch-clamp recordings in e12.5 to e15.5 cortical slices, Vitali et al. revealed a progressive membrane hyperpolarization over this period, regulating the timing of AP competence and associated neuronal diversity. To test whether a similar membrane potential progression occurs in VZ progenitors of the GE, we conducted whole-cell patch-clamp recordings from both cortical and GE progenitors at e13.5 and e15.5 (Fig. 1i). Our recordings confirmed the hyperpolarization in dorsal progenitors observed by Vitali et al., while the membrane potential of ventral progenitors in the GE remained stable between e13.5 and e15.5 (Fig. 1j, Supplementary Fig. 6a), highlighting a difference in developmental regulation between these regions.

Taken together, our findings reveal several marked differences between dorsal and ventral progenitors. While dorsal progenitors exhibit a temporal progression in differentiation competence and undergo hyperpolarization, ventral progenitors show more stable differentiation competence and unvaried membrane potential throughout neurogenesis, with GABAergic precursor states generated independently of developmental stages.

### Clonal divergence is maintained across neurogenesis

Our results so far suggest that, at the population level, progenitors in the GE can give rise to a similar set of precursor states throughout neurogenesis. To investigate whether the clonal progeny of individual progenitors can diverge into distinct precursor states, we next analyzed the barcode lineage tracing (▲) data in our combined dataset (Fig. 1k). We selected multicellular clones — i.e., clones containing multiple cells derived from a single progenitor — with cells located at the branch tips of the Monocle3 trajectory, where branch tips represent distinct developmental endpoints of the differentiation path (Fig. 1l-n, Supplementary Fig. 7a–d). We then grouped these clones based on whether their members were located within a single branch tip (non-dispersing clones) or across multiple branch tips (dispersing clones). Consistent with previous studies, a subset of the TrackerSeq_e12.5 + 96h_ clones dispersed into multiple branch tips (Bandler et al., 2022; Dvoretskova et al., 2024). Notably, a similar proportion of dispersing clones was found in TrackerSeq_e16.5 + 96h_ (Fig. 1m,n,o). The true proportion of dispersing clones is likely higher than observed, as TrackerSeq only partially recovers clones due to cell loss during sample preparation (Bandler et al., 2022).

Clonal resolution enables linking individual mitotic progenitor cells to the fate of their postmitotic progeny. We tested whether the transcriptome of mitotic cells correlates with the transcriptome of their postmitotic daughter cells (see Methods). Mitotic progenitor cells from non-dispersing clones did not show a stronger correlation with the transcriptomic profiles of their clonal progeny compared to randomly selected progenitor cells (Supplementary Fig. 8a–d).

Overall, the single-cell clonal analysis indicates that progenitor cells maintained a stable level of differentiation competence throughout neurogenesis, a conclusion that aligns with the results of the population-level analysis.

### Maturation dynamics differ between early and late born neurons

Next, we examined the maturation and differentiation of postmitotic cells at different stages of neurogenesis, using the fluorescent birthdating (■) data in our combined single-cell trajectory (Fig. 2a). FT^+^ cohorts collected six hours after CFSE application (FT_e12.5 + 6h_ and FT_e16.5 + 6h_) contained mitotic progenitors as well as early postmitotic neuronal precursors. Ninety-six hours after CFSE application (FT_e12.5 + 96h_), FT^+^ cohorts exclusively contained postmitotic cells (Extended Data Fig. 3a), consistent with the notion that FT marks isochronic cohorts of cells that exit the cell cycle shortly after CFSE application (Telley et al., 2016; Mayer et al., 2018). The postmitotic fractions across all three conditions (FT_e12.5 + 6h_, FT_e12.5 + 96h_, FT_e16.5 + 6h_) included cells from the same precursor states, but with differences in their relative population sizes (Fig. 2b). The rarity of some states in our analysis likely reflects the varying maturation stages of isochronic cohorts at the time of capture. For example, cells in the FT_e12.5 + 6h_ cohort appear to be transitioning towards branch tips, as indicated by their intermediate positions on the UMAP-embedding (Extended Data Fig. 3b). States with a low abundance of cells in a particular cohort shared consistent gene-expression profiles with corresponding states in other cohorts (Extended Data Fig. 3c). Next, we quantified the Monocle3 pseudotime scores as a proxy for the degree of maturation acquired by the different FT^+^ cohorts. As expected, given its later collection, FT_e12.5 + 96h_ showed higher pseudotime scores than FT_e12.5 + 6h_. Strikingly, the pseudotime score of FT_e16.5 + 6h_ was markedly higher than that of FT_e12.5 + 6h_, even though both were collected after six hours (Fig. 2c). Next, we performed a differential gene expression (DGE) analysis between postmitotic cells of the six-hour cohorts (FT_e12.5 + 6h_ vs. FT_e16.5 + 6h_; Fig. 2d). Genes upregulated in FT_e16.5 + 6h_ overlapped with those upregulated in FT_e12.5 + 96h_ (Fig. 2e, Extended Data Fig. 3d,e). The intersection analysis of cohort marker genes (see Methods) further supported this result, revealing a higher overlap between FT_e16.5 + 6h_ and FT_e12.5 + 96h_ marker genes (Extended Data Fig. 3e). These findings suggest that late-born neurons reach a similar gene expression profile within six hours as early-born neurons within 96 hours. Many of the genes upregulated in FT_e16.5 + 6h_ were associated with the promotion of neuronal proliferation and migration (Table S1). Some of these genes were specifically linked to neuronal signalling pathways. Overall, our results using FT birthdating suggest that although newborn neurons at different stages transition into similar precursor states, the rate and extent of their maturation differ, with late-born neurons maturing more rapidly compared to early-born neurons.

**Figure 2:**
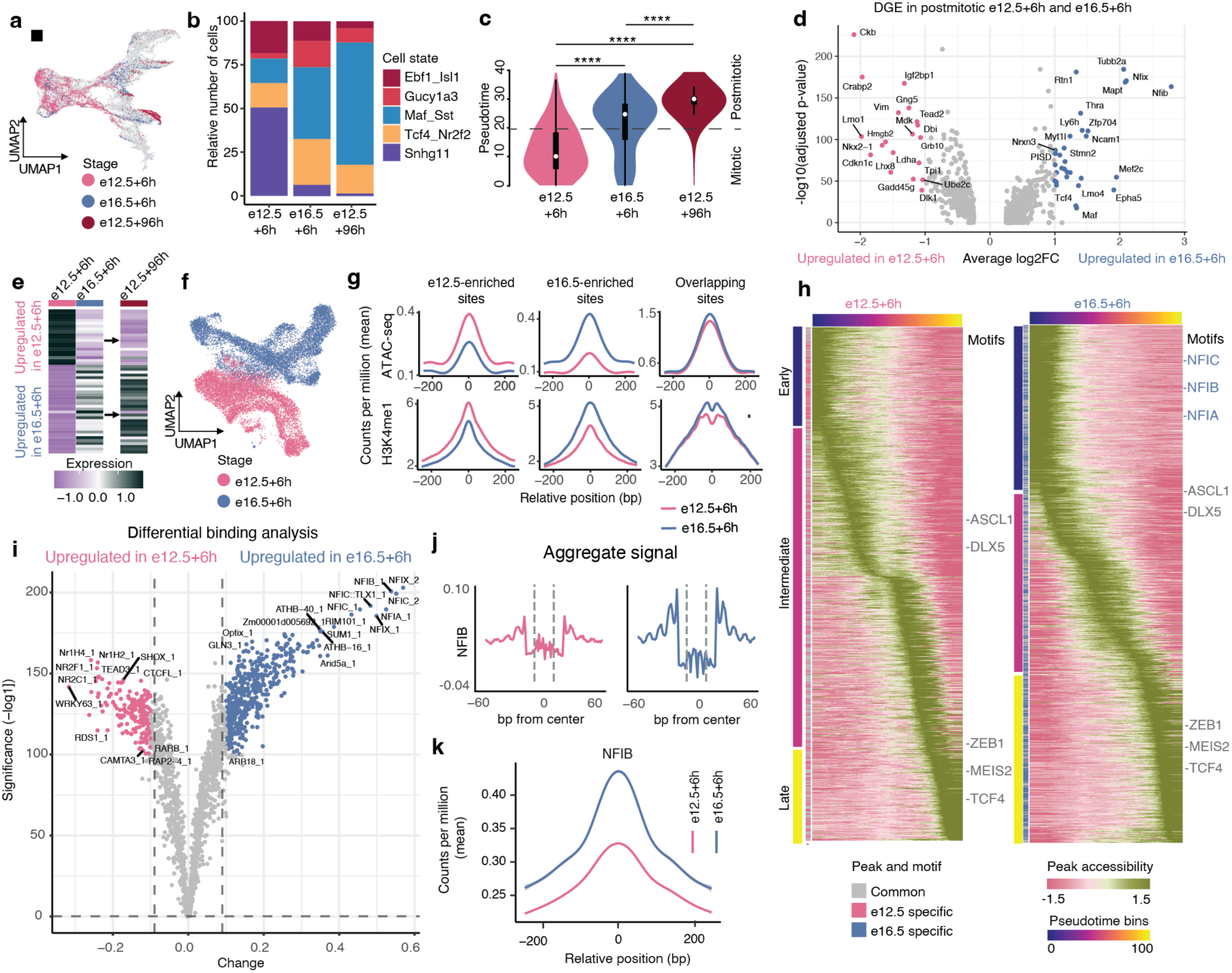
Timing of neurogenesis influences maturation competence. **a**, UMAP plot showing FlashTag (FT) datasets coloured by injection and collection stage; injection at e12.5 and e16.5, scRNA-seq after six hours or 96 hours. **b**, Barplot showing relative cell number of postmitotic neuronal states in FT_e12.5 + 6h_, FT_e16.5 + 6h_ and FT_e12.5 + 96h_. **c**, Violin plots showing the distribution of FT^+^ cells along the combined pseudotime trajectory, displayed for each condition; two-sided Wilcoxon rank sum test (**** adjusted *P*<0.001). The central point within the plot represents the median (50th percentile), the box represents the range between the first and third quartile (25th–75th percentile). **d**, Volcano plot displaying differential gene expression (DGE) in postmitotic cells of FT_e12.5 + 6h_ and FT_e16.5 + 6h_; | log_2_ (FC)| *>* 1, adjusted *% <* 0.05. **e**, Heatmap showing average scaled expression of differential genes in FT_e12.5 + 6h_ and FT_e16.5 + 6h_ postmitotic cells; visualized in all FT^+^ conditions. **f**, UMAP plot showing scATAC-seq datasets; FT injection at e12.5 and e16.5, followed by scATAC-seq after 6 hours. **g**, Coverage plot displaying scATAC-seq and H3K4me1 signal intensity for peak categories. X-axis is relative position (base pairs) and y-axis is counts per million (mean). **h**, Heatmap displaying the accessibility of cis-regulatory elements across pseudotime for FT_e12.5 + 6h_ and FT_e16.5 + 6h_. Peaks are divided into “initial”, “intermediate” and “late” based on accessibility profiles along pseudotime bins. Overlapping peaks are annotated in gray and unique peaks are annotated by stage-specific colors. Overlapping motifs are colored in gray and unique motifs are colored in blue. **i** Volcano plot displaying –log10(p-value) (x-axis) and differential binding score (y-axis) of significant transcription factors. Each dot represents a motif.**j**, Aggregate footprint profiles of NFIB in FT_e12.5 + 6h_ and FT_e16.5 + 6h_ **k**, Coverage plot showing chromatin accessibility dynamics at NFIB footprint sites for FT_e12.5 + 6h_ and FT_e16.5 + 6h_ datasets.

The observed maturation shift in the production of GABAergic neurons during neurogenesis may help adapt newly born neurons to the varying time available for network integration between early- and late-born neurons. We tested this hypothesis using electrophysiological recordings at P8, but were unable to definitively confirm or disprove it (Supplementary Results Supplementary Fig. 21 and Supplementary Fig. 22).

### Maturation shift is paralleled by changes in chromatin accessibility

To explore whether the different maturation dynamics we observed at embryonic stages are associated with changes at the chromatin level, we profiled chromatin accessibility using scATAC–seq (Buenrostro et al., 2015) on samples derived from FT^+^ cohorts in the GE (Extended Data Fig. 4a). We injected CFSE into the ventricles of e12.5 and e16.5 wild-type embryos, anatomically dissected the GE six hours later (FT_e12.5 + 6h_, FT_e16.5 + 6h_, respectively), enriched FT^+^ cells via FACS, and performed scATAC-seq (Extended Data Fig. 4b). Following sequencing, we mapped the paired-end reads to a reference genome and employed the ArchR framework (Granja et al., 2021) for quality control, as well as data processing steps such as dimensionality reduction, clustering and peak calling. Cell annotations were determined based on gene body accessibility patterns of cell state marker genes (Fig. 2f; Extended Data Fig. 4c,d).

In contrast to the scRNA-seq experiments (Fig. 1e), the isochronic cohorts FT_e12.5 + 6h_ and FT_e16.5 + 6h_ in the scATAC-seq experiment occupied distinct regions on the UMAP plot, both in mitotic and postmitotic cell states (Fig. 2f, Extended Data Fig. 4d). To identify and quantify the cis-regulatory elements (CREs) responsible for this separation, we independently conducted peak calling on FT_e12.5 + 6h_ and FT_e16.5 + 6h_. We then categorized the resulting peaks to identify genomic sites with e12.5-enriched peaks, e16.5-enriched peaks, and shared sites that were not stage-specific (’e12.5-sites’, ‘e16.5-sites’, and ‘overlapping-sites’, respectively; see Methods; Supplementary Fig. 9a). Subsequently, we computed the scATAC-seq fragment distribution and displayed the results in coverage plots. At e12.5-sites, we observed higher accessibility in FT_e12.5 + 6h_ than in FT_e16.5 + 6h_. Conversely, e16.5-sites had higher accessibility in FT_e16.5 + 6h_ than FT_e12.5 + 6h_ (Fig. 2g). The peak sets were divided by genomic region into promoters, distal, exonic, and intergenic regions (Extended Data Fig. 4e). At e12.5-sites and e16.5-sites, distal and intergenic regions represented the largest proportion of peaks. Together, this indicates that the chromatin accessibility undergoes marked changes between different stages of development, implying a dynamic process of chromatin remodeling that predominantly occurs at distal and intronic regions. To further categorize the identified sites as poised-active distal regulatory elements, we analyzed the distribution of H3K4me1 fragments, a well-established enhancer mark (Heintzman et al., 2007), utilizing forebrain ChIP-seq data from ENCODE (Gorkin et al., 2020). H3K4me1 profiles closely aligned with chromatin accessibility profiles (Fig. 2g). Specifically, e12.5-sites exhibited a stronger H3K4me1 signal at e12.5 compared to e16.5, and the contrary was observed for e16.5-sites. These observations suggest that distal regulatory elements are potentially maintained in a poised-active state and likely drive the stage-specific dynamics in chromatin accessibility.

To complement our earlier analysis (Fig. 2g) that identified e12.5- or e16.5-enriched sites, we performed a differential peak analysis (see Methods). This analysis resulted in 11,957 peaks differentially accessible at e12.5, 14,825 peaks differentially accessible at e16.5, and 122,129 non-significant peaks (Extended Data Fig. 4f). To visualize changes in chromatin accessibility, coverage plots were generated, revealing trends consistent with those observed using the previous peak set (Extended Data Fig. 4g), further validating the stage-specific changes in chromatin accessibility between e12.5 and e16.5.

To explore how the stage-specific accessibility of CREs relates to the maturation process, we used ArchR to assign a pseudotime score to cells, capturing their position along the maturation trajectory (from APs to BPs to precursor cells; Fig. 2h). We used ArchR to perform peak calling along the inferred trajectory and grouped the identified CREs into three main phases based on their accessibility profiles along pseudotime: “initial,” “intermediate,” and “late” CREs, corresponding broadly to APs, BPs, and precursor cells. We found more peaks in “initial” CREs at FT_e16.5 + 6h_ in respect to FT_e12.5 + 6h_, suggesting an early opening of additional regulatory elements in e16.5 progenitors (Supplementary Fig. 9b). To identify associated TFs, we subsequently conducted motif scanning on the peaks that were specific to the “initial”, “intermediate”, and “late” CREs at both stages. From this analysis, we identified both common and stage-specific motifs (Fig. 2h). Motifs of TFs associated with inhibitory neuron development, such as TCF4, MEIS2, EBF1, and ISL1 (Supplementary Fig. 2b), were detected at both stages. Conversely, several motifs from the NFI family (NFIA, NFIB, NFIC) were linked exclusively to “initial” CREs in FT_e16.5 + 6h_. The NFI TFs are known for regulating key steps during brain development (Zenker et al., 2019b), such as neural and glial cell differentiation (Bunt et al., 2017), neuronal migration (Heng et al., 2012), and maturation (Hickey et al., 2019).

DNA-binding proteins, like TFs, protect genomic regions from Tn5 integration during scATAC-seq sample preparation, creating a measurable “footprint” that indicates the binding patterns of TFs on chromatin. These footprints, thus predict the strength of TF binding (i.e. TF activity) and binding locations. We conducted a footprint analysis on the FT^+^ cohorts, using TOBIAS (Bentsen et al., 2020), and performed a differential binding analysis. Among the differential TFs, the NFI family demonstrated the most substantial and statistically significant increase of TF binding activity in FT_e16.5 + 6h_ (Fig. 2i). To visualize and evaluate this finding, we generated stage-specific aggregate footprint profiles for select TFs (Fig. 2j, Extended Data Fig. 4h). NFIX, NFIC, and NFIA displayed TF activity only in FT_e16.5 + 6h_ while NFIB displayed TF activity already in FT_e12.5 + 6h_, which significantly increased in FT_e16.5 + 6h_ (Fig. 2j; Extended Data Fig. 4h). This aligns with the gradual increase in gene expression patterns of NFI family TFs observed in the transcriptomic data (Supplementary Fig. 9c). Next, to assess whether sites where NFI family TFs bind (footprint sites) exhibit dynamic changes in accessibility, we calculated the fragment distribution within these regions. Coverage plots displayed a temporal increase in accessibility from FT_e12.5 + 6h_ to FT_e16.5 + 6h_ at NFIB, NFIA, NFIC and NFIX footprint sites (Fig. 2k, Supplementary Fig. 9d). Our findings suggest a link between specific TFs and the observed chromatin dynamics, underscoring their potential role in chromatin remodeling.

Taken together, these findings demonstrate that isochronic FT^+^ cohorts exhibit stage-specific chromatin accessibility, driven mainly at CREs. Furthermore, the NFI family of TFs plays a crucial role in characterizing FT_e16.5 + 6h_ cells based on their expression and early activation of regulatory elements.

### NFIB modulates the network underlying maturation competence

Our analysis of scATAC-seq profiles between FT_e12.5 + 6h_ and FT_e16.5 + 6h_ revealed that CREs, such as enhancers, are the primary source of heterogeneity. To infer enhancer-driven regulatory interactions, we applied SCENIC+ (Gonzalez-Blas et al., 2023) to integrate scRNA-seq and scATAC-seq data from FT_e12.5 + 6h_ and FT_e16.5 + 6h_. This approach enables the identification of genomic binding events (i.e., TFs binding to regulatory sites) and their links to downstream target genes. We grouped cells by collection stage (e12.5 and e16.5) and broad states (APs, BPs, and precursors), obtaining six groups in total (Extended Data Fig. 10a). After running the SCENIC+ pipeline with standard filtering, the resulting eGRN contained 147 TFs that bound on average 168 sites, with each site regulating one to three target genes (mean = 1.1; Supplementary Fig. 10b–d). The activity of regulatory modules (i.e., expression of TF and associated target genes) was scored in each cell using a previously established method (Aibar et al., 2017), and enriched modules for each group were identified (Supplementary Fig. 10e). Modules of canonical cell state markers were enriched in their respective groups: Hes5, Hes1, and Pax6 modules in APs (Ohtsuka et al., 2001; Thakurela et al., 2016); Ascl1 and Dlx2 modules in BPs (Raposo et al., 2015; Lindtner et al., 2019); and Dlx5 or Lhx6 modules in neuronal precursors (Lindtner et al., 2019; Liodis et al., 2007). We also found modules exhibiting patterns that were specific to certain cell states or developmental stages. For example, Nkx2-1 was active in BPs and precursor states, yet remained restricted to FT_e12.5 + 6h_. In contrast, modules of NFI family TFs were active across all cell states in FT_e16.5 + 6h_, with the highest activity in APs compared to BPs and precursor cells (Supplementary Fig. 10e).

Next, we inferred active gene regulatory interactions specific to the six groups by filtering the eGRNs for modules active in over 50% of cells within each group and applying an additional filter on the target genes based on expression level (see Methods). We obtained six subnetworks, each containing state and stage-specific modules of active TFs and target genes. We compared subnetworks of APs, BPs, and precursors across stages to infer dynamic modules and the regulatory interactions between them (see Methods). Specifically, we focused on subnetworks of APs to identify modules that maintain or modulate progenitor competence. Modules of canonical inhibitory neuron markers like Dlx1, Dlx2, and Arx (Lindtner et al., 2019; Colasante et al., 2015) were maintained throughout both stages, whereas modules linked to progenitor self-renewal, like Hmga2, Nr2f1, and Nr2f2 (Nishino et al., 2008; Bertacchi et al., 2020), were enriched in e12.5 APs (Fig. 3a). APs at e16.5 were characterized by enriched activity of Nfib, together with Nfia, Nfix, Pou3f2, Meis2 and Tcf4. In line with previous studies, NFIB acts as an upstream regulator of NFIX (Matuzelski et al., 2017), but also as an upstream regulator of NFIA, POU3F2, MEIS2, and TCF4 (Fig. 3a). The NFIB-led regulatory module was consistently enriched in BPs and precursors at e16.5 (Extended Data Fig. 5a-c), suggesting a role of NFIB as a central regulator.

**Figure 3:**
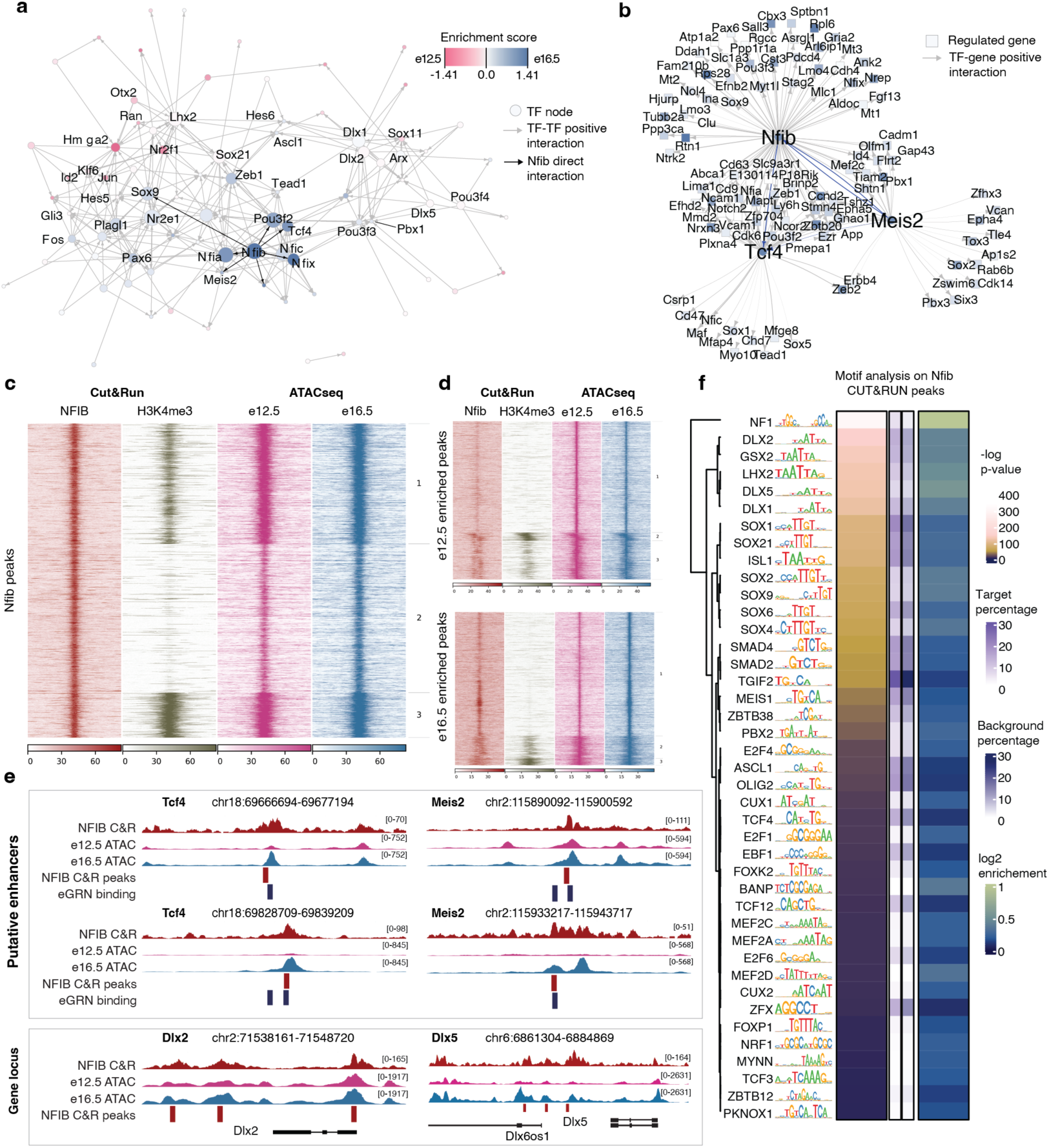
Nfib regulates a shift in gene-regulatory programs. **a**, An eGRN graph displaying positive interactions between TFs active in APs. Node color indicates enrichment score by stage and node size indicates the number of direct targets per TF. Select TFs are annotated. Direct interactions originating from Nfib are highlighted. **b**, An eGRN subgraph highlighting downstream targets of *Nfib*, *Tcf4* and *Meis2* at e16.5. *Nfib*, *Tcf4* and *Meis2* nodes are indicated by node shape. Interactions between *Nfib*, *Tcf4* and *Meis2* are highlighted. Node color reflects the enrichment score by stage. **c**, Heatmap displaying signal enrichment of NFIB peaks across datasets: NFIB and H3K4me3 CUT&RUN at e16.5 GE, and scATAC-seq at e12.5 and e16.5. **d**, Heatmap displaying signal enrichment of e12.5 and e16.5 enriched peaks across datasets: NFIB and H3K4me3 CUT&RUN at e16.5 GE, and scATAC-seq at e12.5 and e16.5.**e**, Genome browser tracks of putative enhancer regions for *Tcf4* and *Meis2* and gene loci for *Dlx2* and *Dlx5*, featuring NFIB CUT&RUN and scATAC-seq at e12.5 and e16.5. **f**, Enriched TF-motifs in NFIB CUT&RUN peaks. TFs are ordered by their p-value. For each TF, the motif logo, target-and background percentage and the resulting enrichment are shown. The dendrogram on the left shows the sequence similarity of motif logos.

Of particular interest to us were the interactions between NFIB with MEIS2 and TCF4, which are TFs specific to the development of inhibitory PNs and INs, respectively (Su et al., 2022; Wang et al., 2022; Dvoretskova et al., 2024). Moreover, these TFs share common direct target genes in different cell states of FT_e16.5 + 6h_ (Supplementary Fig. 11a), suggesting combinatorial binding of NFIB with TCF4 or MEIS2. To test this hypothesis, we used TFCOMB, a tool for identifying enriched TF binding motifs in chromatin accessibility data (Bentsen et al., 2022), to analyze peaks from FT_e12.5 + 6h_ and FT_e16.5 + 6h_ scATAC-seq datasets. Interestingly, NFIB was found to collaborate with these TFs at both stages, with higher cosine scores and increased binding events for NFIB-TCF4 and NFIB-MEIS2 in e16.5 peaks, suggesting a stage-specific enhancement of regulatory interactions that may drive late-stage maturation processes (Extended Data Fig. 5d,e). Next, using SCENIC+, we identified direct downstream target genes shared between NFIB, MEIS2, and TCF4 (Fig. 3b). Gene Ontology (GO) enrichment analysis of these downstream genes revealed roles in brain development, neuron fate specification, and the positive regulation of cell proliferation (Supplementary Fig. 11b). We then identified a group of genes exhibiting dynamic expression across the maturation trajectory and inferred their upstream TFs in FT^+^ cohorts, sorting TFs by the number of regulated maturation genes. Temporally conserved TFs such as DLX1 and LHX2, along with e16.5-specific TFs like NFIB and NFIX, regulated the largest number of genes, further supporting our previous observations (Extended Data Fig. 5g-h). To assess the functional relevance of the e12.5- and e16.5-enriched peaks, we quantified the proportion of these peaks that are contained in the eGRN. This analysis revealed substantial overlap: 71.43% of e12.5-enriched peaks and 70.93% of e16.5-enriched peaks were predicted to be part of TF – enhancer – target gene interactions (Supplementary Fig. 11c), suggesting that the majority of peaks are likely to have functional relevance.

Next, we performed CUT&RUN on unfixed, dissociated cells from the GE of e16.5 mice using an NFIB antibody, with IgG and H3K4me3 as controls, to identify and validate genomic targets of NFIB *in vivo*. Mapping and sample processing were carried out using widely used tools and pipelines (see Methods). MACS2 peak calling identified approximately 21,000 narrow peaks (p-value cutoff of 1 × 10^−4^) corresponding to NFIB binding relative to the IgG control. To investigate the relationship between NFIB binding and chromatin accessibility during development, we plotted signal intensities at NFIB-binding sites for NFIB, H3K4me3, and FT_e12.5 + 6h_ and FT_e16.5 + 6h_ scATAC-seq datasets (Fig. 3c). K-means clustering of these binding sites revealed three distinct clusters, all characterized by strong NFIB binding. Cluster 2 lacked H3K4me3 enrichment, suggesting these regions may represent non-promoter elements with increased chromatin accessibility at e16.5 relative to e12.5 (Supplementary Fig. 12b). In contrast, Clusters 1 and 3 showed intermediate to high levels of H3K4me3, indicating that many of these regions are promoters. To further examine NFIB binding at temporally dynamic peaks, we compared NFIB and H3K4me3 signal intensities at e12.5- and e16.5-enriched sites identified in our scATAC-seq data. NFIB binding was significantly higher at e16.5-enriched sites, whereas e12.5 sites showed markedly lower or no signal (Fig. 3d, Supplementary Fig. 12c,d). These findings support our hypothesis that NFIB is associated with chromatin remodeling at e16.5.

We validated predicted eGRN interactions of NFIB (e.g., NFIB-*Tcf4*, NFIB-*Meis2*) by confirming NFIB binding at predicted enhancers (Fig. 3e). Additionally, we observed NFIB binding at promoters of TFs involved in inhibitory neuron development, such as *Dlx2* and *Dlx5* (Fig. 3e). Furthermore, we quantified the fraction of eGRN predicted target regions of NFIB that was validated by NFIB CUT&RUN, by calculating the fraction of target regions with a binding event (43.7%). Motif analysis of NFIB peaks using HOMER (Heinz et al., 2010) displayed significant enrichment of additional TF motifs associated with inhibitory neuron development including DLX1/2/5, ISL1, SOX2, ASCL1, MEIS1/2 and TCF4 (Fig. 3f).

In summary, we observed gene-regulatory interactions that drive cell state- and stage-specific dynamics, with NFIB playing a leading role in late-born progenitors through direct and combinatorial binding at genes involved in maturation and differentiation.

### Influence of extrinsic environment on maturation competence

To investigate whether extrinsic environment influences maturation competence in APs at different stages, we conducted homo-and heterochronic transplantation experiments, assessing cell’s pseudotime scores and expression of genes downstream of NFIB, TCF4, and MEIS2. We injected CFSE into the ventricles of donor mouse embryos at e12.5 and e16.5. One hour later, we dissected and dissociated the GE, obtaining a cell suspension that included FT-labelled APs, unlabelled BPs, and unlabelled precursor cells. The cell suspension was transplanted homo- and hetero-chronically into host embryos via intraventricular injection (AP_e12.5→e12.5_, AP_e12.5→e16.5_, AP_e16.5→e16.5_, and AP_e16.5→e12.5_), as described by Oberst *et al*. (Oberst et al., 2019). Forty-eight hours after transplantation, we collected the GE from host embryos, isolated FT^+^ cells by FACS, and assessed their transcriptome using bulk RNA-seq (Fig. 4a; Extended Data Fig. 6a,b). By the time of collection, cells had already entered the tissue and begun migrating away from the VZ (Extended Data Fig. 6c).

**Figure 4:**
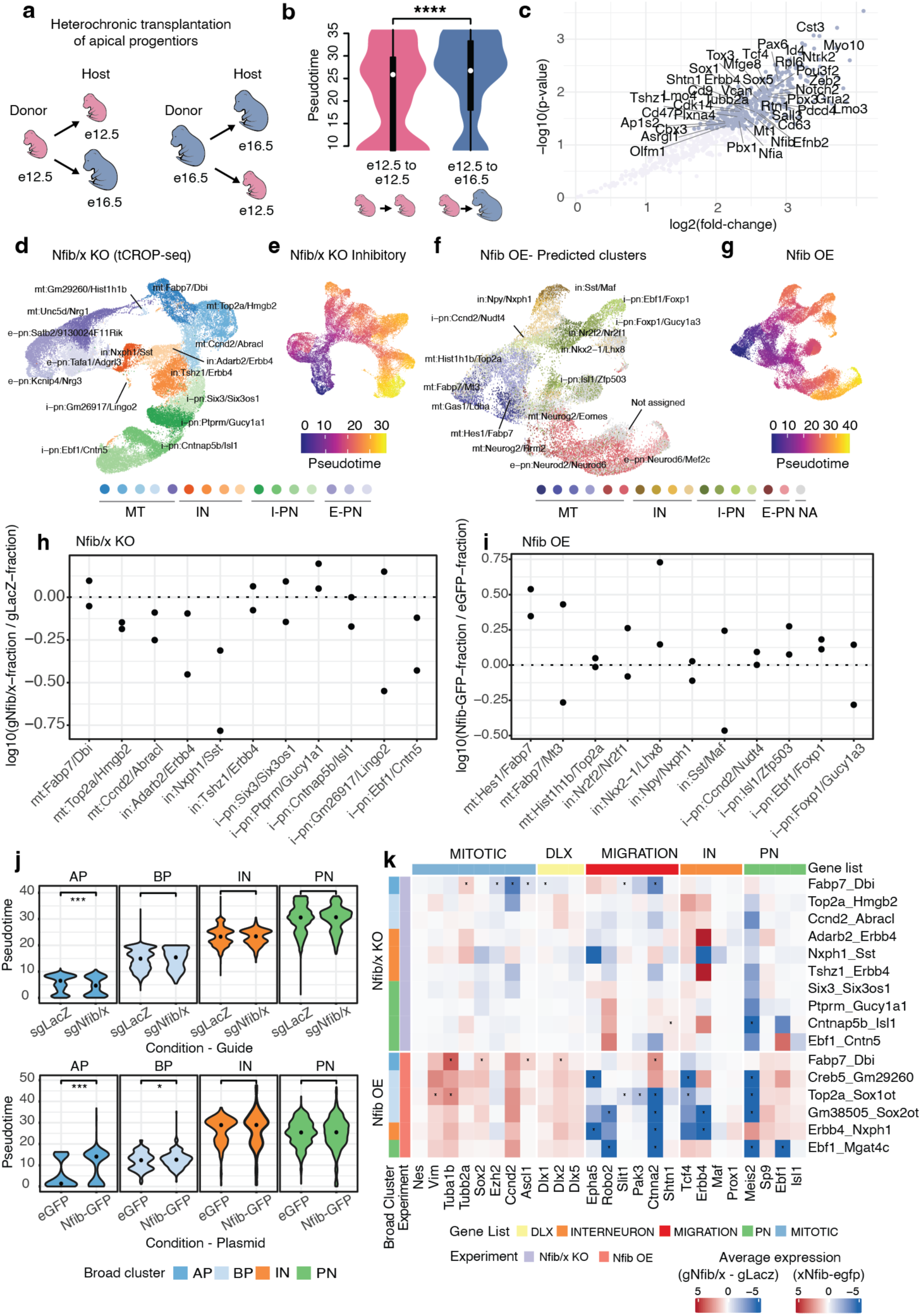
Intrinsic and extrinsic factors regulating progenitor competence. **a**, Schematic overview of donor and host stages for homo- and heterochronic transplantation experiments. **b**, Distribution of transplanted cells along pseudotime in AP_e12.5→e12.5_ and AP_e12.5→e16.5_; two-sided Wilcoxon rank sum test (**** *P*<0.001). **c**, Differentially expressed genes between AP_e12.5→e12.5_ and AP_e12.5→e16.5_; 1<log_2_FC<–1, *P*<0.05. Only genes downstream of Nfib, Meis2 and Tcf4 are labelled. **d**, UMAP-embedding of cells collected in Nfib/x KO. Cells are annotated by broad cell state and the cluster’s top 2 marker genes; mt: mitotic, in: interneuron precursor, i-pn: inhibitory projection neuron precursor, e-pn: excitatory projection neuron precursor. **e**, UMAP-embedding of subsetted inhibitory neuron precursors and their progenitors in Nfib/x KO. Cells are colored by inferred pseudotime scores. **f**, Cells from Nfib OE shown in UMAP-embedding. Cell labels were predicted using label transfer. Cells with low prediction score are labelled as ‘not assigned’ (na). **g**, UMAP-embedding of cells in Nfib OE. Cells are colored by inferred pseudotime scores. **h**, Proportion change per cluster in Nfib/x KO. For each biological replicate, the fraction of cells containing sgNfib/x was compared to the fraction of cells containing sgLacZ. **i**, Proportion change per predicted label in Nfib OE. For each biological replicate, the fraction of cells containing NFIB-GFP plasmid was compared to the fraction of cells containing EGFP control plasmid. **j**, Distribution of pseudotime scores between conditions across broad cell states in Nfib/x KO (upper row) and Nfib OE (bottom row). Dot shows median of corresponding distribution. Two-sided Wilcoxon rank-sum test, *** *P*<0.001, ** *P*<0.01, * *P*<0.05. **k**, Change in gene expression upon perturbation for selected genes. Average gene expression was calculated per cluster and condition. Expression change was calculated by dividing average expression in cells containing sgNfib/x by sgLacZ (for Nfib/x KO) or by dividing cells containing Nfib-GFP plasmid by control plasmid (for Nfib OE). Rows are annotated by broad cell state and experiment, columns are annotated by gene list. Stars indicate differential expression which was inferred using Seurat’s *FindMarker*-function with default parameters; * adjusted p-value < 0.01.

Using clusters from our combined scRNA-seq data as a reference, we applied Bisque (Jew et al., 2020) to estimate the proportions of different neuronal states within the transplantation-derived datasets (Extended Data Fig. 6d,e; Supplementary Fig. 13a). We then assigned a maturation score to each replicate by using the average pseudotime score per reference cluster and weighted it according to the inferred cell state proportions (see Methods). The pseudotime scores were higher when APs were transplanted into an e16.5 environment (AP_e12.5→e16.5_, AP_e16.5→e16.5_) compared to an e12.5 environment (AP_e12.5→e12.5_, AP_e16.5→e12.5_; Fig. 4b, Extended Data Fig. 6f). To identify transcriptomic differences induced by transplantation, we filtered the count matrix by highly variable genes from our combined scRNA-seq datasets and used DeSeq2 (Love et al., 2014) for differential expression analysis (Fig. 4c, Extended Data Fig. 6g). Notably, *Nfib* and many of its downstream genes (among other genes) exhibited increased expression in AP_e12.5→e16.5_ compared to AP_e12.5→e12.5_. We did not observe significantly downregulated genes (Fig. 4c). Furthermore, only two genes downstream of NFIB (*Mlc1* and *Aldoc*) were significantly downregulated in AP_e16.5→e12.5_ compared to AP_e16.5→e16.5_. These findings indicate an involvement of the extrinsic environment in shaping the maturation competence of transplanted cells. The patterns of pseudotime and gene expression were reminiscent of the recipient stage. The gene expression changes observed after transplantation suggest that maturation competence may be more closely associated with the acquisition of specific genes rather than their loss, though this remains to be further explored.

### *Nfib* knockout inhibits while overexpression promotes maturation in inhibitory neurons

To functionally validate the influence of NFIB on maturation competence, we employed two experimental approaches: in vivo CRISPR perturbation using tCROP-seq (Dvoretskova et al., 2024) to knockout *Nfib* and *Nfix* (Nfib/x KO), and overexpression of *Nfib* (Nfib OE). For the tCROP-seq experiment, we performed in-utero electroporation (IUE) at e12.5 to introduce single-guide RNAs (sgRNAs) and Cas9 vectors targeting progenitor cells in the GE of wild-type mouse embryos (C57BL/6). To maximize perturbation efficiency, we employed a combination of sgRNAs targeting both *Nfib* and *Nfix* (sgNfib and sgNfix), as *Nfix* is part of the same downstream transcriptional program through which NFIB coordinates maturation (Matuzelski et al., 2017) (Supplementary Figs. 20 and 11a) and may compensate for *Nfib* loss. This dual-target approach aimed to ensure robust perturbation of the NFIB pathway. Control embryos were targeted with sgRNAs for LacZ (sgLacZ). Cortices, striata, and olfactory bulbs were dissected at e16.5, and cells were enriched by FACS based on TdTomato fluorescence, which labeled sgRNA-expressing cells, and GFP fluorescence, which labeled Cas9-expressing cells (see Methods). To minimize batch effects, we pooled cells from several embryos which received either sgNfib and sgNfix or sgLacZ and then performed multiplexed scRNA-seq (Extended Data Fig. 7a). In total we acquired four replicates for the Nfib/x knockout, consisting of two biological replicates, each with two technical replicates.

The *Nfib* overexpression experiments were conducted in a similar manner by targeting progenitor cells in the GE at e12.5 via IUE. A pCAG vector encoding *Nfib*-GFP was used, along with an additional pCAG vector encoding RFP to facilitate efficient sorting, due to the low GFP signal produced by the *Nfib* overexpression vector. Control embryos were electroporated with the pCAG-eGFP vector. At e14.5, cortices and striata were dissected, and RFP+ cells were enriched by FACS for Nfib OE, while GFP+ cells were used as controls (Extended Data Fig. 7b). We acquired two biological replicates for Nfib OE and control. To confirm the production of functional protein from the exogenous Nfib OE vector, *Nfib* overexpression was performed in Neuro2A cells (see Methods). Detection of NFIB and the HA tag was carried out by western blot using anti-HA and anti-NFIB antibodies (Extended Data Fig. 7c).

The transcriptomic landscape of cells collected from Nfib/x KO and Nfib OE was profiled using scRNA-seq and analyzed using a standard Seurat pipeline (see Methods). For Nfib/x KO, the filtered dataset contained 47,079 cells with 5,887 cells containing sgNfib and/or sgNfix and 30,328 cells containing sgLacZ. Cells were clustered and annotated by their top 2 marker genes (Fig. 4d; Supplementary Fig. 14a). Our dataset contained a fraction of excitatory precursors expressing the marker genes *Neurod2* and *Neurod6* (Supplementary Fig. 17). This likely reflects that targeting GE progenitors via IUE also labels some progenitors of excitatory neurons, presumably located at the interface of ventral and dorsal progenitor domains (Bandler et al., 2022; Dvoretskova et al., 2024). For inferring pseudotime scores we used Monocle 3 on subsetted precursors of inhibitory neurons and their progenitors (see Methods) (Fig. 4e; Extended Data Fig. 7d) (Haghverdi et al., 2018).

Cells from Nfib OE experiments were processed using a workflow similar to Nfib/x KO. To address batch-specific variability, including contributions from ambient RNA observed in one replicate, we excluded cells containing hemoglobin transcripts and performed batch correction using Harmony (Korsunsky et al., 2019). The filtered dataset included 30,019 cells, comprising 5,859 Nfib-GFP^+^ cells and 7,702 eGFP^+^ control cells. We applied label transfer, using the integrated dorso-ventral scRNA-seq dataset as a reference, labelling cells as ‘not assigned’ when their maximum prediction score was below 0.5, thus minimizing the impact of low-confidence assignments on downstream analyses (Fig. 4f; Supplementary Fig. 14b,d). Pseudotime scores were calculated using Monocle3 (Fig. 4g).

We aggregated clusters (in Nfib/x KO) or predicted labels (in Nfib-OE) into broad groups consisting of mitotic cells, INs, PNs, and excitatory precursors (Supplementary Fig. 14c,d), and calculated the proportional changes in these cell states following Nfib/x KO or Nfib OE (see Methods). Across both experiments, the relative fraction of mitotic cells remained stable (Extended Data Fig. 7e,f). However, the overall fraction of post-mitotic inhibitory neurons decreased with Nfib/x KO and increased with Nfib OE (Extended Data Fig. 7e,f).

The decrease in inhibitory neuron precursors following Nfib/x KO was not uniform across finer-grain clusters of INs and PNs, with only some clusters being affected (Fig. 4h,i). To refine our understanding of cell state shifts, we utilized Milo (Dann et al., 2022), a computational tool designed to infer differential abundance within neighborhoods of single cells. Milo identified localized changes in population structure, showing decreased abundances of inhibitory precursors in neighborhoods corresponding to clusters of both INs and PNs (*Adarb2_Npas3*, *Nxph1_Sst*, *Ebf1_Pou3f1*, and *Cntn5_Cdh8*) (Supplementary Fig. 15a,b). This finding was consistent with cell proportion changes observed across clusters (Fig. 4h).

Additionally, we analyzed the effect of the perturbation on postmitotic precursors of excitatory neurons, finding an increased abundance in Nfib/x KO and a decreased abundance in Nfib OE (Extended Data Fig. 7e,f). Changes in abundance were further explored using Milo for Nfib/x KO, with some cell states being more affected than others. Detailed results are provided in the Supplementary Data (Supplementary Fig. 16a-c).

In addition to changes in cell state proportions, we also observed alterations in gene expression and pseudotime trajectories. To assess the transcriptional impact of Nfib/x KO or Nfib OE, we performed DGE analyses between conditions within each cluster (see Methods) and quantified the number of differentially expressed (DE) genes. In Nfib/x KO and Nfib OE, pronounced changes in cell state abundance were not always accompanied by a high number of DE genes. For example, APs in both Nfib/x KO and Nfib OE displayed a relatively high number of DE genes despite minimal changes in cell proportions (Extended Data Fig. 7g,h). Next, we aimed to determine whether the affected genes were direct targets of NFIB. We overlapped DE genes from Nfib/x KO or Nfib OE with genes whose promoters were bound by NFIB in CUT&RUN data (Supplementary Fig. 15c,d). We observed that more than half of the DE genes were directly bound by NFIB (62.9% for Nfib/x KO and 60.1% for Nfib OE). The true proportion of direct NFIB targets is likely higher, as genes regulated via enhancer regions were not considered.

To infer maturation shifts along the pseudotime trajectory following perturbation, we compared pseudotime scores across conditions, with APs showing significantly reduced scores in Nfib/x KO and significantly increased scores in Nfib OE (Wilcoxon rank-sum test) (Fig. 4j). However, this effect did not extend to more mature cell states, as only BPs in the Nfib OE showed a significant increase in pseudotime scores (Fig. 4j).

Next, we focused on genes with various functional roles during neurogenesis and visualized their aggregated expression differences across conditions in each cluster for both the Nfib/x KO and Nfib OE experiments (Fig. 4k). A detailed analysis of gene expression changes, including validation using *in situ* hybridization images from the Allen Brain Institute’s Developing Mouse Brain Atlas (Henry and Hohmann, 2012) and insights into the regulation of cytoskeleton, progenitor markers, migration genes, and markers of post-mitotic cell states, is provided in the Supplementary Data (Supplementary Fig. 19).

Taken together, the shift in pseudotime maturation scores of APs, changes in post-mitotic precursor abundance, and alterations in gene expression underscore NFIB’s regulatory influence. However, not all post-mitotic cell states were equally affected, highlighting a complex, cell state-dependent regulatory landscape.

## Discussion

We describe the regulatory mechanisms that govern progenitor competence during the development of inhibitory neurons. Our results show that the competence of GABAergic progenitors is closely tied to the timing of neurogenesis. This timing primarily influences the maturation of their neuronal progeny, with little impact on their differentiation. Both cell-intrinsic attributes (including TF expression, chromatin remodeling, and reorganization of the gene-regulatory network) as well as cell-extrinsic cues collectively define stage-specific maturation competence. The results suggest a mechanism that may compensate for variations in the time available for migration and network integration between early- and late-born neurons. Data presented in this study are accessible through an interactive online platform, enabling users to explore scRNA-seq, scATAC-seq, and eGRN datasets (http://141.5.108.55:3838/mind_shiny/).

The birthmark of maturation is likely passed from GABAergic mitotic progenitors to their progeny and is primed in chromatin at regulatory regions. In particular, NFIB, a member of the NFI family of TFs, exhibited extensive genomic binding and high regulatory activity at late stages of neurogenesis. NFI TFs are known to regulate both neuronal and glial lineages during central nervous system development (Bunt et al., 2017). Furthermore, they function as cofactors for FOXP2 to facilitate chromatin opening and activate neuronal maturation genes in human subplate and deep layer cortical neurons (Hickey et al., 2019). NFI TFs have been shown to regulate chromatin through various mechanisms, such as binding to nucleosomes (Chávez and Beato, 1997) and chromatin modifiers (Liu et al., 2001), opening chromatin (Adam et al., 2020), controlling chromatin loop boundaries (Pjanic et al., 2013), and by directly altering histone modifications (Pjanic et al., 2013). Furthermore, NFIX has been shown to regulate the timely generation of intermediate progenitor cells from radial glia, partly through the transcriptional upregulation of *Insc* (Harris et al., 2016). In our data, NFIB promotes and forms partnerships with essential regulators of GABAergic IN and PN development, such as TCF4 and MEIS2 (Wang et al., 2022; Su et al., 2022; Dvoretskova et al., 2024), and binds to promoters of the *Dlx* family of genes, known to promote the identity and expansion of GABAergic neurons (Panganiban and Rubenstein, 2002). We propose that NFIB may prime enhancer regions in APs of the GE, initiating chromatin remodeling and leading to stage-specific maturation competence.

We found that the overexpression of *Nfib* in GE progenitors accelerated the acquisition of postmitotic neuronal identity, whereas knockout of *Nfib* and *Nfix* delayed maturation. Although these findings highlight NFIB’s regulatory role, the mechanisms remain unclear. Knockout studies in mice have revealed that deficiency in these genes leads to overlapping brain defects, such as hydrocephalus, corpus callosum abnormalities, and enlarged ventricles (Driller et al., 2007), while neuronal progenitors in the mouse cortex and retina fail to differentiate (Betancourt et al., 2014; Harris et al., 2016; Clark et al., 2019). In humans, haploinsufficiency of *NFI* genes results in overlapping neurodevelopmental phenotypes, including intellectual disability, macrocephaly, and brain anomalies (Zenker et al., 2019a).

The decrease in inhibitory neuron precursors observed after *Nfib/x* knockout was not uniform across all IN and PN branches (Fig. 4h,i; Supplementary Fig. 15a,b). This suggests that NFIs specifically regulate the maturation of certain GABAergic neuron lineages, rather than uniformly affecting all inhibitory neuron subtypes.

Other mechanisms have been proposed to govern neuronal maturation, such as the rate of metabolic activity in mitochondria (Iwata et al., 2023) or selective translation of epigenetic modifiers (Wu et al., 2022). The release of epigenetic barriers sets the timing of maturation in neural progenitor cells, with key factors including EZH2, EHMT1/2, and DOT1L (Ciceri et al., 2024; Appiah et al., 2023). In our study, we observed that *Ezh2*, a member of the polycomb repressive complex 2 (PRC2), is depleted in APs following *Nfib/x* knockout in inhibitory neurons. Interestingly, in *Nfib* knockout mice, *Ezh2* showed upregulated expression within hippocampus and neocortex (Piper et al., 2014). Together, this suggests an interaction between NFIB and members of PRC2, albeit following different regulatory rules in GE and neocortex.

The maturation shift may involve an interplay of extrinsic and intrinsic factors, as GABAergic progenitors in heterochronic transplantation adjust to the host environment by acquiring new gene expression patterns. Potential extrinsic contributors include feedback from newborn cells (Reillo et al., 2017), extracellular vesicle exchange (Pipicelli et al., 2023), and tissue stiffness (Ryu et al., 2021).

While multiple studies described temporal and spatial differentiation patterns in GABAergic neurons (Kelly et al., 2018; Inan et al., 2012; Wonders et al., 2008; Butt et al., 2008; Flames et al., 2007; Fogarty et al., 2007; Miyoshi et al., 2007), there is little evidence of a fate birthmark transmitted from APs to their daughter cells. By contrast, glutamatergic neurons display a birthdate-dependent generation of transcriptomically distinct postmitotic cells that is linked to a progression in the differentiation competence of their progenitors (Di Bella et al., 2021; Telley et al., 2019; Vitali et al., 2018; Yoon et al., 2018). If not through a sequential mechanism, what drives diversity within the GE? Other factors, such as the mode of cell-division (Petros et al., 2015; Kelly et al., 2018), cell-cycle length (Glickstein et al., 2007; Lodato et al., 2011; Zong et al., 2022), progenitor heterogeneity (van Heusden et al., 2021), TFs that transduce patterning signals (Rubenstein and Puelles, 1994; Shimamura et al., 1995; Wichterle et al., 2001; Nery et al., 2002; Xu et al., 2004; Wonders and Anderson, 2006; Flames et al., 2007; Fragkouli et al., 2009; Flandin et al., 2010; Sandberg et al., 2016; Dvoretskova et al., 2024), and differential enhancer activation across spatial regions (Dvoretskova et al., 2024), have been shown to underlay the generation of diverse GABAergic types.

This study contributes to the broader discourse on neuronal maturation, offering insights into the plasticity and commitment of GABAergic progenitors.

## Supporting information

Supplementary Results

## Acknowledgements

We thank members of the Mayer and Myoga laboratories for feedback and discussion; J. Kuhl (somedonkey.com) for illustrations; R. H. Kim from the Max Planck Institute for Biological Intelligence (MPIBI) Next-Generation Sequencing core facility, I. Velasques and G. Eckstein from the Genomics Core facility at the Helmholtz Zentrum Munich (HMGU), M. Spitaler and M. Oster from the MPIB Imaging and FACS core facility, C. Polisseni from the MPIBI Imaging core facility, members of the MPIB/MPIBI animal facility for their technical expertise and the Joyner Laboratory for donating the Nes-FlpoER mouse line. This work was supported by the Max Planck Society, the European Research Council (ERC) under the European Union’s Horizon 2020 Research and Innovation program (ERC-2018-STG, grant agreement no. 803984, GIDE; to C.M.), and the European Commission (SMART GRANT: ERA-NET NEURON, SMART: 01EW1605; to J.W. and M.T.).

## Author contributions statement

A.R.B., Y.K., and C.M. conceived the project and designed the experiments. A.R.B., Y.K., and C.M. led the experimental work. A.R.B., Y.K., and F.N. analysed the data. I.V. and Y.K. performed the TrackerSeq experimental work. E.Dö. performed the RNAscope experiments and imaging, supervised by Y.K. C.P. and C.F. performed and analyzed embryonic electrophysiological experiments. D.R. and M.H.M. performed and analysed the postnatal electrophysiological experiments. E.Dv. and C.M. conducted the tCROPseq experiments. F.N. developed the interactive web platform. A.R.B., Y.K., F.N., and C.M. prepared the manuscript. All authors discussed the results and contributed to the manuscript.

## Extended Data Figures

**Extended Data Fig. 1:**
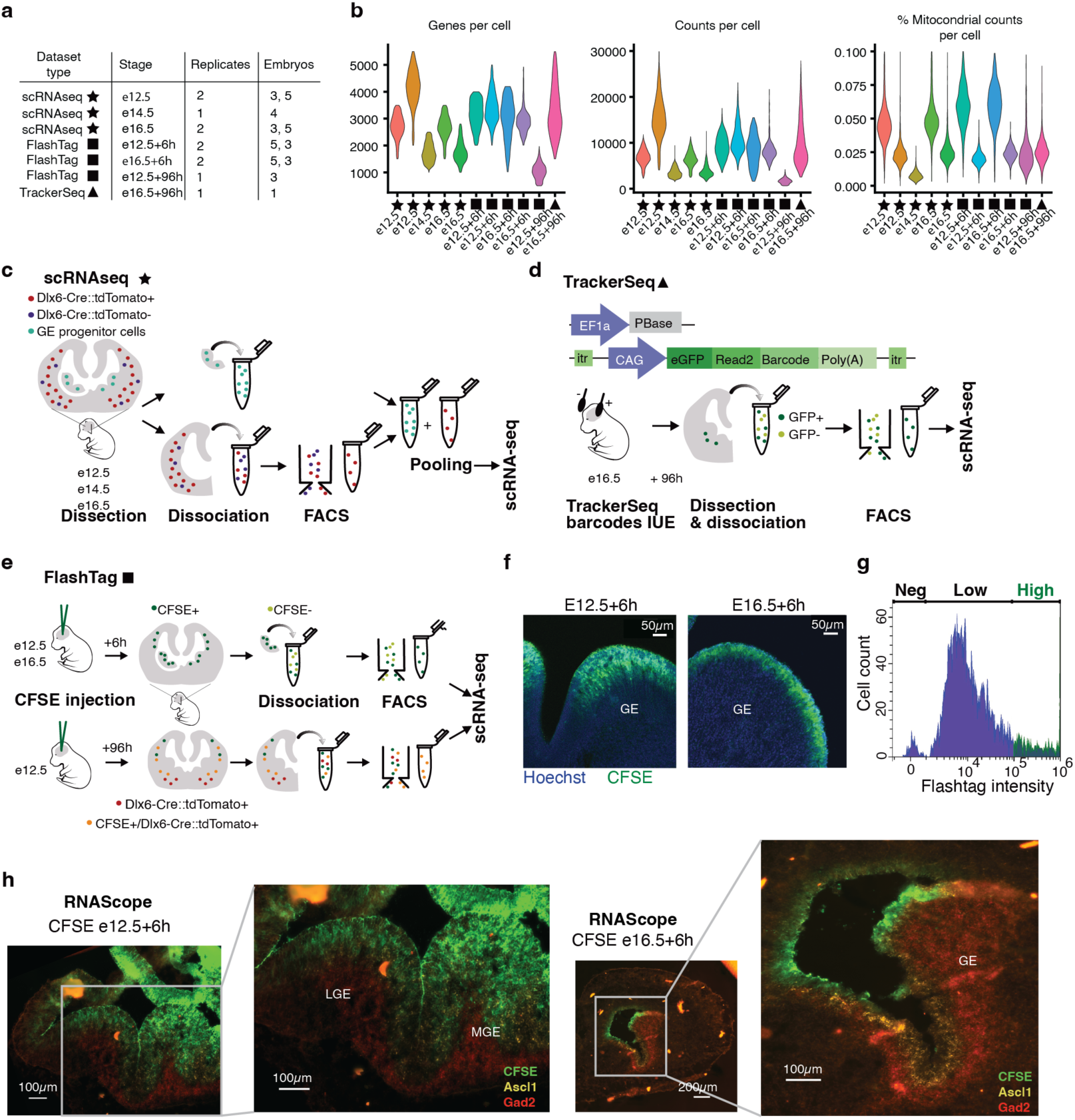
Experimental workflow and dataset characteristics. **a**, Table with dataset type, collection stage, number of replicates, and number of collected embryos per replicate. **b**, Violin plot illustrating the number of genes per cell, counts per cell and mitochondrial gene fraction per cell for each replicate. **c**, Schematic representation of the experimental procedure to generate the scRNAseq datasets. **d**, Schematic representation of the experimental procedure to generate the TrackerSeq datasets. **e**, Schematic representation of the experimental procedure to generate FlashTag datasets. **f**, FT^+^ cells, injected with CFSE. Injection at e12.5 and e16.5, collection after six hours; coronal sections of ganglionic eminences (GE). **g**, FACS plot showing high-intensity FT^+^ cells. **h**, Coronal sections of the GE at e12.5 (left) and e16.5 (right). Cells are labelled with CFSE (in green), RNAscope hybridization probes for Ascl1 (in yellow), and for Gad2 (in red).

**Extended Data Fig. 2:**
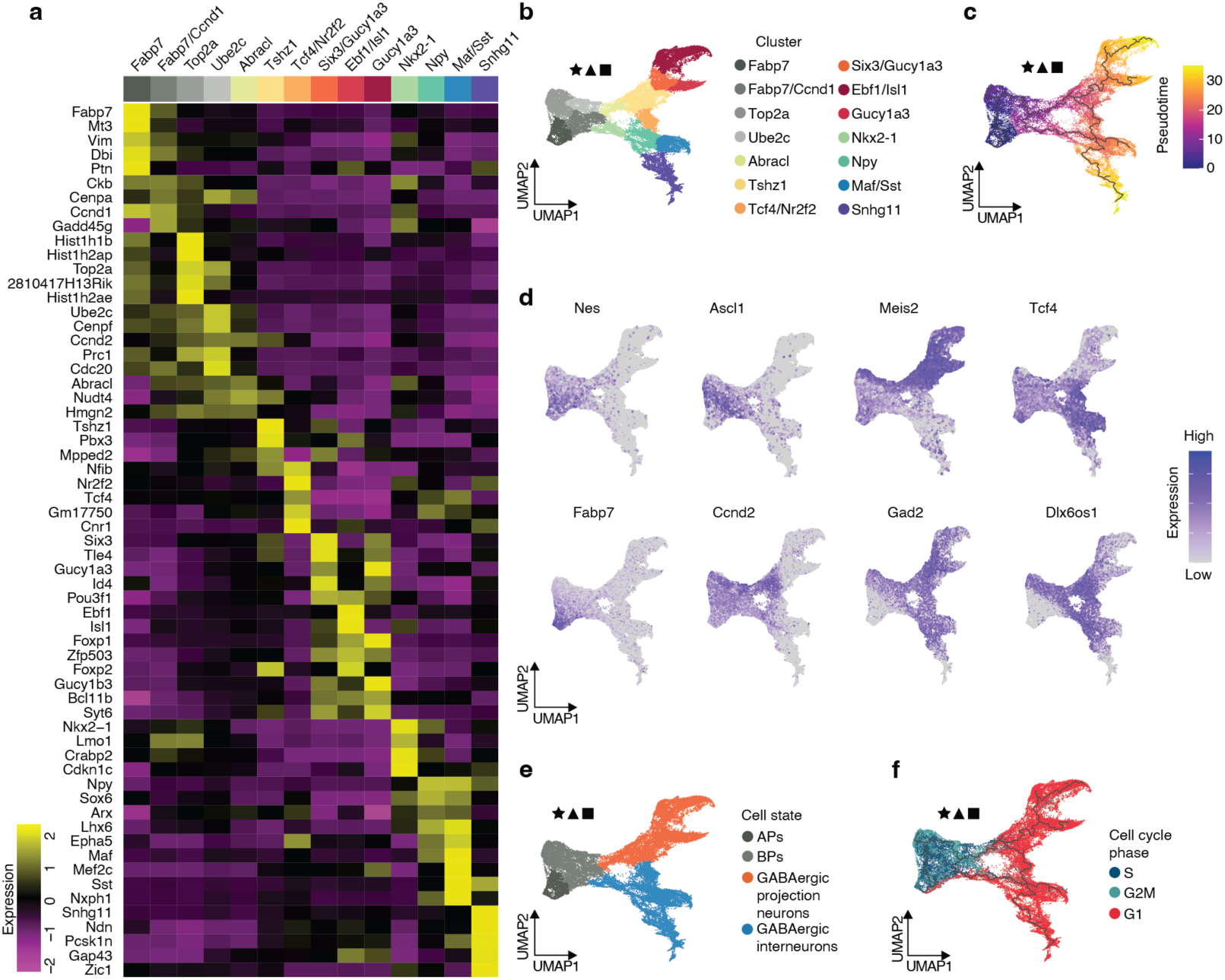
Cell type characterization in the GABAergic lineage. **a**, Heatmap of top five differentially expressed genes in GABAergic cell clusters. **b**, UMAP plot of combined datasets with cells colored by cluster identity. Clusters are annotated by one or two top marker genes. **c**, UMAP plot of combined datasets with inferred Monocle3 trajectory. Cells are colored by pseudotime. **d**, Expression of marker genes in the combined dataset. Nes and Fabp7 label APs, Ascl1 and Ccnd2 are markers for BPs. Post-mitotic inhibitory neurons express Gad2 and Dlx6os1. Meis2 labels PNs and Tcf4 labels INs. **e**, UMAP plot of combined datasets, with cells colored by broad cell states. **f**, UMAP plot of combined datasets colored by cell cycle phases.

**Extended Data Fig. 3:**
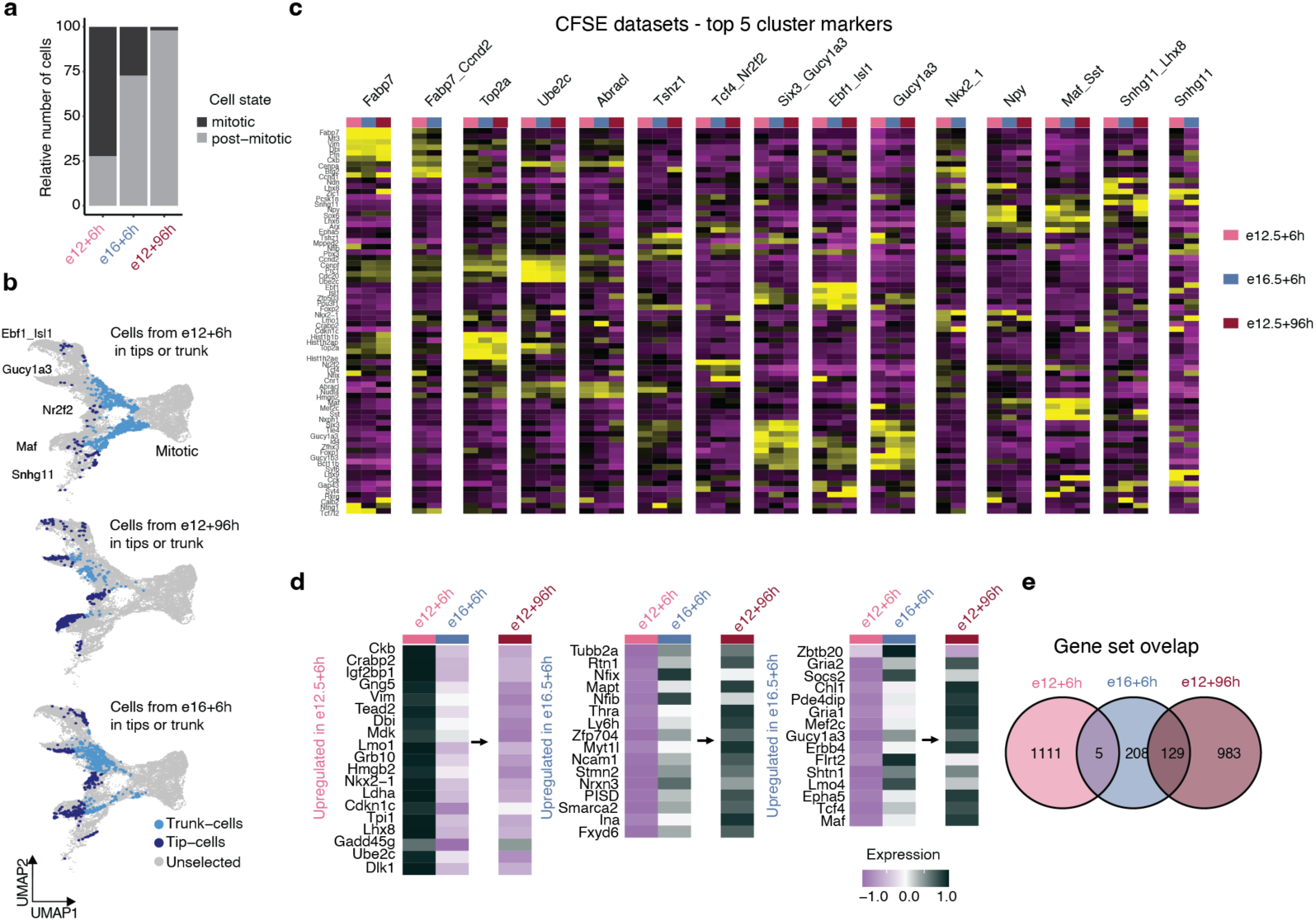
Comparison of early- and late-born cohorts. **a**, Barplot showing the cell ratio in different cell-cycle states for the FT_e12.5 + 6h_, FT_e16.5 + 6h_, and FT_e12.5 + 96h_ datasets. **b**, Post-mitotic cells of FT_e12.5 + 6h_, FT_e16.5 + 6h_, and FT_e12.5 + 96h_ highlighted in the UMAP-embedding of the merged dataset. Color indicates whether cells are part of branch-clusters or in post-mitotic trunk. Comparison of isochronic cohorts displayed side by side shows that cells from the FT_e12.5 + 6h_ cohort predominantly occupy intermediate positions, indicating progression toward the branch tip (Snhg11), whereas cells from FT_e16.5 + 6h_ and FT_e12.5 + 96h_ cohorts have reached the branch tips. **c**, Heatmap of marker gene expression for branch tips split by isochronic cohorts demonstrates that low-abundance states exhibit gene-expression profiles consistent with other cells at the branch tips, supporting their correct classification. **d**, Detailed heatmap of differentially expressed genes between FT_e12.5 + 6h_ and FT_e16.5 + 6h_; 1<log_2_FC<–1, adjusted *P*<0.05. Expression is visualized in FT_e12.5 + 6h_, FT_e16.5 + 6h_ and FT_e12.5 + 96h_ datasets. **e**, Venn diagram showing the intersection of FT_e12.5 + 6h_, FT_e16.5 + 6h_, and FT_e12.5 + 96h_ marker genes; FC > 0.25, pval < 0.05.

**Extended Data Fig. 4:**
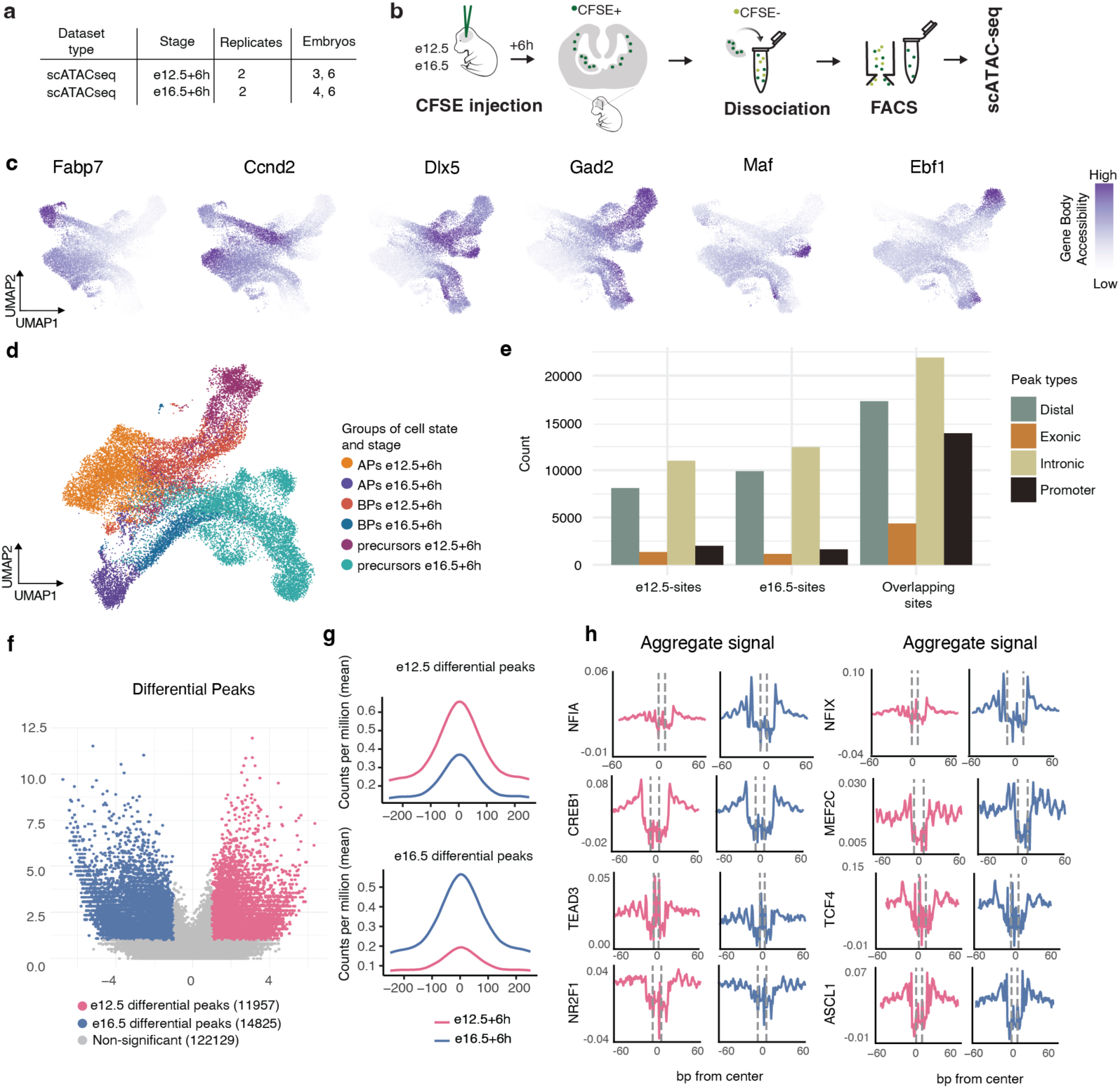
Transcription factor activity and chromatin remodeling in early and late cohorts. **a**, Overview of FT^+^ scATAC-seq datasets: collection stage, number of replicates, and number of embryos per replicate. **b**, Schematic representation of the experimental procedure to generate FT^+^ scATAC-seq datasets. **c**, UMAP depiction of gene body accessibility for marker genes. Fabp7 for APs, Ccnd2 for BPs, Dlx5 and Gad2 for postmitotic inhibitory neurons, Maf for INs and Ebf1 for PNs. **d**, UMAP-embedding of cells in FT^+^ scATAC-seq datasets. Cells are grouped and colored by broad cell state and stage. **e**, Barplot quantifying peak types (distal, exonic, intronic, and promoter) at e12.5, e16.5, and overlapping sites. **f.** Volcano plot displaying differentially accessible peaks between stages, with x-axis showing fold-change and y-axis showing –log10(*p*-value). Significant TFs are highlighted (Pval <= 0.1 and 1<log_2_FC<–1. **g.** Coverage plots of e12.5-enriched sites (top) and e16.5-enriched sites (bottom). Aggregated coverage was calculated for each stage separately. **h**, Aggregate footprint profiles of select transcription factors in FT_e12.5 + 6h_ and FT_e16.5 + 6h_.

**Extended Data Fig. 5:**
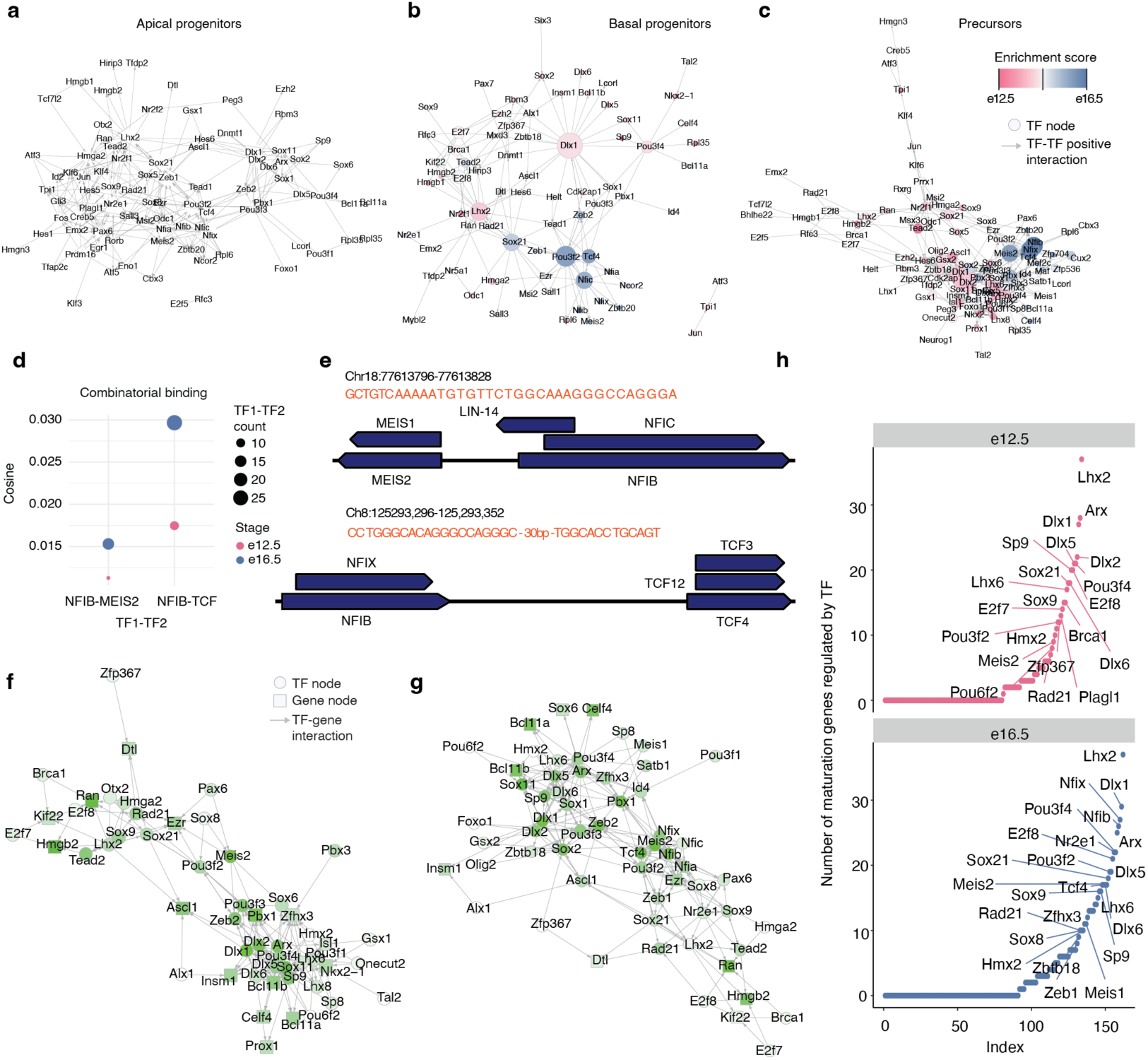
Transcription factor interaction analysis in subnetworks. Subnetwork for APs (**a**), BPs (**b**) and precursors (**c**). Each subnetwork is merged across e12.5 and e16.5, with node color indicating the difference in expression between stages. The size of nodes reflects the number of downstream targets per TF. Subnetworks show only interaction between TFs. **d**, Dot plot showing cosine score on the y-axis and TF pairs on the x-axis. Color and size indicate the stage and the number of occurrences of TF1-TF2, respectively. **e**, Genomic view of transcription factor binding sites for Nfib-Meis2 and Nfib-Tcf4. **f**, TF interaction network of TFs that regulate genes dynamic along the maturation trajectory in e12.5. Nodes are colored by their average expression at e12.5. **g**, TF interaction network showing transcription factors (TFs) regulating genes with dynamic expression along the maturation trajectory at e16.5. Nodes are colored based on their average expression levels at e16.5. **h**, Number of bound genes per TF (out-degree) at e12.5 and e16.5; subsetted for genes dynamic along the maturation trajectory and their upstream TFs. TFs with an out-degree higher or equal than eight are labelled.

**Extended Data Fig. 6:**
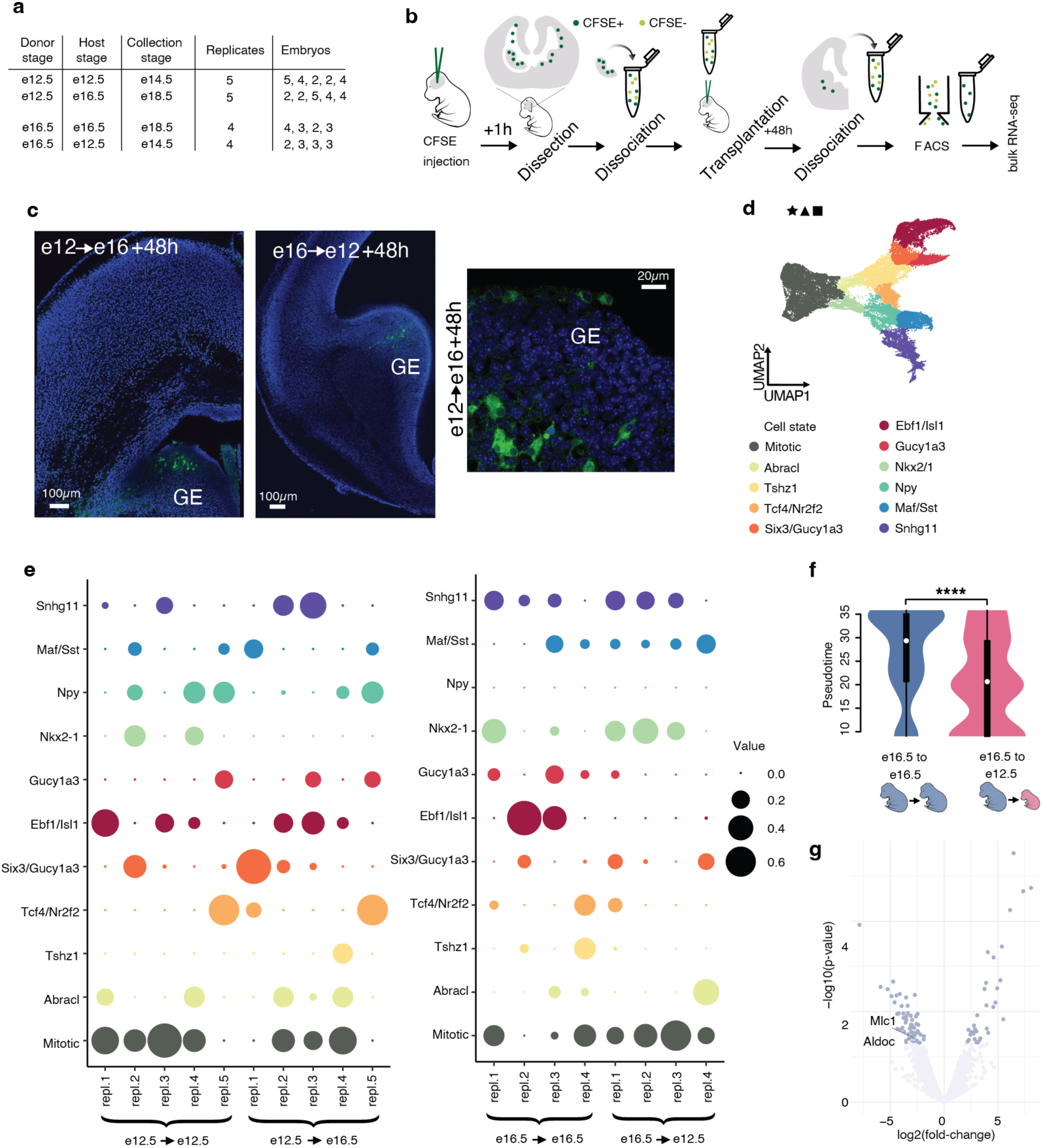
Homo- and hetero-chronic transplantation datasets. **a**, Overview table of homo-and hetero-chronic transplantation experiment datasets: donor, host, and collection stages, number of replicates and number of embryos for each condition. **b**, Schematic representation of the experimental procedure for AP labeling and homo- and heterochronic transplantation. **c**, Images of coronal brain sections after FT^+^ APs transplantation. APs labelled with CFSE; GE, ganglionic eminence. **d**, UMAP plot of the combined dataset utilized for cluster reference. **e**, Predicted cell state composition in each replicate. **f**, Distribution of transplanted cells along pseudotime in AP_e16.5→e16.5_ and AP_e16.5→e12.5_; two-sided Wilcoxon rank sum test (**** adjusted *P*<0.001). **g**, Differentially expressed genes between AP_e16.5→e16.5_ and AP_e16.5→e12.5_; 1<log_2_FC<–1, *P*<0.05. Only genes downstream of Nfib, Meis2 and Tcf4 are labelled.

**Extended Data Fig. 7:**
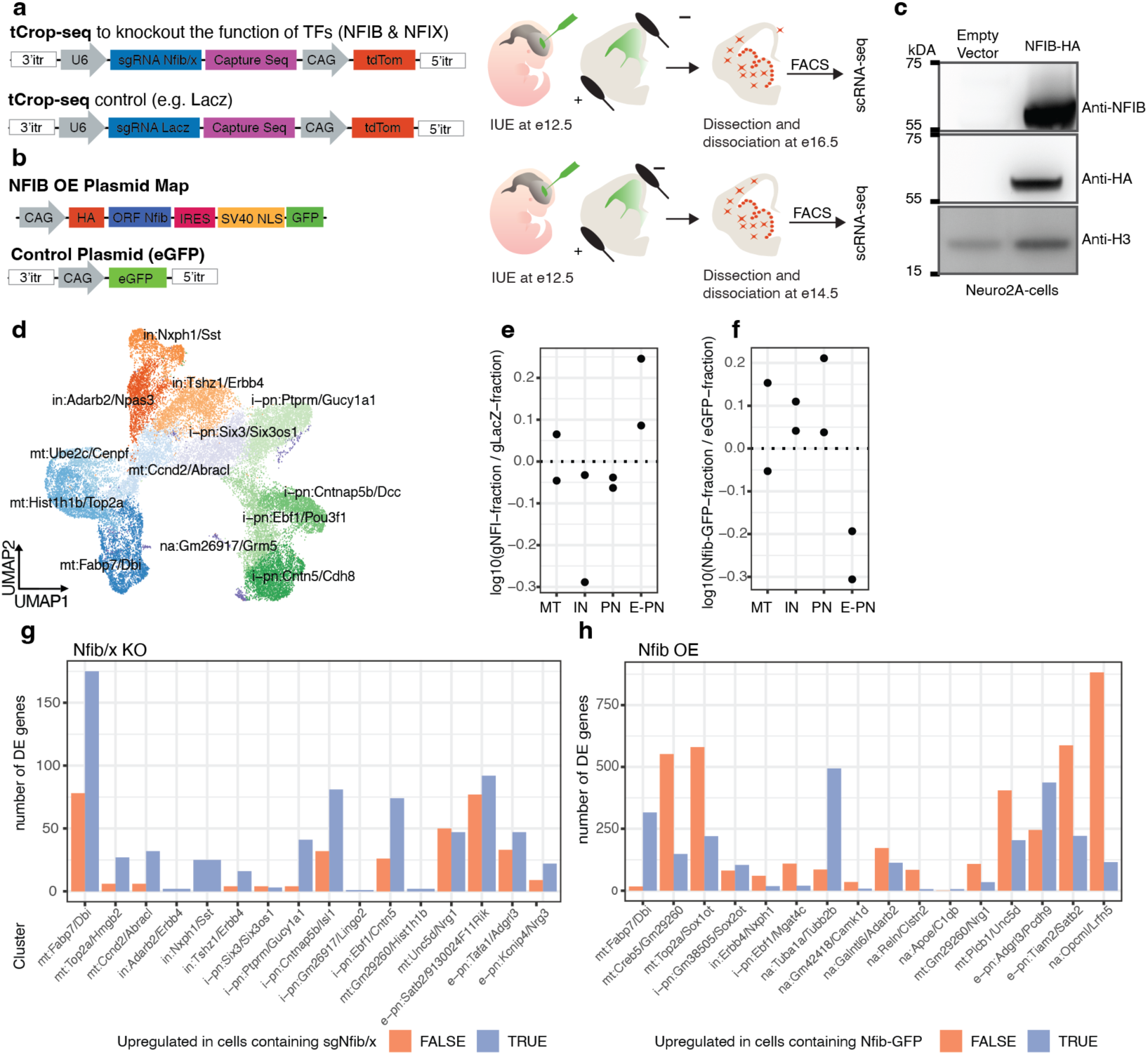
Experimental perturbation of *Nfib* and associated phenotype. **a**, Design of sgRNAs for tCROP-seq experiments and overview of experimental procedure. **b**, Design of plasmids for Nfib OE experiments and overview of experimental procedure. **c**, Western plot showing increased expression of exogenous NFIB in Neuro2A-cells. **d**, UMAP embedding of inhibitory precursors and their progenitors in Nfib/x KO. Cells are colored by clusters, which are annotated based on broad cell states and the top two marker genes; mt: mitotic; in: interneuron precursor; i-pn: inhibitory projection neuron precursor; na: not assigned. **e**, Proportion change in Nfib/x KO. Cells were grouped into broad cell states, by aggregating clusters. **f**, Proportion change in Nfib OE. Cells were grouped into broad cell states by aggregating predicted labels. **g**, Number of DE genes per cluster between conditions in Nfib/x KO. Color indicates positive or negative enrichment. **h**, Number of DE genes per cluster between conditions in Nfib OE. Color indicates positive or negative enrichment.

## Methods

### Animals

All experiments were conducted according to institutional guidelines of the Max Planck Society and the regulations of the local government ethical committee (Beratende Ethikkommission nach §15 Tierschutzgesetz, Regierung von Oberbayern). All mouse colonies were maintained in accordance with protocols approved by the Bavarian government. Mice were group housed in isolated ventilated cages (room temperature 22 ± 1^◦^C, relative humidity 55 ± 5%) under a 12 h dark/light cycle with *ad libitum* access to food and water. Mouse strains used are the following: wild type C57BL/6NRj, Tg(dlx6a-cre)1Mekk (Dlx6-Cre; JAX:008199) (Monory et al., 2006), Rosa26LSL-tdTomato (Ai9; JAX:007909) (Madisen et al., 2010), Tg(Nes-flpo/ERT2)1Alj (Nes-FlpoER; MGI:5532191) (Lao et al., 2012), Gad2<tm1(cre/ERT2)Zjh> (Gad2-CreER, JAX:010702) (Taniguchi et al., 2011), Ai65(RCFL-tdT)-D (Ai65D, JAX:021875) (Madisen et al., 2015). Embryos were staged in days post-coitus, with e0.5 defined as 12:00 of the day a vaginal plug was detected after overnight mating.

### Cell line

Mouse Neuro2a neuroblastoma cells (ECACC, 89121404) were cultured in Dulbecco’s modified Eagle medium (DMEM, Sigma, D6429) supplemented with 10% (v/v) fetal bovine serum (FBS, Sigma, F9665) and containing 1% (v/v) antibiotics (100 U/mL penicillin, 100 mg/mL streptomycin, Sigma, P0781). Neuro2a cells were incubated at 37 ℃ in a 5% CO2 humidified atmosphere and passaged twice a week. Cell passage numbers were limited to no more than 10.

### scRNA-seq (★) datasets: sample and library preparation

Three to six brains from Dlx5/6-Cre::tdTomato mouse embryos were collected at e12.5, e14.5 or e16.5 in ice-cold L-15 medium containing 5% FBS. Ganglionic eminences were manually dissected and dissociated with the Miltenyi Bio Tech Neural Tissue Dissociation Kit (P) (#130-092-628) on a gentleMACS Dissociator according to the manufacturer’s protocol. From the same brains, cortical and striatal regions were dissected, dissociated, and FACS-enriched for tdTomato-positive cells using a SY3200 Cell Sorter (software WinList3D version 8.0.2) or BD FACSAria III Cell Sorter (BD FACSDiva Software, version 8.0.2) with a 100 µm nozzle. TdTomato-positive neurons from the cortex and striatum were pooled with neurons from the GEs, and scRNA-seq was performed. For experiments employing the 10x Genomics platform, Chromium Single Cell 3’ Library & Gel Bead Kit v3 (PN-1000075), Chromium Single Cell 3’ Chip Kit v3 (PN-1000073), and Chromium i7 Multiplex Kit (PN-120262) were used according to the manufacturer’s instructions. Additionally, Chromium Single Cell 3’ Library & Gel Bead Kit v3.1 (PN-1000268), Chromium Single Cell 3’ Chip Kit v3.1 (PN-1000127), and Dual Index Kit TT Set A (PN-1000215) were used according to the manufacturer’s instructions in the Chromium Single Cell 3’ Reagents Kits v3.1 User Guide (Dual Index). Libraries were quantified using a BioAnalyzer (Agilent) and sequenced either on an Illumina NextSeq500 or Novaseq at the Genomics Core Facility of the Helmholtz Center, at the Next Generation Facility of the Max Planck Institute of Biochemistry, or at MLL Münchner Leukämielabor GmbH.

### TrackerSeq (▲) datasets: sample and library preparation

Timed pregnant mice were anesthetized with isoflurane (5% induction, 3% during the surgery) and treated with the analgesic Metamizol (WDT). *In utero* electroporation (IUE) of the TrackerSeq library was performed at e16.5 as previously described in Bandler *et al*. (Bandler et al., 2022). Embryos were injected unilaterally in the lateral ventricle with 700 nL of DNA plasmid solution made of 0.5 µg µL^−1^ pEF1a-pBase (piggyBac-transposase) and the TrackerSeq library 0.5 µg µL^−1^, diluted in endo-free TE bu er and 0.002% Fast Green FCF (Sigma). Embryos were then electroporated with 5 electric pulses (50 V, 50 ms at 1 Hz) with a square-wave electroporator (BTX, ECM 830). The transcriptome libraries were prepared utilizing the 10x Genomics platform as previously described. The lineage barcode library retrieved from RNA was amplified with a standard NEB protocol for Q5 Hot Start High-Fidelity 2X Master Mix (#M094S) in a 50 µL reaction, using 10 µL of cDNA as a template. Specifically, each PCR contained the following: 25 µL Q5 High-fidelity 2X Master Mix, 2.5 µL 10 µmol P7 indexed reverse primer, 2.5 µL 10 µmol i5 indexed forward primer, 10 µL molecular-grade H2O, 10 µL cDNA. The PCR protocol for amplifying TrackerSeq lineage libraries was: (1) 98 °C for 30 s, (2) 98 °C for 10 s, (3) 63 °C for 20 s, (4) 72 °C for 10 s, (5) repeat steps 2–4 for 11 to 18 times, (6) 72 °C for 2 min, and (7) 4 °C hold. Libraries were purified with a dual-sided SPRI selection using Beckman Coulter Agencourt RNAClean XP beads (Beckman Coulter, A63987) and quantified with a BioAnalyzer.

### FlashTag (■) transcriptome datasets: sample and library preparation

Timed pregnant mice were anaesthetised with isoflurane and treated with the analgesic Metamizol as previously described. A CFSE working solution was prepared by adding 8 µL of DMSO and 1 µL of Fast Green to one vial of CellTrace CFSE (CellTraceTM CFSE, Life Technologies, #C34554) for a final concentration of 10 mM, following the instructions from Govindan *et al*. (Govindan et al., 2018). For FT_e12.5 + 6h_ and FT_e16.5 + 6h_, 500 nL of CFSE working solution was injected into ventricles of wild-type C57BL/6NRj embryos at e12.5 and e16.5 respectively. The abdominal wall was then closed, and the embryos were left to develop until collection. After six hours, ganglionic eminences were manually dissected and dissociated on the gentleMACS Dissociator according to the manufacturer’s protocol. FlashTag positive cells with high intensity (> 105) were sorted using FACS (Fig. S1g) and scRNA-seq was performed. For FT_e12.5 + 96h_, 500 nL of CFSE working solution was injected into ventricles of Dlx5/6-Cre::tdTomato embryos at e12.5. After 96 hours, the striatum and cortex were dissected and dissociated on the gentleMACS Dissociator. FlashTag and tdTomato positive cells were sorted using FACS, and scRNA-seq was performed.

### FlashTag (■) chromatin accessibility datasets: sample and library preparation

Sample preparation followed the same protocol described for FT_e12.5 + 6h_ and FT_e16.5 + 6h_ in the previous sections. Single-cell ATAC-seq was performed according to the Chromium Single Cell ATAC Reagent Kits v1 user guide (10x Genomics). FACS sorted cells were centrifuged at 500 rcf for 5 min at 4 °C and resuspended in 100 µL chilled diluted lysis bu er and incubated for 5 min at 4 °C. 1 ml of chilled wash bu er was added to the lysed cells and mixed five times with a pipette, followed by centrifugation at 500 rcf for 5 min at 4°C. The isolated nuclei were counted (using a c-chip hemocytometer) and resuspended in an appropriate volume of chilled diluted nuclei bu er to reach the desired final nuclei concentration. The nuclei were immediately used to generate single-cell ATAC libraries, followed by paired-end sequencing on the Illumina NextSeq 500 platform.

### Electrophysiological analysis of membrane potential in progenitors

After decapitation, the brain was placed in an ice-cold cutting solution saturated with a mixture of 95% O_2_ and 5% CO_2_ containing (in mM): 30 NaCl, 4.5 KCl, 1 MgCl_2_, 26 NaHCO_3_, 1.2 NaH_2_PO_4_, 10 glucose, 194 sucrose. The brain was cut at a thickness of 350 µm on a vibratome (Leica VT1000S, Germany), and the slices were transferred into an artificial cerebrospinal fluid (aCSF) solution containing (in mM): 124 NaCl, 4.5 KCl, 1 MgCl_2_, 26 NaHCO_3_, 1.2 NaH_2_PO_4_, 10 glucose, and 2 CaCl_2_ (310–320 mOsm), saturated with 95% O_2_/5% CO_2_ at approximately 32 °C for 1 hour before being moved to room temperature. Finally, the brain slices were transferred to a recording chamber continuously perfused with aCSF solution saturated with 95% O_2_/5% CO_2_ at 30 °C to 32 °C. Patch pipettes were prepared from filament-containing borosilicate micropipettes (World Precision Instruments) using a P-1000 micropipette puller (Sutter Instruments, Novato, CA), with a resistance of 10 MΩ to 12 MΩ. The intracellular solution contained 130 mM potassium gluconate, 10 mM KCl, 2 mM MgCl_2_, 10 mM HEPES, 2 mM Na-ATP, 0.2 mM Na_2_GTP, pH 7.35, and 290 mOsm. Slices were visualized with a fluorescence microscope equipped with IR–DIC optics (Olympus BX51). Data were obtained using a MultiClamp 700B amplifier, Digidata 1550 digitizer (Molecular Devices), and the software Clampe× 10.3 (Molecular Devices, Sunnyvale, CA). Data were sampled at 10 kHz, filtered at 2 kHz, and analyzed with Clampfit (Molecular Devices). For resting membrane potential recordings, when stable, the membrane potential was recorded for 2 min and the average obtained every 30 s was used.

### RNAscope on FlashTag labelled cells

FlashTag labelling of the cells was performed as for FT_e12.5 + 6h_ and FT_e16.5 + 6h_, by injecting CFSE in the mouse brain ventricles at e12.5 and e16.5 and collecting 6h later. The brains were fixed overnight in 4% PFA solution in 1X PBS at 4 °C. After two washes with 1X PBS, the brains were treated in a series of sucrose solutions (10%, 20%, and 30%) for 12 hours each. The brains were then embedded in OCT. Coronal slices of 10 µm thickness were obtained using a cryostat (Leica CM 3050), placed on Superfrost™ Plus slides, and washed three times with 1X PBS to remove OCT residues. Sample pretreatment and hybridization steps were executed according to the manufacturer protocol (RNAscope® Multiplex Fluorescent Reagent Kit v2 - Cat. No. 323100 from Advanced Cell Diagnostics). Akoya Biosciences Opal fluorophores 570 (1:1500) and 690 (1:5000) and Bio-Techne RNAscope® Probes for Ascl1 (313291) and Gad2 (439371) were utilized for signal detection. The slides were mounted with Prolong Gold Antifade Mountant (P10144 from Invitrogen), stored in the dark at room temperature overnight, and visualized using a Zeiss AxioScan Z.1.

### Transplantation datasets: sample and library preparation

To generate the APe12.5 → e12.5, APe12.5 → e16.5, APe16.5 → e16.5, and APe16.5 → e13.5 datasets, timed pregnant mice were anaesthetised with isoflurane and treated with the analgesic Metamizol as previously described. To target APs, injection of CFSE working solution was performed into wild type C57BL/6NRj embryos at e12.5 and e16.5. One hour later, FT+ APs were collected from three to six embryonic brains. After manual dissection of the ganglionic eminences in ice-cold L-15 medium containing 5% FBS, the tissue was dissociated on a gentleMACS dissociator according to the manufacturer’s protocol. Cells were resuspended in ice-cold HBSS containing 10mmol EGTA and 0.1% Fast Green to a final concentration of 40000 cells/µL to 80000 cells/µL. The cell suspension was split into two separate pools, and 1 µL was injected homo- or hetero-chronically into the ventricles of embryonic brains at e12.5 or e16.5. Forty-eight hours later, ganglionic eminences were dissected and dissociated as described above. CFSE-labelled cells were isolated with flow cytometry and centrifuged 500 rpm, 5min, 4°C. Total RNA-seq libraries were prepared using the SMART-Seq® Stranded Kit (634442, Takara), according to standard manufacturer’s protocol (Low-input Workflow, PCR 1: 5 cycles and PCR 2: 12–15 cycles). The library quality was assessed by using a Qubit™ Flex Fluorometer (Q33327, Thermo Fisher Scientific) and a 4200 TapeStation (G2991BA, Agilent). A total of 10 samples were multiplexed and sequenced in a lane of a NovaSeq 6000 SP flow cell with the 100 cycles kit for paired-end sequencing (2◊ 60 bp) to reduce sequencing batch e ects (100 pM final loading, 42 M reads per sample on average). BCL raw data were converted to FASTQ data and demultiplexed by the bcl2fastq Conversion Software (Illumina).

### Cut&Run sample and library prep

CUT&RUN was performed using the EpiCypher CUTANA CUT&RUN protocol. Two biological replicates were included for each antibody condition: Anti-NFIB, Anti-H3K4me3, and Anti-IgG. Ganglionic eminences were manually dissected from the brains of C57BL/6N mouse embryos collected at e16.5 in ice-cold L-15 medium supplemented with 5% fetal bovine serum (FBS). Tissue dissociation was performed using the Neural Tissue Dissociation Kit (P) (Miltenyi Biotec, #130-092-628) on a gentleMACS Dissociator, following the manufacturer’s protocol. For each sample, 750,000 cells were processed with the following antibodies: Anti-NFIB (Sigma, HPA003956), Anti-H3K4me3 (EpiCypher,#13-0041), and Anti-IgG (EpiCypher, #13-0042), according to manufacturer’s protocols. Library preparation was carried out using the NEB Next Ultra II DNA Library Prep Kit for Illumina (New England Biolabs, #E7645).

### Nfib/x tCROP-seq sample and library preparation

#### gRNA selection and vector construction

The sgRNAs were designed using CRISPick for CRISPRko (Doench et al., 2016; Sanson et al., 2018) and validated with inDelphi (Shen et al., 2018) for high frame shift efficiency. At least 3 sgRNAs per gene were cloned into the backbone using ssDNAs oligo (IDT) and NEBuilder HiFi

DNA Assembly (NEB, E5520). The backbone is a piggyBac plasmid, which encodes TdTomato and sgRNA under the human U6 promoter and has a capture sequence at the sca old of sgRNA for 10x feature barcode retrieval (cs1 incorporated at the 3’ end; (Replogle et al., 2020). The efficiency of the sgRNAs was measured in Neuro2A cells. Cells were transfected with pCAG-Cas9-EGFP (gift of Randy Platt) and sgRNA plasmids using FuGENE 6 Transfection Reagent (Promega, E2691). After 48 h, cells were sorted with Beckman Coulter Cytoflex SRT for TdTomato and EGFP. Genomic DNA was extracted using the Quick-DNA Miniprep Plus Kit (Zymo, D4068), and the region around the sgRNA target was amplified using Q5 polymerase (NEB, M094S) with primers listed in (Table S2), and subsequently sent to Microsynth Seqlab GmbH for Sanger sequencing. Knockout efficiency was quantified using TIDE software (Brinkman et al, Nucl. Acids Res. (2014)). The results for selected sgRNAs are shown in (Table S2).

#### Mice and in utero surgeries

C57BL/6NRj wild-type females (from inhouse breeding) were crossed to wild-type males. Embryos were staged in days post coitus, with E0.5 defined as 12:00 of a day that a vaginal plug was detected after overnight mating. Timed pregnant mice were anesthetized with isoflurane (5% induction, 2.5% during the surgery) and treated with the analgesic Metamizol (WDT). A microsyringe pump (Nanoject III Programmable Nano-liter Injector, DRUM3-000-207) was used to inject 700 nL of DNA plasmid solution made of 0.6 µL of pEF1a-pBase (piggyBac transposase) and pCAG-Cas9-EGFP (both a gift from R. Platt); and the sgRNA plasmid 0.5-8 µL, diluted in sterile 0.9% NaCl solution and 0.002% Fast Green FCF (Sigma, F7252), into the lateral ventricle. Embryos were then electroporated by holding the head between platinum-plated tweezer electrodes (5 mm in diameter, BTX, 45-0489) across the uterine wall, while five electric pulses (35 V, 50 ms at 1 Hz) were delivered with a square-wave electroporator (BTX, ECM830) (Saito, 2006). We used these relatively large electrodes to target all areas of the GE (MGE, CGE and LGE). Before preparing brain tissue for scRNA-seq, each brain was examined under a stereo microscope and only brains that met the following criteria were processed for scRNA-seq: (1) Dispersed tdTomato positive neurons throughout the neocortex. (2) Dense tdTomato positive neurons throughout the striatum. (3) TdTomato positive neurons in the OB

#### Sample collection and sequencing

We collected electroporated brains from mouse embryos at E16.5 in ice-cold Leibovitz’s L-15 Medium (ThermoFisher, 11415064) with 5% FBS (Sigma, F9665). The same media was used during flow cytometry sorting. Papain dissociation system (Wortington, LK003150) was carried out according to the protocol described in Jin *et al*. (Jin et al., 2020) on the gentleMACS™ Octo Dissociator (Miltenyi Biotec). To isolate positive cells for TdTomato and EGFP, flow cytometry was done using a Beckman Coulter Cytoflex SRT with a 100-µm nozzle. After sorting 16,000 individual cells per sample, in PBS (Lonza) with 0.02% BSA (ThermoFisher), were loaded onto a 10X Genomics Chromium platform for Gel Beads-in-emulsion (GEM) and cDNA generation carrying cell- and transcript-specific barcode using the Chromium Single Cell 3’ Reagent Kit v3.1 with Feature Barcoding technology (PN-1000121) following manufacture protocol (document number CG000205, 10X Genomics). We generated 3’ gene expression and sgRNA libraries according to the manufacturer’s manual using the Chromium Library v.3.1 kit (PN-1000121), Feature Barcode Library Kit (PN-1000079) and Single Index Kit (PN-1000213) from 10X Genomics. The quantification of the libraries was performed with the 4200 TapeStation

### Nfib overexpression sample and library preparation

#### Mice and in utero surgeries

Timed pregnant mice were anaesthetised with isoflurane and treated with the analgesic Metamizol as previously described. In utero electroporation was performed at e12.5. Embryos were injected unilaterally in the lateral ventricles with 700 nL of DNA plasmid solution. For the Nfib overexpression (OE) samples, the plasmids used were pCAGG-NFIB2 (Addgene, #112700) and pBCAG-mRFP (Addgene, #40996). The target concentrations for each embryo were 1.5 µg of pCAGG-NFIB2, 1 µg of pBCAG-mRFP, and 0.1% Fast Green to aid injections. For control embryos, the plasmid pBCAG-eGFP (Addgene, #40973) was used at a concentration of 1 µg with 0.1% Fast Green.The abdominal wall was then closed, and the embryos were left to develop until collection.

#### Sample collection and library preparation

At e14.5, electroporated brains were collected in ice-cold Leibovitz’s L-15 Medium with 5% FBS. Cell were dissociated on a gentleMACS Dissociator according to the manufacturer’s protocol. For Nfib overexpression samples, RFP-positive cells were isolated using FACS, while eGFP-positive cells were sorted for control samples. Cells were collected in PBS supplemented with 1% BSA. Libraries were prepared using the Chromium Next GEM Single Cell 3’ Reagent Kit v3.1 (10x Genomics), according to the manufacturer’s instructions. Quality control of the libraries was performed using TapeStation and qubit to ensure proper fragment distribution and concentration. Sequencing was carried out on an Element AVITI sequencer.

### Western Blotting

Neuro2A cells (2◊106 cells/well) were seeded in a 10 cm dishes the day before transfection. The following day, cells were transfected with 8 µg of NFIB-GFP or of empty pcDNA plasmid using Turbofect transfection reagent (R0533, ThermoFisher). Cells were collected 72 hours after transfection by scraping in ice-cold PBS and centrifuging at 400 ◊ g for 5 minutes at 4 °C. Nuclei were extracted by suspending in 2 ml of Bu er A (10 mM HEPES (pH 7.9), 10 mM KCl, 10 mM EDTA, 0.5% Igepal, 1 mM DTT and complete protease inhibitor (Roche, 4693132001) and incubating on ice for 10 min with vortexing at maximum speed every 2 min for 10 s. Nuclei were then collected by centrifugation (800g, 10 min, 4 °C) and the supernatant was carefully removed. Nuclei were disrupted in 0.15 ml Lysis Bu er (50 mM HEPES (pH 7.5), 150 mM NaCl, 5 mM EGTA, 1.5 mM MgCl2, 1% Triton X-100, 1% glycerol and complete protease inhibitor) by shaking at 1500 rmp in Thermomixer for 2 h at 2 °C, with vortexing at maximum speed for 10 s every half an hour. Samples were centrifuged (13000 rmp, 15 min, 4 °C) and the supernatant was collected.

Cell lysates were diluted to 1x in 4x NuPAGE™ LDS Sample Bu er (ThermoFisher, NP0007) with NuPAGE™ Sample Reducing Agent (ThermoFisher, NP0009). Samples were then boiled for 7 minutes at 90°C, 25 µL of each sample was loaded onto NuPAGE™ Bis-Tris Mini Protein Gels,4–12% (NP0322) for electrophoresis and transferred to PVDF membrane (ThermoFisher, PB5210) using a Power-Blotter Semi-dry transfer system (Thermo Fisher Scientific). Membranes were blocked with 5% milk, and then incubated in blocking bu er with rabbit anti-NFIB (1: 1500, Atlas Antibodies, HPA003956), or anti-HA (Proteintech, 51064-2-AP, 1:5,000), and Anti-Histone H3 (1:10000, Sigma,H0164) overnight at 4°C. Proteins were detected using horseradish peroxidase (HRP)-labeled secondary anti-rabbit antibodies (Thermo Scientific, G21234) and developed using SuperSignal West Pico PLUS Chemiluminescent Substrate (Thermo Scientific, 34577).

### scRNA-seq (★), TrackerSeq (▲), and FlashTag (■) transcriptome datasets: pre-processing and merging

Sequencing reads were processed using CellRanger v3.0.2 or v6.1.2 (Zheng et al., 2017), using the mouse reference genome mm10 v2.1.0. Resulting count matrices were analysed using the Seurat package v4.3.0 (Hao et al., 2021) in R v4.1.0. For each dataset, high-quality cells were filtered by the number of genes and mitochondrial read fraction (Extended Data Fig. 1b). Subsequently, counts were normalized and corrected for sequencing depth using Seurat’s *NormalizeData* function. Cell-cycle assignments for each cell were calculated using the cell-cycle gene list from (Tirosh et al., 2016). After identification of highly variable features as described in Butler *et al*. (Butler et al., 2018), we calculated scaled gene-expression values by applying z-normalisation to the 2000 most variable genes, whilst simultaneously regressing out unwanted sources of variation: number of counts per cell, number of genes per cell, mitochondrial read fraction and estimated di erence between cell-cycle phases (*ScaleData* function). The *FindClusters* function with default parameters was used to identify cell clusters. The *FindAllMarkers* function was used to identify cluster marker genes. Clusters with marker genes of excitatory neurons (e.g. Neurod1, Neurod6, Tbr2) or non-neuronal cells (e.g. Apoe, Olig1, Flt1, Pdgfra) were filtered out and excluded from the following steps. Raw counts of samples from scRNA-seq, TrackerSeq, and FlashTag datasets were merged using the Seurat package and aligned using Monocle3 v1.0.0 (Trapnell et al., 2014; Qiu et al., 2017; Cao et al., 2019). For this purpose, the scaled matrix from the Seurat object was converted into a Monocle3 object of the cell_data_set class and preprocessed without the default normalisation, as the dataset was already normalised. Batch-correction was performed using Batchelor v1.8.1 (Haghverdi et al., 2018) followed by Leiden-clustering (using fine resolution) and dimensional reduction using UMAP (McInnes et al., 2020). A developmental trajectory was fitted as a principal graph through fine clusters based on the UMAP-embedding. The root of the trajectory was defined as the cells with the highest *Nes* gene expression, identified in the “Fabp7” cluster. A pseudotime score was assigned to each cell based on its projected position on the trajectory. Leiden clustering (using coarser resolution in Monocle3) identified distinct clusters of cell states. Marker genes specific to each cluster were identified by running di erential expression analysis (in Seurat) using the *FindAllMarkers* function. Clusters were aggregated into broad cell states according to marker gene expression of cell states: Nes and Fabp7 for AP; Ascl1 and Ccnd2 for BP; Tcf4, Lhx6 and Sst for INs; and Meis2, Ebf1 and Isl1 for PNs. Clusters were manually annotated based on broad cell state and marker gene expression. The transition between mitotic and postmitotic cells was defined by selecting the highest pseudotime score of mitotic clusters as the threshold.

To relate post-mitotic precursors to mature cell types in the adult brain, we performed label-transfer on cells in branch tips using scRNA-seq data of GABAergic neuron populations at P10 from Bandler *et al*. (Bandler et al., 2022) as a reference dataset. For label-transfer we used code from Mayer *et al*. (Mayer et al., 2018), which does a correlation-based mapping of cells with the possibility to not assign a label if prediction scores are low.

### scRNA-seq (★) datasets and published datasets: analysis

We downloaded raw counts of e13.5 and e15.5 datasets from Bandler *et al*. (Bandler et al., 2022) (GSE IDs: GSM5684874, GSM5684875, GSM5684876, GSM5684877, GSM5684878, and GSM5684879) and raw counts from the developing mouse somatosensory cortex (Di Bella et al., 2021) at stages e12.5 to e16.5 (GSE IDs: GSM4635073, GSM4635074, GSM4635075, GSM4635076, and GSM4635077). The count matrices were merged with our scRNA-seq (★) datasets to create a combined Seurat object. We filtered cells based on mitochondrial read fraction (≤ 10%). Normalization, scaling, batch correction, dimensionality reduction, and clustering were performed as previously described. Clusters were manually annotated based on top marker gene expression. For cells originating from Di Bella *et al*., we utilized the annotations available from the original paper (Di Bella et al., 2021).

Relative fraction of cells per cell state were calculated for each stage and tissue origin (dorsal vs. ventral) separately, by counting the number of cells per cell state and normalizing by the total number of cells. For cells from the dorsal telencephalon we used annotation from Di Bella *et al*. (Di Bella et al., 2021) for defining cell states. For cells from the ventral telencephalon we used transcriptomic clusters. To ensure that results were not biased by di erent methods for defining cell states, we repeated our analysis based on cell types defined on fine clusters.

We screened for genes that are variable along the pseudotime in inhibitory and excitatory lineages with the following steps: (1) For each lineage, we binned cells from each stage into ten sections based on their inferred pseudotime. (2) Enriched genes were selected based on two criteria: high expression and high gene abundance. High expression was inferred by calculating the fold change between the expression in all cells inside the bin compared to all cells outside the bin. High gene abundance was calculated by comparing the fraction of cells that express a gene inside versus outside the bin (a gene was considered to be expressed in a cell if its scaled expression value was higher than 0.5). (3) A normal distribution was fitted to the changes in expression and abundance. Significantly enriched genes were selected when the di erence in expression and abundance was higher than the corresponding average di erence plus two times the standard deviation of the corresponding fitted distribution. (4) Steps two and three were repeated for each bin. (5) The trajectory of inhibitory neurons diverged as cells leave the cell cycle; therefore, we ran this algorithm for each branch independently and only considered genes that appeared in at least two out of five branches. (6) Finally, by taking the union of dynamic genes of all stages in inhibitory or excitatory lineages, we created a stable set of genes that are dynamic along pseudotime, but conserved across stages.

To compare the correlation of apical progenitors between excitatory and inhibitory datasets, we selected the inhibitory datasets from our study (e12.5, e14.5, e16.5) and the corresponding time points from the excitatory datasets. Apical progenitor cells were then subset from these datasets, followed by normalization and scaling. During this process, we regressed out the e ects of mitochondrial genes, as well as the number of genes and gene counts per cell. Two thousand highly variable genes for apical progenitors were identified using the Seurat function *FindVariableFeatures*. Next, the average expression of the identified genes was calculated for each cluster: excitatory e12.5, e14.5, e16.5 and inhibitory e12.5, e14.5, e16.5. Pearson’s correlation analysis was performed based on these genes. To assess the robustness of the results, we downsampled the datasets to ensure comparable UMI counts across all datasets (*nCount_RNA* < 10,000). Despite this adjustment, similar correlation patterns were observed. Marker genes for excitatory (e12.5, e14.5, and e16.5) and inhibitory (e12.5, e14.5, and e16.5) apical progenitors were identified using the *FindAllMarkers* function in Seurat (min.pct = 0.25, logfc.threshold = 0.25). The intersection between these marker genes and highly variable genes revealed that 30% of the highly variable genes were also marker genes.

### TrackerSeq (▲) datasets: analysis

TrackerSeq barcode reads were pre-processed as described in Bandler *et al*. (Bandler et al., 2022). To assess the clonal coupling between cell states, we calculated z-scores between clusters (Wagner et al., 2018). The z-score is defined as the number of shared barcodes relative to randomized data, with values ranging from positive (coupled clusters) to negative (anticoupled clusters). We utilized these random permutations to calculate empirical *P*-values. For coupled pairs of clusters, the null hypothesis is that the observed coupling is not higher than random coupling. Conversely, for anticoupled pairs, the null hypothesis is that the observed coupling is not lower than random couplings. A random coupling is in contradiction to the null hypothesis when a permutation for one pair of clusters scores above the observed coupling (for positively coupled pairs), or below the observed coupling (for negatively coupled pairs). The relative fraction is reflected in the empirical *P*-value, which was consequently corrected for multiple comparisons using the Benjamini-Hochberg (FDR) method (Benjamini and Hochberg, 1995).

Clones were identified as “dispersing” or “non-dispersing” depending on whether their cells were distributed in multiple or just one branch tip, respectively. We tested whether the transcriptome of mitotic progenitors in “non-dispersing” clones was predictive of their postmitotic state. “Non-dispersing” clones were grouped by their postmitotic cell state in both TrackerSeq_e12.5 + 96h_ and TrackerSeq_e16.5 + 96h_ combined. Separate data frames were created for mitotic and postmitotic subsets of each group. Pearson correlation coefficients were calculated between pairs of gene expression within di erent subsets, generating all the possible combinations of pairs within the columns of the data frames. The subsets of postmitotic clones, mitotic clones, and randomly selected mitotic cells were all correlated to the postmitotic reference group.

### FlashTag (■) transcriptome datasets: analysis

FlashTag datasets were subset from the common trajectory, and di erences in pseudotime between cohorts were assessed using a two-sided Wilcoxon rank-sum test (conf.level = 0.95). Next, di erential gene expression was calculated between the postmitotic fractions of FlashTag_e12.5 + 6h_ and FlashTag_e16.5 + 6h_. The expression of these genes was visualized in a heatmap for FlashTag_e12.5 + 6h_, FlashTag_e16.5 + 6h_ and FlashTag_e12.5 + 96h_.

For the Venn diagram, marker genes for each cohort (FlashTag_e12.5 + 6h_, FlashTag_e16.5 + 6h_ and FlashTag_e12.5 + 96h_) were calculated separately with the *FindMarkers* function in Seurat. Genes with average log2FC > 0.25 and adjusted pval < 0.05 were then intersected, to find common markers across the cohorts.

### FlashTag chromatin datasets: analysis

The raw sequencing data (BCL files) were converted to the fastq format using *Cellranger-atac mkfastq* function from Cell Ranger ATAC v1.2.0 (Satpathy et al., 2019). The reads were aligned to the mm10 (GRCm38) mouse reference genome and fragment files were generated using the *Cellranger-atac count* function. Both time points included 2 replicates and the aligned fragment files were converted to arrow files and analysed further using the ArchR package v1.0.1 (Granja et al., 2021). Dimensionality reduction was performed using latent semantic indexing (LSI), followed by batch correction using Harmony v0.1.1 (Korsunsky et al., 2019). To extract the trajectories of interest and integrate them into an ArchR project, we employed the *getTrajectory* and *addTrajectory* functions, respectively. To visualize enriched motifs, we generated pseudotime heatmaps.

### FlashTag chromatin datasets: temporal dynamics and coverage plots

Peak calling was performed using *addReproduciblePeakSet* function, which runs MACS2 (Zhang et al., 2008) to identify marker peaks for FT_e12.5 + 6h_ and FT_e16.5 + 6h_ datasets. Peaks were classified by either identifying for peaks that overlap across stages, based on genomic position (e12.5-enriched, e16.5-enriched or non-enriched peaks); or by conducting di erential peak analysis with p-value cuto *p* = 0.05 using ArchR *getMarkerFeatures* function. As an additional quality check we counted the number of reads that map to peak regions and calculated their fraction in respect to all reads (Supplementary Fig. 9e). For both scATAC-seq and H3K4me3 ChIP-seq datasets, we calculated peak coverage for each peak category using the *ScoreMatrixList* function.

### FlashTag chromatin datasets: transcription factor footprint analysis

Footprint analysis was carried out on FT_e12.5 + 6h_ and FT_e16.5 + 6h_ datasets using transcription factor occupancy prediction tool: TOBIAS v0.14.0 (Bentsen et al., 2020). We employed the Jaspar non-redundant motif database (Castro-Mondragon et al., 2022) as the primary reference source for motif data. Bias correction was performed to generate corrected bigwig files using the *ATACorrect* function with default parameters. Footprint scores were calculated on corrected bigwig files using the *FootprintScores* function, and di erential binding TFs were detected using *BINDetect* function. Predicted TFs were categorised as significant based on two criteria: their di erential binding score (greater than 0.2 for e12.5 and less than –0.4 for e16.5:referred to as change) and the –log10 of the p-value from the statistical test against a background model. The footprints were visualized using the *PlotAggregate* and *PlotHeatmap* functions.

### FlashTag chromatin datasets: co-binding analysis

To detect co-occurring TF binding sites, we utilized TF-COMB v1.1 (Bentsen et al., 2022). A distinct CombObj was created by loading unique peak sets (e12.5 and e16.5) identified previously. Transcription factor binding sites were identified within the peak regions followed by market basket analysis. TFs co-occurring with NFIB were then subsetted and further assessed for their co-binding (cosine score) and binding events via dot plot.

### FlashTag transcriptome and chromatin datasets: gene regulatory network prediction

We used Scenic+ (v0.1) (Gonzalez-Blas et al., 2023) to predict GRNs for CFSE-labelled cells at e12.5 and e16.5. As scRNA-seq and scATAC-seq data were unpaired we created a common annotation, by defining broad cell states (AP, BP, and precursor) in the transcriptomic data, merging clusters based on marker gene expression. Annotations in the scATAC-seq data were created by applying label transfer, based on gene-scores predicted by ArchR. These broad cell states were split by stage, resulting in 6 stage-specific cell states (Supplementary Fig. 10a). Scenic+ performs co-accessibility analysis of regions and links regions to upstream TFs by searching for enriched TF motifs in regions. To make this analysis more coherent with prior results, we used the previously calculated peak-set from ArchR as input for Scenicplus, instead of recalculating a new peak-set using pycisTopic (Bravo González-Blas et al., 2019). Following the Scenic+ workflow, we created topics of co-accessible regions and performed binarization and motif enrichment of regions in the 20 most important topics. Networks were created by aggregating 10 cells from both modalities of corresponding cell states into pseudocells and then inferring TFs and regions that are predictive of a gene, based on co-accessibility and motif enrichment. The regions considered for a gene have to lie within a genomic interval of 150 kb up- and downstream of the gene. The results are so-called “eRegulons”, i.e. regulatory triplets of one TF, bound regions and corresponding target genes. For each eRegulon the activity in each cell was calculated using AUC-scores (Aibar et al., 2017). Each eRegulon was filtered using standard filtering (*apply_std_filtering_to_eRegulons* function) and high quality eRegulons were selected by filtering for eRegulons where TF-expression and AUC scores correlated more than 0.5 or less than −0.5.

We reconstructed cell-state specific subnetworks by running AUC-binarization (*binarize_AUC*-function). Here, we filtered for (1) eRegulons that are active in at least 50% of cells within a corresponding cell state and for (2) corresponding target genes that have a higher normalized expression than 0.5 (normalized expression is log1p transformed after correcting for sequencing depth). Cell state specific networks (APs, BPs and precursors) were created by merging the corresponding e12.5- and e16.5-subnetwork using the igraph library v1.5.0 (Csardi and Nepusz, 2006). Stage specific networks (e12.5 and e16.5) are similarly created by merging subnetworks of APs, BPs, and precursors of the same stage. In both approaches, the merged networks consisted of the union of vertices and edges. GO-enrichment analysis of target genes was performed using DAVID with default parameters (Dennis et al., 2003).

### CUT&RUN preprocessing and analysis

Raw sequencing reads were mapped to the Mus musculus reference genome (mm10) using Bowtie2 (Langmead and Salzberg, 2012) using parameters –end-to-end –very-sensitive –no-mixed –no-discordant –phred33 -I 10 -X 700 for mapping of inserts 10-700 bp in length. Reads were also aligned to the spike-in genome. Following alignment, duplicate were marked using Picard (https://broadinstitute.github.io/picard/). Peaks were called using MACS2 (p-value cuto of 1 * 10^−4^) using IgG as background. Signal tracks were then generated in bigWig format for visualization in genome browsers. Peak heatmaps and genome browser profiles were generated by using flu heatmap and flu profile function (Georgiou and van Heeringen, 2016). Enriched motifs were identified using findMotifsGenome.pl of HOMER (Heinz et al., 2010). Motif heatmap was generated using visualization package (Zhang, 2024). Peak heatmap signal quantification was done using normalized read counts(RPKM) by averaging read coverage from peak summit +/− 500bp.

### Transplantation datasets: analysis

We used the Galaxy web platform on the public server at usegalaxy.eu to analyse the data (Afgan et al., 2018). Paired end reads were trimmed with the Trimmomatic tool and quality control was performed with FastQC. Reads were mapped to the mouse reference genome using the HISAT2 algorithm (Kim et al., 2019) and the number of reads per annotated genes was counted using featureCounts (Liao et al., 2014). After Fragments Per Kilobase of transcript per Million mapped reads (FPKM) normalization of the count matrices, the proportion of single cell states within each replicate was inferred with Bisque v1.0.5 (Jew et al., 2020) by using the annotated combined single-cell clusters as reference. A weighted pseudotime score was assigned to each replicate by calculating the median of the pseudotime score per cluster from the combined single cell datasets. For di erential gene expression analysis, the count matrices were subset by variable genes of inhibitory neuron datasets and DESeq2 v1.42.0 was utilized (Love et al., 2014).

### tCROP datasets: analysis

Reads from transcriptome and guide libraries of all 4 replicates were mapped to the mm10 reference genome and demultiplexed using C Cellranger (v8.0.1) (Zheng et al., 2017). Single-cell count matrices from transcriptomic libraries of the 4 replicates were merged in Seurat (Hao et al., 2021). We excluded cells with more than 10% fraction of mitochondrial reads, cells predicted to be doublets according to DoubletFinder (McGinnis et al., 2019) and cells that contained both sgNfib/ sgNfix and sgLacZ. After cleaning the dataset, count data was log-normalized and variable features were calculated using Seurat’s *FindVariableFeatures* function. Log-normalized expression data of variable genes was scaled using Seurat’s *ScaleData* function, while regressing out e ects of read-depth (number of genes and number of UMIs) and fraction of mitochondrial reads. Based on the scaled expression matrix, we calculated low-dimensional representations of cells (PCA and UMAP). Cells were clustered using *FindNeighbors* and *FindClusters* functions with default parameters. Clusters were annotated by calculating marker genes for each cluster, using *FindAllMarkers* function, and naming clusters by their top 2 positively enriched marker genes.

By counting the number of cells that contain either sgNfib and/or sgNfix or contain sgLacZ in each cluster per biological replicate, we calculated proportion changes induced by Nfib/x knockout in each cluster. Proportion change was calculated by dividing the number of cells that contained gNfib/x per cluster by the number of cells that contained gLacZ per cluster and applying *log*10 transformation to the result of the division. This was repeated for broad cell states, which were generated by aggregating individual clusters. To infer the e ect of perturbation on gene expression we performed DE-analysis between cells containing gNfib/x and cells containing gLacZ in each cluster using Seurat’s *FindMarkers* function with default parameters. The number of di erentially expressed genes per cluster was inferred by setting a cut-o of the adjusted p-value being smaller than 0.01.

Due to spread of CRISPR constructs during IUE in the ventricle, progenitors of excitatory neurons that lie dorsal of GEs were also targeted. Based on expression of marker genes for excitatory precursors (*Eomes*, *Neurod2*, *Neurod6*) and transcriptomic clustering, we filtered the dataset for inhibitory precursors and their progenitors, by removing cells belonging to 5 clusters (*Gm29260_Hist1h1b*, *Unc5d_Nrg1*, *Satb2_9130024F11Rik*, *Tafa1_Adgrl3* and *Kcnip4_Nrg3*). Processing of the inhibitory subset, from inferring variable features to clustering, was repeated in the same way as described above. We inferred pseudotime scores for the inhibitory subset by converting scaled data and UMAP representation into a cell-data-set object. This was used to infer a trajectory of single cells and infer pseudotime scores using Monocle3 (Haghverdi et al., 2018). We ran Milo on both inhibitory and excitatory subsets after excluding cells that did not contain any guide RNA. Seurat preprocessing was repeated and Milo was executed by setting *k* = 40 for both subsets of data.

### Nfib OE datasets: analysis

For mapping reads from Nfib OE experiments we added the sequence of Nfib-GFP, eGFP and RFP to the mm10 reference genome, using Cellranger’s *mkref* function. Subsequently, mapping and demultiplexing were performed for all four experiments using Cellranger (v8.0.1) with the custom reference genome. Single-cell count matrices from all 4 experiments were merged in Seurat. We detected high levels of ambient RNA (as indicated by ambient expression of hemoglobin genes). Therefore, cells that expressed both *Hbb-bt* and *Hbb-bs* were removed from further analysis. Additionally we also removed cells with more than 20% of mitochondrial reads and cells that were predicted to be doublets according to DoubletFinder (McGinnis et al., 2019). As described above, read counts were normalized, variable features were inferred, data was scaled and we performed PCA. To account for di erences in data quality between experiments, we performed batch-correction using Harmony (Korsunsky et al., 2019). Based on the Harmony-corrected data we inferred clusters and UMAP-embedding. Clusters were annotated by their top 2 marker genes. Assigning clusters to cell states was less straightforward in this dataset, as some clusters contained marker gene expression for multiple cell states. In order to circumvent this problem, we ran label transfer using our integrated dorsal-ventral scRNA-seq dataset as a reference. Cells with a low prediction score (*<* 0.5) were labelled as ‘not assigned’.

Proportion changes of predicted cell states upon over-expression of Nfib, were calculated by comparing the number of cells that express Nfib-GFP (and not eGFP) to the number of cells that express only eGFP per predicted cell state. This was done twice, once for predicted clusters and once for aggregated cell states following the same rationale as for Nfib/x KO. Di erentially expressed genes across conditions were inferred by running Seurat’s *FindMarkers* function for each predicted cluster. Pseudotime scores were inferred in the same way as for tCROP experiments.

### Shiny-based webserver

Results from scRNA-seq experiments (dorsal and ventral wild type scRNA-seq datasets with CFSE and lineage tracing datasets), together with results from scATAC-seq, NFIB CUT&RUN and eGRN analysis were made publicly available via Shiny-based webserver (Chang et al., 2024).

