## Supplementary Results for "NFIB influences progenitor competence in maturation of GABAergic neurons in mice"

#### Electrophysiological and morphological characterisation of temporal cohorts at postnatal stages

To explore the effects of different maturation dynamics of early- and late-born cohorts on postnatal neurons we developed a genetic mouse strategy for fluorescent labelling of temporal cohorts, enabling a basic characterisation of the electrophysiological properties of isochronic cohorts in postnatal brain slices. We generated a triple mutant mouse line containing an Ai65D intersectional reporter allele, a NesFlpoER allele and a Gad2CreER allele. In this line, administration of tamoxifen (TX) induced the expression of tdTomato in a cohort of cells that was transitioning from a mitotic (Nes) to a post-mitotic (Gad2) cell state at the time of TX administration (i.e. $TX_{e12.5}$ and $TX_{e16.5}$; Figure S21a-d). Given the broad diversity of GE-derived GABAergic neurons in the telencephalon, we decided to focus on the striatum.

Following embryonic administration of TX, we prepared brain slices at P8, a developmental stage at which mature spines are known to emerge in the striatum (Lee and Sawatari, 2011), and recorded the membrane response to hyperpolarising and depolarising current steps (Figure S22a). The maximum action potential firing rate, the resting potential and the membrane capacitance were similar between $TX_{e12.5}$ and $TX_{e16.5}$ cohorts (Figure S22e,f,g). Rheobase currents have been shown to increase (Peixoto et al., 2016) and membrane resistance to decrease (Dehorter et al., 2011) during early striatum development. In line with this, the $TX_{e12.5}$ cohort showed higher rheobase currents (Figure S22b) and lower input and membrane resistances (Figure S22c,d) than the $TX_{e16.5}$ cohort. The relationship between injected current and spiking frequency (I-F curve) was shifted rightward in $TX_{e12.5}$ cohorts (Figure S22h) and the membrane responses to current injection (I-V curves) showed decreased potential responses to hyperpolarizing or depolarizing current injections in respect to the $TX_{e16.5}$ cohort (Figure S22i). Previous research showed a decrease in the number of proximal dendrite intersections during postnatal striatum development (Krajeski et al., 2019). Supporting this, a Sholl analysis on biocytin-reconstructed striatal cells revealed that the $TX_{e12.5}$ cohort exhibited shorter total dendritic lengths (Figure S22j) and simpler dendritic arborization overall (Figure S22k). Taken together, early-born cohorts at P8 display advanced physiological and morphological properties, despite observed differences in maturation dynamics at embryonic stages.

##### Tamoxifen induction

Tamoxifen was dissolved in corn oil to generate a concentration of 20 $\mathrm{mg\,mL^{-1}}$ by constant stirring at 37°C. Mouse strains Nes-FlpoER$^{+/-}$, Gad2-CreER$^{+/-}$ and Ai65D$^{+/+}$ were crossed to obtain Nes-FlpoER::Gad2-CreER::Ai65D embryos. At e12.5 and e16.5 100 $\mathrm{mg\,kg^{-1}}$ of tamoxifen was administered to the pregnant dam by gavage. The embryos were allowed to develop until collection at P8.

##### Whole-cell patch-clamp recordings

Acute striatal slices (300 µm thick) were obtained from triple transgenic mice at P8 of pregnant mice after tamoxifen gavage at e12.5 and e16.5. Mice were decapitated, and brains were rapidly removed and placed in a carbogenated (95% $O_2$/5% $CO_2$) N-methyl-D-glucamine-based artificial cerebrospinal fluid (NMDG-aCSF) solution containing the following (in $\mathrm{mmol\,L^{-1}}$): 92 NMDG, 2.5 KCl, 1.2 $NaH_2PO_4$, 30 $NaHCO_3$, 20 HEPES, 25 glucose, 5 sodium ascorbate, 2 thiourea, 3 sodium pyruvate, 10 $MgSO_4{\cdot}7H_2O$, and 0.5 $CaCl_2{\cdot}2H_2O$ (pH 7.2–7.4, 300–310 mOsm). Slices were cut using a VT1000S vibratome (Leica Microsystems) and then incubated for 11 min at 32–34 $^\circ\mathrm{C}$ with carbogenated NMDG-aCSF solution, followed by at least one-hour recovery at RT in an aCSF solution containing the following (in $\mathrm{mmol\,L^{-1}}$): 119 NaCl, 2.5 KCl, 1.2 $NaH_2PO_4$, 24 $NaHCO_3$, 12.5 glucose, 2 $MgSO_4{\cdot}7H_2O$, and 2 $CaCl_2{\cdot}$2H2O (pH 7.2–7.4, 300–310 mOsm).

Recording pipettes were pulled from BF150-110-7.5HP borosilicate glass capillaries (Science Products) on a P97 horizontal puller (Sutter Instruments) with a typical resistance of 2–5 $\mathrm{M\Omega}$. Pipettes were filled with K-gluconate internal solution containing the following (in $\mathrm{mmol\,L^{-1}}$): 135 K-gluconate, 4 KCl, 2 NaCl, 10 HEPES, 0.2 EGTA, 4 MgATP, 0.5 NaGTP, and 10 phosphocreatine (adjusted to pH 7.3 with KOH and 290 mOsm with sucrose). Biocytin (0.5%) was added to the electrode solution, and the pipettes were backfilled to minimize background fluorescence.

Whole-cell patch-clamp recordings were performed after seal rupture under a SliceScope upright microscope (Scientifica) equipped with epifluorescence and infrared differential interference contrast (IR-DIC) optics. All recordings were performed at RT (22–25 $^\circ\mathrm{C}$) from fluorescently labelled neurons located in the dorsal striatum. Intrinsic properties were measured using the voltage response to a series of hyperpolarizing and depolarizing current injections ($10\,\mathrm{pA}$ steps, starting at $-150\,\mathrm{pA}$). Rheobase current was defined as the smallest positive current step capable of eliciting at least one action potential. Input resistance was calculated with a $-150\,\mathrm{pA}$ hyperpolarizing step from the resting membrane potential and from a linear fit to a voltage-current plot. Whole-cell capacitance was measured using a $5\,\mathrm{mV}$, $200\,\mathrm{ms}$ step from $-90\,\mathrm{mV}$, with a 4-pole Bessel filter with a cutoff frequency of $10\,\mathrm{kHz}$. A bi-exponential fit of membrane response following the $5\,\mathrm{mV}$ depolarization was used to calculate the weighted tau and consequent whole-cell capacitance as described by Gertler *et al.* (Gertler et al., 2008).

Data were acquired using a MultiClamp 700B amplifier and digitized using a Digidata 1550B acquisition system (Molecular Devices). Signals were further filtered at $2\,\mathrm{kHz}$ and digitized at $10\,\mathrm{kHz}$. For current-clamp recordings, the bridge balance was adjusted and pipette capacitance was neutralized. pClamp 11 software was used for analysis (Clampfit, Axon Instruments).

##### Dendritic reconstruction

For anatomical reconstruction, brain slices containing biocytin-filled neurons were post-fixed in 4% paraformaldehyde for 12 h at 4 °C. After fixation, slices were washed 3 times in phosphate buffer saline (PBS, pH 7.5) and permeabilized with PBS containing 0.2% Triton-X100 for 24 h at 4 °C. Following 5 washes for 10 min in PBS, biocytin localization was visualized by incubating the slices in Alexa 488-coupled streptavidin (diluted 1:1000 in PBS, Molecular Probes) for 48 h at 4 °C. Slices were washed a further 5 times in PBS and mounted on Superfrost slices (ThermoFisher Scientific). Z-stack serial images were acquired on a SP8 Leica confocal microscope (Leica Microsystems) using a 40× oil immersion objective. Neuronal arbour reconstruction and analysis were carried out using the Simple Neurite Tracer (SNT) plugin in the Fiji ImageJ package (https://imagej.net/Fiji). Total neurite arbour size and branching were measured within Fiji ImageJ. The Sholl analysis was performed using the skeletonized neuron from SNT, with the centre of the cell body as the origin. No correction was applied for tissue shrinkage during fixation.

#### Gene Expression Dynamics and Functional Insights in *Nfib/x* KO and *Nfib* OE

We analyzed genes with various functional roles during neurogenesis to investigate their expression dynamics across conditions in each cluster for both the *Nfib/x* KO and *Nfib* OE experiments. Aggregated expression differences were visualized (Supplementary Fig. 19), and validation was performed using *in situ* hybridization (ISH) images from the Allen Brain Institute's Developing Mouse Brain Atlas (Henry and Hohmann, 2012).

Progression through the cell cycle requires proper regulation of the cytoskeleton. In *Nfib* OE, we observed increased expression of neuronal intermediate filaments Nes and Vim (Lendahl et al., 1990; Park et al., 2010) and tubulin-associated genes *Tuba1b* and *Tubb2a*, which were also enriched in late-born precursors. Additionally, *Sox2*, a marker of neuronal multipotent progenitors, was enriched in *Nfib* OE. In contrast, *Ezh2*, an enhancer of the polycomb repressive complex 2 (PRC2), was depleted in APs in *Nfib/x* KO, suggesting a potential relationship between NFIs and PRC2.

Markers of basal progenitors, such as the proneuronal gene Ascl1 and the cell cycle-associated gene *Ccnd2* (Vainorius et al., 2023; Tsunekawa et al., 2012), were depleted in progenitors in *Nfib/x* KO but enriched in Nfib OE. Ascl1 is known to promote cell cycle exit and neuronal differentiation; its upregulation could accelerate the transition from progenitors to post-mitotic precursors, potentially explaining the increased abundance of post-mitotic precursors in *Nfib* OE. A similar pattern was observed for members of the DLX family of transcription factors, which are critical regulators of inhibitory neuron development (Lindtner et al., 2019). In *Nfib* OE, DLX-family TFs showed increased expression in APs, BPs, and precursors, while in *Nfib/x* KO, their expression decreased in APs.

Among genes involved in migration, Slit1, Pak3, and Ctnna2 were depleted in APs in *Nfib/x* KO, while Ctnna2 was enriched in APs in Nfib OE. This trend reversed in BPs and precursors in Nfib OE, where most migration-related genes were significantly depleted.

The interneuron marker *Tcf4* was predicted to be activated by Nfib according to the eGRN. In Nfib OE, *Tcf4* expression exhibited cell state-dependent regulation, with depletion in BPs and interneurons but a slight increase in PNs. Similarly, the interneuron marker *Erbb4* (Batista-Brito et al., 2023) followed this pattern. By contrast, genes such as Maf and Prox1 remained unaffected by *Nfib/x* KO or *Nfib* OE.

Meis2, also predicted to be activated by *Nfib*, exhibited expected decreases in *Nfib/x* KO but cell state-dependent expression in *Nfib* OE, with reduced expression in BPs and PNs and increased expression in INs. Other PN markers, such as *Sp9* and *Ebf1*, increased in *Nfib* OE, with Ebf1 being depleted only in PNs.

### Supplementary Discussion

#### Transcriptomic correlation of clones

The correlation analysis was performed at the single cell level to capture possible heterogeneity in the progenitor population that might lead to more or less fate committed cells. However, the range of correlation scores in the mitotic fraction of non-dispersing clones does not indicate greater transcriptional similarity to their clonal output compared to randomly selected progenitors. This finding suggests that mitotic progenitor cells do not exhibit transcriptomic patterns that specify the developmental outcome observed in their clonal progeny. The current single-cell RNA-seq data may lack the resolution to detect subtle gene signatures linked to specific clonal outputs. Future studies using higher-resolution sequencing approaches, combined with perturbation experiments, will be required to determine whether non-dispersing clones harbor distinct transcriptional programs or reflect broader heterogeneity within the progenitor pool.

#### Sampling strategy for inhibitory precursors

We aimed to include postmitotic cells both during and after their migration to their final destinations. To achieve this, we collected tdTomato+ cells from the cortex and striatum separately because this allowed us to specifically enriched for the relatively rare cortical interneurons (INs), which are typically underrepresented compared to the abundant GABAergic projection neurons (PNs) in the striatum. It is important to note that spatial aspects were not the focus of this study. Our analysis includes precursor states for INs that represent precursors of both striatal and cortical INs. At the embryonic stages we studied, these populations cannot be distinguished based on their transcriptome, as we have previously demonstrated (Bandler and Mayer, 2023). Furthermore, pooling cells collected from the cortex and striatum allowed us to minimize technical batch effects. In the revised manuscript, we discuss this in the supplementary data.

#### Fractions of post-mitotic precursors

In wildtype scRNA-seq experiments as well as in CFSE-cohorts and lineage-tracing experiments, we consistently observed all states of post-mitotic precursors. Qualitatively, this shows the maintained differentiation competence of progenitors throughout neurogenesis. On a quantitative level, we observed some cell states being very rare, especially in the CFSE-cohorts. We argue that this observation is confounded by varying maturation levels of CFSE cohorts, i.e. that cells have not yet reached the maturity of the cell states of this analysis (branch tips), as illustrated in Extended Data Fig. 3b. Another confounder is our sampling strategy which enriches for GABAergic cells of the striatum and cortex, and therefore is not an unbiased representation of all GABAergic neurons.

#### Cell loss during transplantation

Cell loss during transplantation may result in a subset of CFSE+ cells failing to integrate, potentially biasing the observed pseudotime and gene expression patterns toward those of the recipient stage. A limitation of this technique is the possible failure of a subset of CFSE+ cells to integrate into the tissue during transplantation, which will lead to cell loss. Thereby, potentially biasing observed pseudotime and gene expression patterns. However, we tried to minimize this effect by sorting CFSE+1h cohorts to enrich for APs. Further investigation is needed to uncover the mechanisms through which GABAergic APs synchronize with the host environment.

### Supplementary Figures

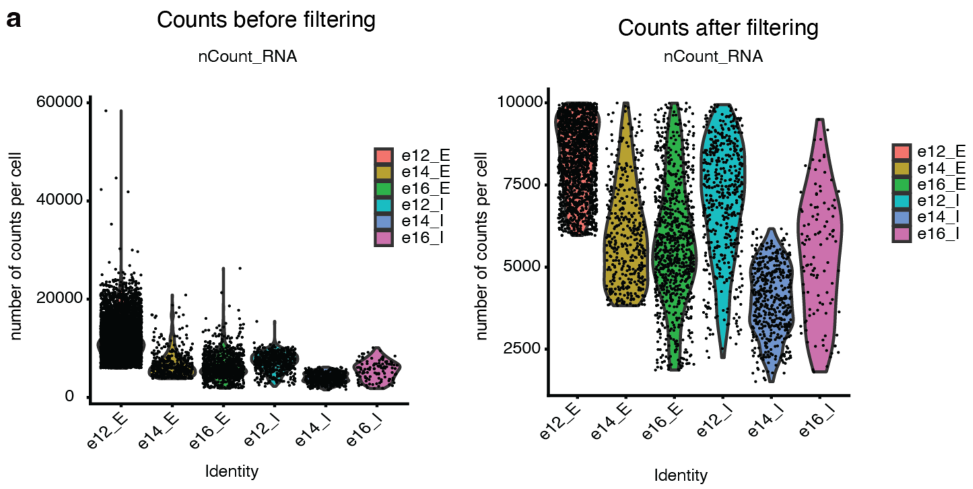


**Supplementary Fig. 1: Filtering of scRNA-seq datasets. a - b**, Distribution of number of counts per sample before applying filtering (**a**) and after applying filtering (**b**).

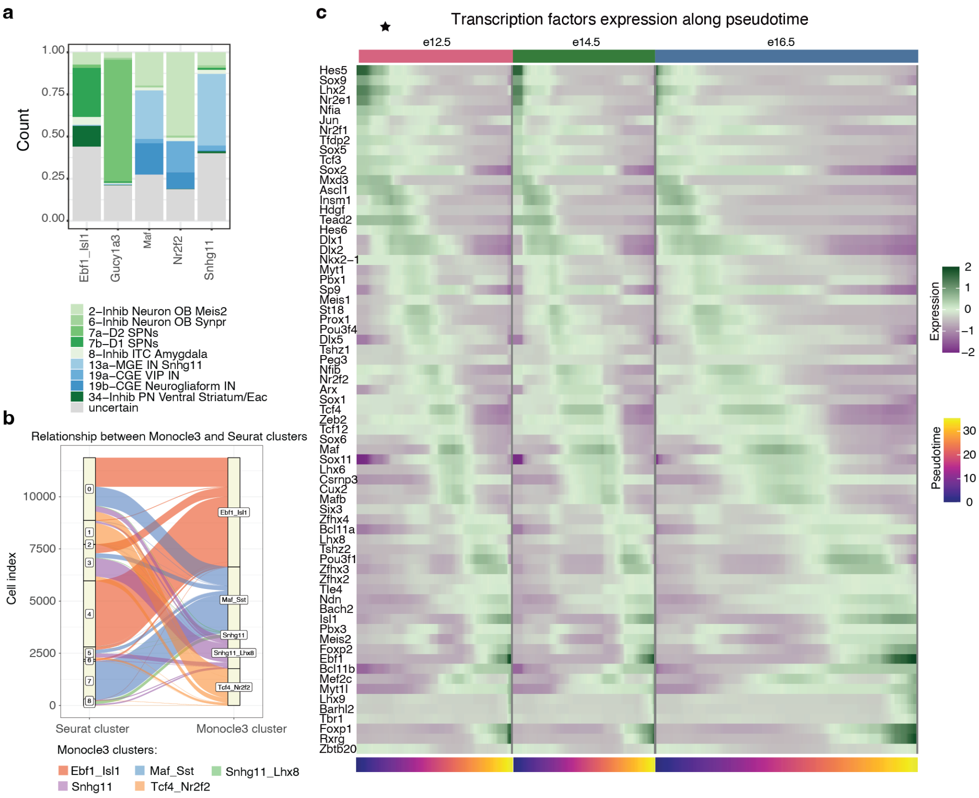


**Supplementary Fig. 2: Transcriptional progression of GABAergic lineage. a**, Mapping of post-natal inhibitory neuron identities to cells in branch clusters using label transfer. For each branch cluster (x-axis) the fraction of mapped post-natal identities is shown. Cells with low prediction scores are annotated as 'uncertain'. **b**, Relationship between Seurat and Monocle3 clusters. Flow diagram showing how cluster membership changes between Seurat and Monocle3. Graph only contains cells that are included in branch clusters (from Monocle3). Colour indicates branch cluster. **c**, Scaled and smoothed expression of dynamic TFs in GABAergic cells. TFs are ordered by their maximum expression along the pseudotime trajectory.

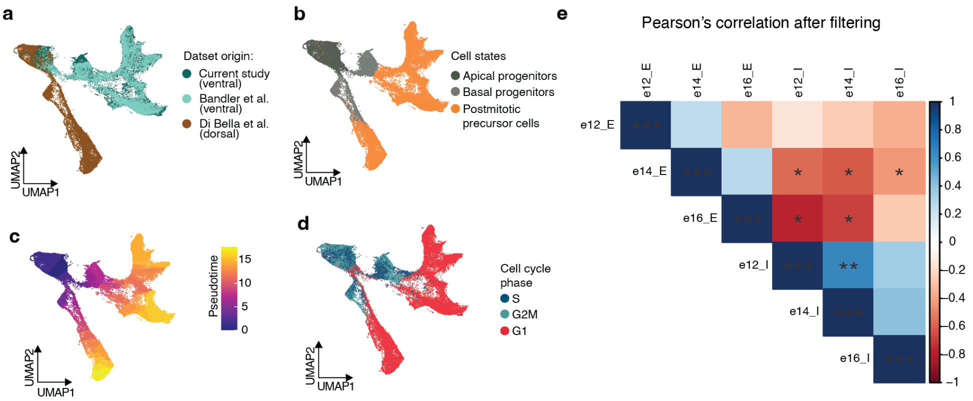


**Supplementary Fig. 3: Developmental dynamics in ventral and dorsal lineages.** **a**, UMAP plot of ventral and dorsal lineage datasets from different publications. **b-d**, UMAP plot of ventral and dorsal lineage datasets, with cells coloured by broad cell state (**b**), pseudotime (**c**) and cell cycle phase (**d**). **e**, Pearson correlation between dorsal and ventral APs at different stages. Correlation coefficients were calculated after controlling for sequencing depth via sub-sampling; * $P<0.05$, ** $P<0.01$.

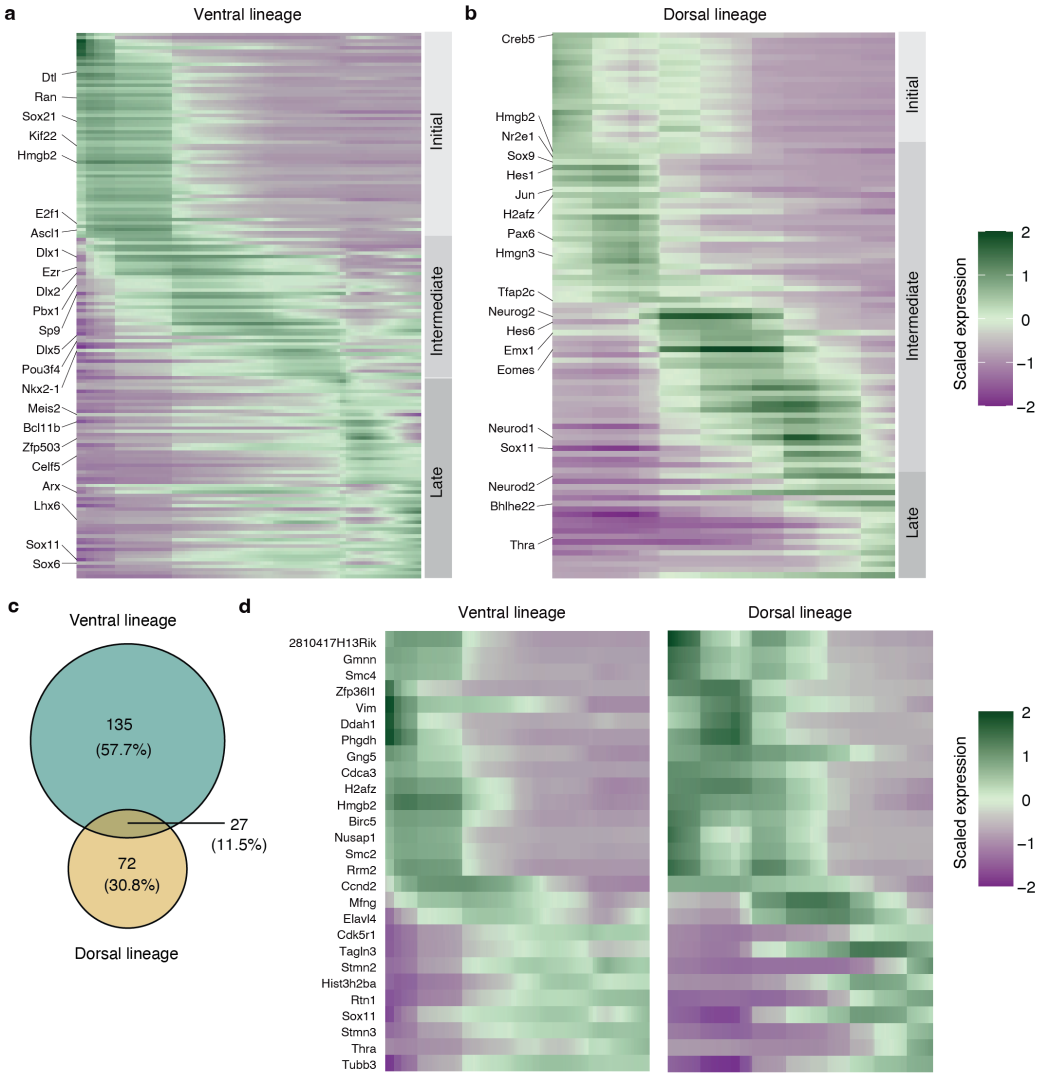


**Supplementary Fig. 4: Dynamic genes in ventral and dorsal lineages.** **a**, Scaled and smoothed expression of genes dynamic along the maturation trajectory in ventral lineage. **b**, Scaled and smoothed expression of genes dynamic along the maturation trajectory in dorsal lineage. **c**, Number of shared and distinct dynamic genes between ventral and dorsal lineages. **d**, Expression of common dynamic genes in dorsal and ventral lineages.

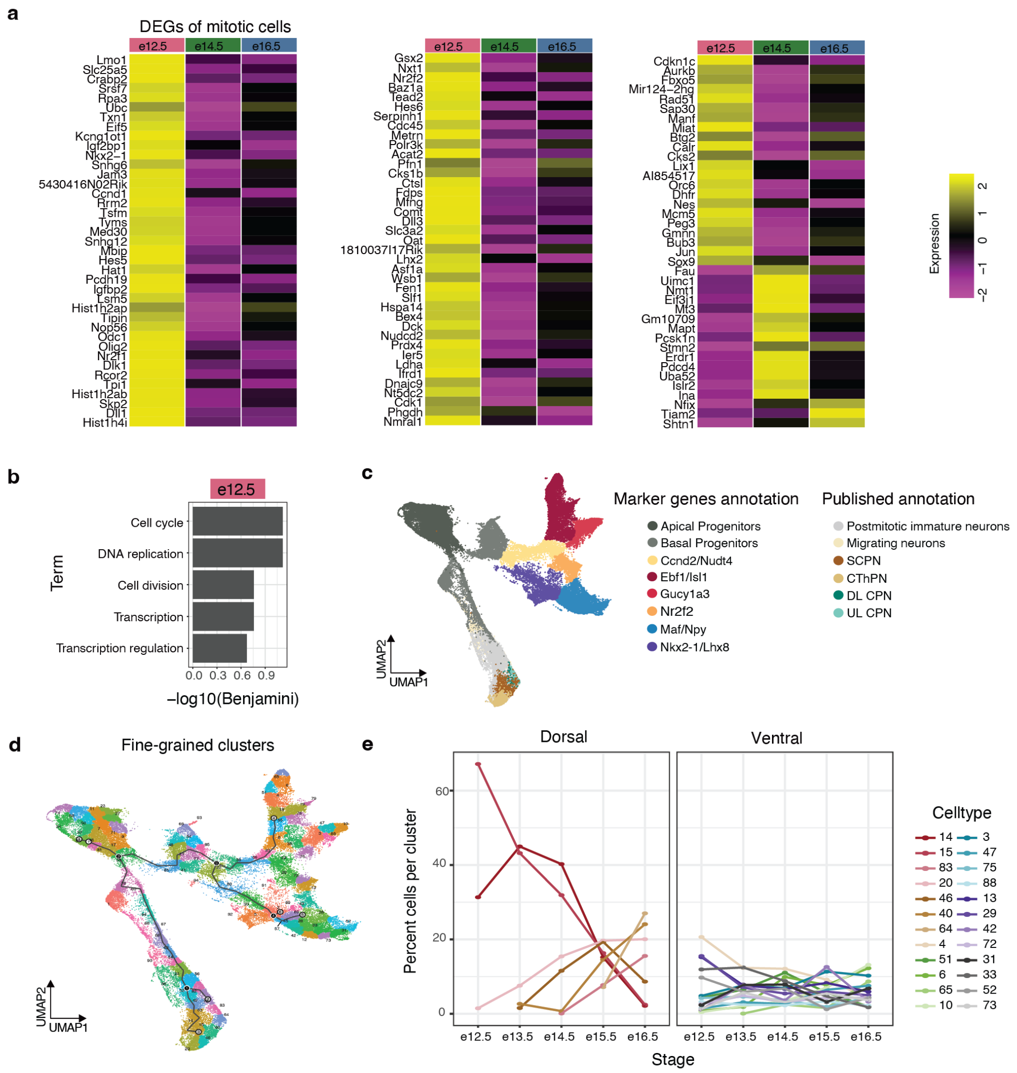

**Supplementary Fig. 5: Temporal dynamics of gene expression and cell state abundance.** **a**, Average gene expression heatmap of differentially expressed genes in mitotic progenitors at e12.5, e14.5, and e16.5 in the scRNAseq dataset. **b**, Functional annotation of differentially expressed genes in e12.5 APs. Top five GO terms ranked by Benjamini-Hochberg corrected *P* values (DAVID). **c**, UMAP plot of ventral and dorsal lineages. The dorsal lineage is annotated according to Di Bella *et al.* (Di Bella et al., 2021). The ventral lineage is annotated based on top marker gene expression. **d**, UMAP plot of ventral and dorsal lineages with trajectory inferred with Monocle3. Cells are grouped and annotated by fine-grained clusters. **e**, Abundance of fine-grained clusters throughout all stages (e12.5 - e16.5) in dorsal and ventral lineages. Line color indicates cluster membership, which is ordered by dorsal vs. ventral lineage.

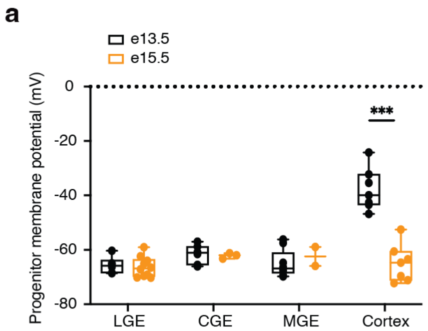


**Supplementary Fig. 6: Resting membrane potential of progenitors across spatial domains. a**, Progenitor membrane potential in LGE, CGE, MGE and cortex at e13.5 and e15.5; two-sided t-test; *** *P*<0.001.

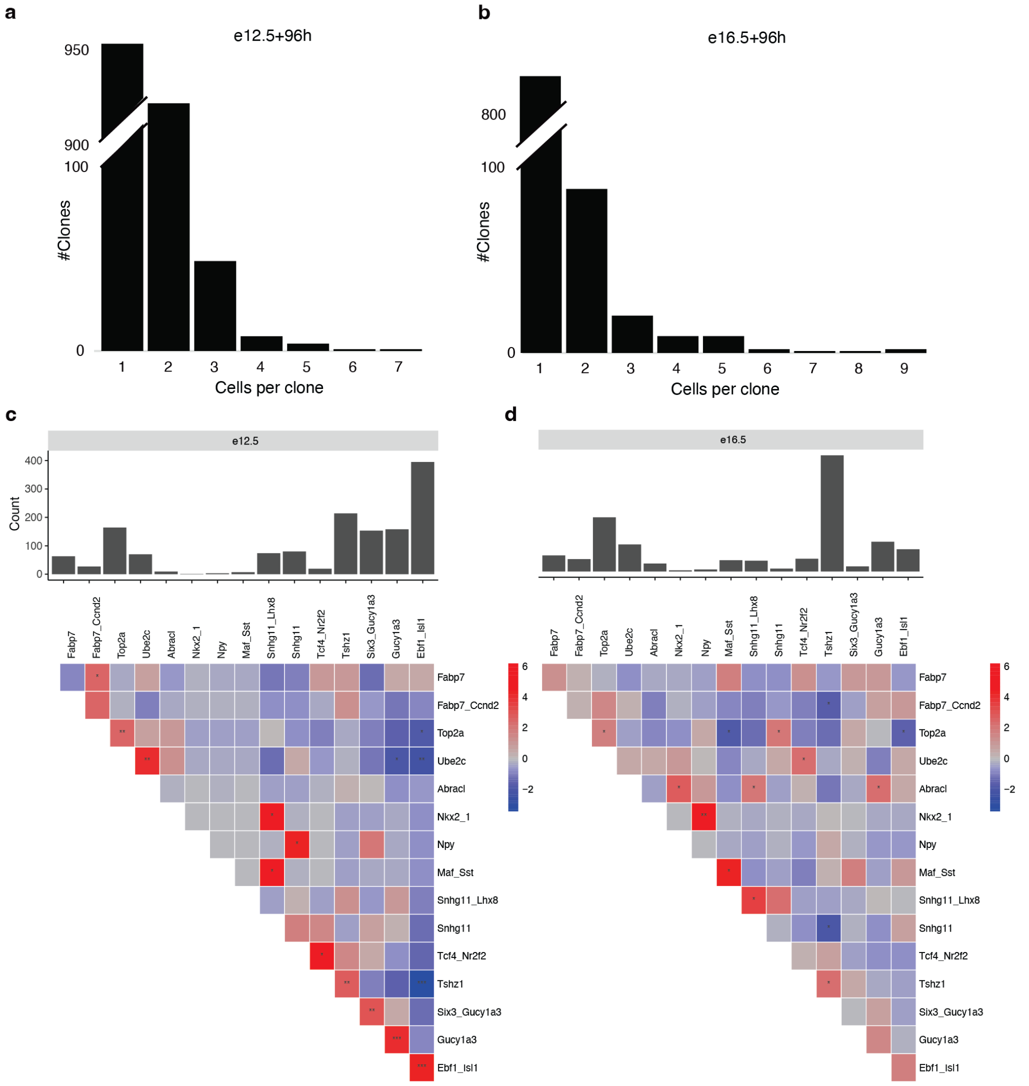


**Supplementary Fig. 7: Clonal distribution and coupling in TrackerSeq datasets.** **a**, Clonal distribution in the TrackerSeq$_{e12.5+96h}$ dataset (left) and TrackerSeq$_{e16.5+96h}$ dataset (right). **b**, Barplots showing the number of cells containing clonal information per cluster in e12.5 (left) and e16.5 (right). **c**, Clonal coupling between cell states in TrackerSeq$_{e12.5+96h}$ (left) and TrackerSeq$_{e12.5+96h}$ (right). Z-scores indicate clonal coupling, with positively coupled pairs of cell states having positive z-scores and anticoupled pairs of cell states having negative z-scores. Empirical *P* values were calculated for each comparison and FDR-corrected (*** *P*<0.001).

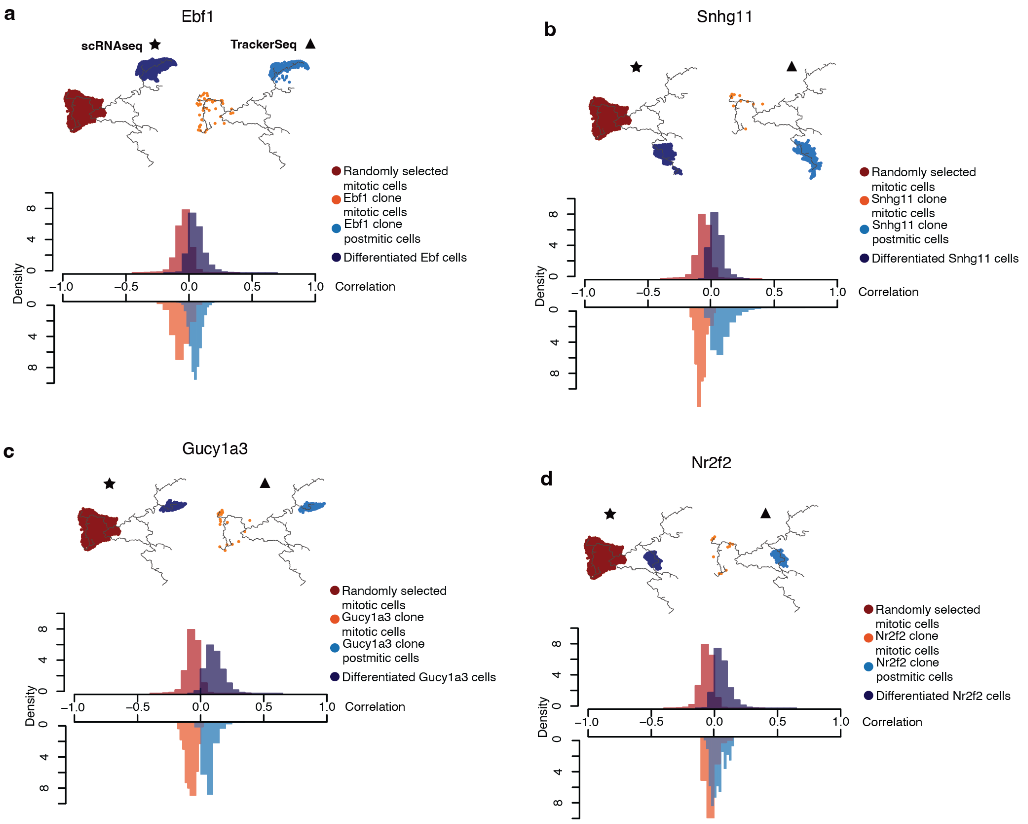


**Supplementary Fig. 8: Analysis of TrackerSeq-labelled clones. a-d**, Correlation analysis of "non-dispersing" TrackerSeq-labelled clones. Pearson correlation coefficient calculated between cell pairs within mitotic and postmitotic clone subsets, and reference subsets. **a**, Postmitotic clones of the Ebf1 cluster, mitotic clones of the Ebf1 cluster, and randomly selected mitotic cells, correlated to the Ebf1 postmitotic reference group. **b**, Postmitotic clones of the Snhg11 cluster, mitotic clones of the Snhg11 cluster, and randomly selected mitotic cells, correlated to the Snhg11 postmitotic reference group. **c**, Postmitotic clones of the Gucy1a3 cluster, mitotic clones of the Gucy1a3 cluster, and randomly selected mitotic cells, correlated to the Gucy1a3 postmitotic reference group. **d**, Postmitotic clones of the Nr2f2 cluster, mitotic clones of the Nr2f2 cluster, and randomly selected mitotic cells, correlated to the Nr2f2 postmitotic reference group.

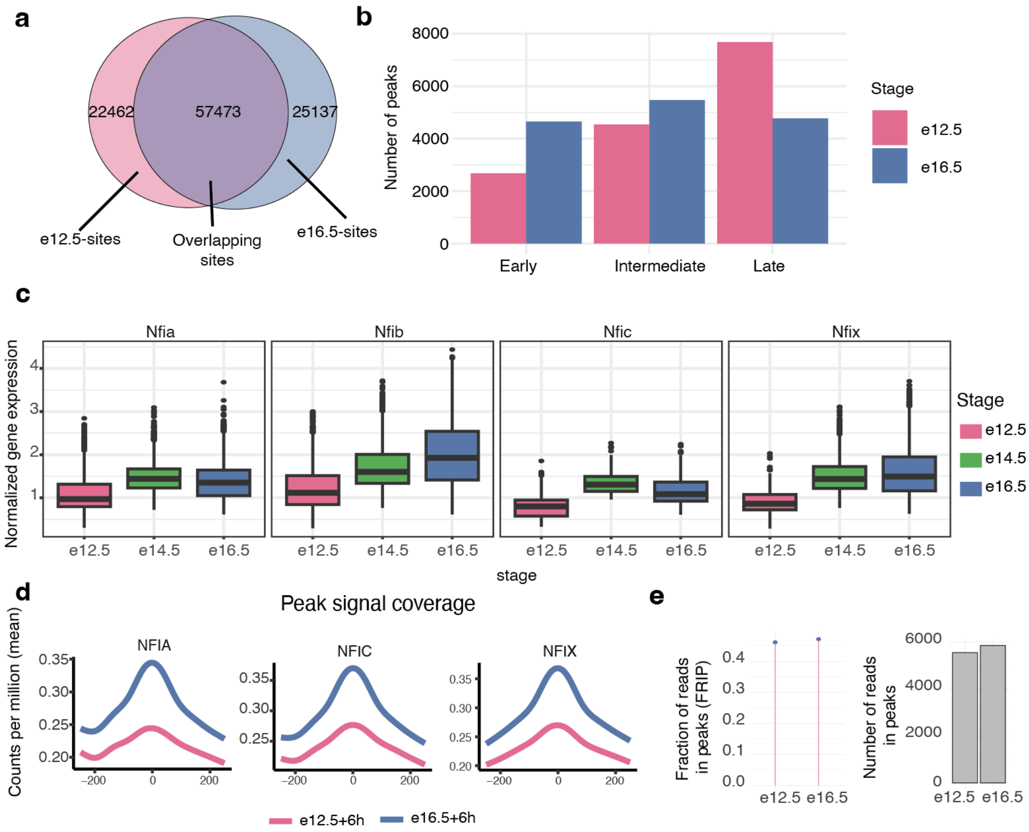


**Supplementary Fig. 9: Chromatin dynamics and associated dynamics of TFs.** **a**, Venn diagram showing the three categories of sites (e12.5 sites, e16.5 sites and overlapping sites) and the number of non-overlapping and overlapping sites. **b**, Barplot quantifying peak distribution across initial, intermediate, and late phases of pseudotime. **c**, Boxplot showing normalised gene expression levels at e12.5, e14.5 and e16.5; scRNA-seq dataset. Cells with no expression of the selected genes are not displayed. **d**, Coverage plot showing chromatin accessibility dynamics at Nfib, Nfia, Nfic and Nfix footprint sites in $FT_{e12.5+6h}$ and $FT_{e16.5+6h}$ datasets. **e** Fraction and absolute number of reads that are contained within e12.- or e16.5-enriched peaks.

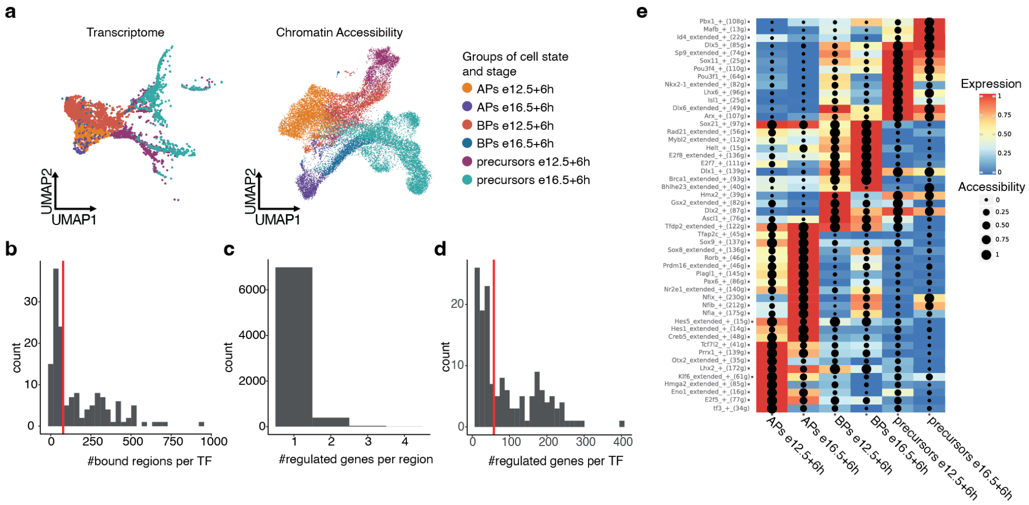


**Supplementary Fig. 10: Gene regulatory network of $\mathbf{FT_{e12.5+6h}}$ and $\mathbf{FT_{e16.5+6h}}$.** **a**, UMAP embedding of cells from scRNA-seq (left) and scATAC-seq (right). Cells are annotated based on broad cell state and stage. **b-d**, Distribution of the number of bound regions per TF (**b**), regulated genes per region (**c**), and target genes per TF (**d**). Red horizontal lines in (**b**) and (**d**) indicate the median of the corresponding distribution. **e**, Heatmap displaying active marker modules for stage-specific cell states. Colour indicates the scaled average expression of the module (i.e. TF expression and target genes expression). Dot size indicates scaled accessibility of associated regulatory regions. Modules are named after the TF, type of regulation (here only positive) and number of target genes.

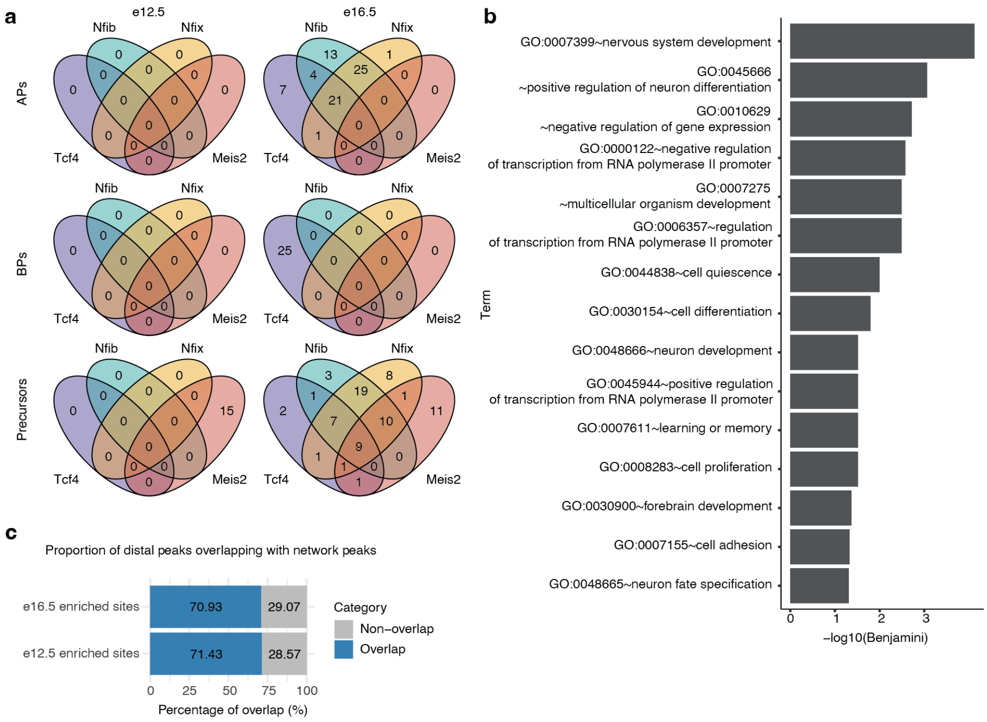


**Supplementary Fig. 11: Shared target genes of Nfib, Nfix, Meis2 and Tcf4.** **a**, Number of shared and distinct regulated target genes of Nfib, Nfix, Tcf4 and Meis2 across cell states and stages. **b**, Significantly enriched GO-terms (Benjamini-Hochberg corrected $P < 0.05$) for target genes of Nfib and Nfix and/or Tcf4 in e16.5 cells. **c**, Fraction of e12.5- or e16.5-enriched peaks that overlap with sites that are contained in the eGRN.

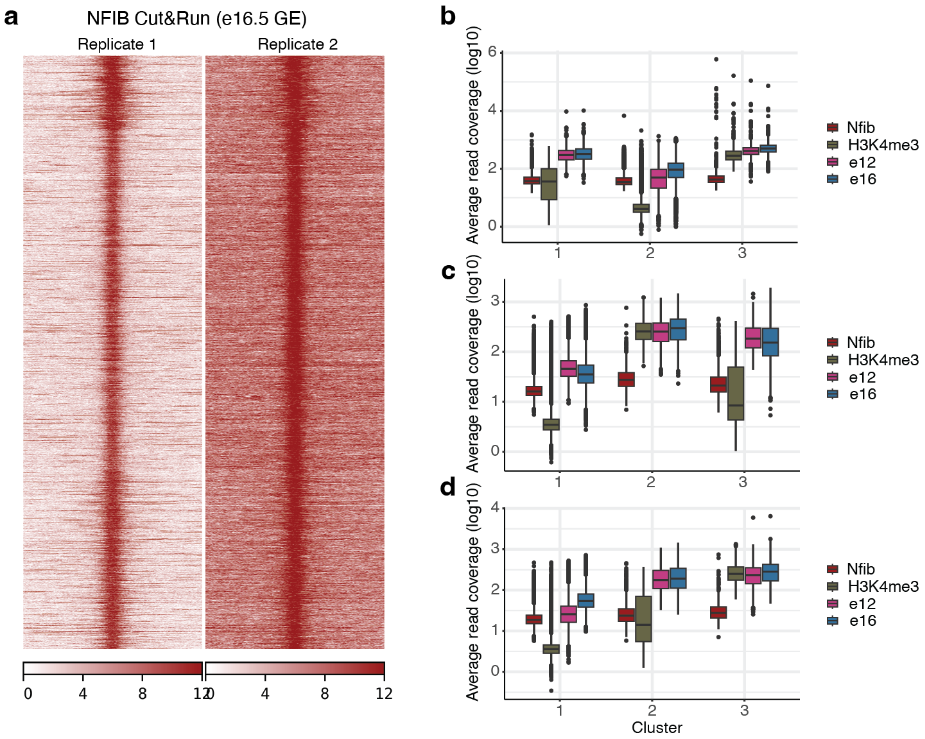


**Supplementary Fig. 12: Genomic binding of NFIB. a**, Coverage at NFIB binding peaks across both replicates. **b-d**, Aggregated coverage at specific sites of NFIB CUT&RUN, H3K4me3 CUT&RUN and chromatin accessibility of $FT_{e12.5 + 6h}$ and $FT_{e16.5 + 6h}$. Read coverage was normalized using RPKM and aggregated by calculating the mean coverage per peak region (peak center +/- 500bp). Aggregated coverage was log10-transformed and is shown for each cluster of peaks. In (**a**), aggregated read coverage is shown at NFIB CUT&RUN peaks; in (**b**), coverage is shown at e12.5 enriched sites and in (**c**), coverage is shown at e16.5 eniched sites.

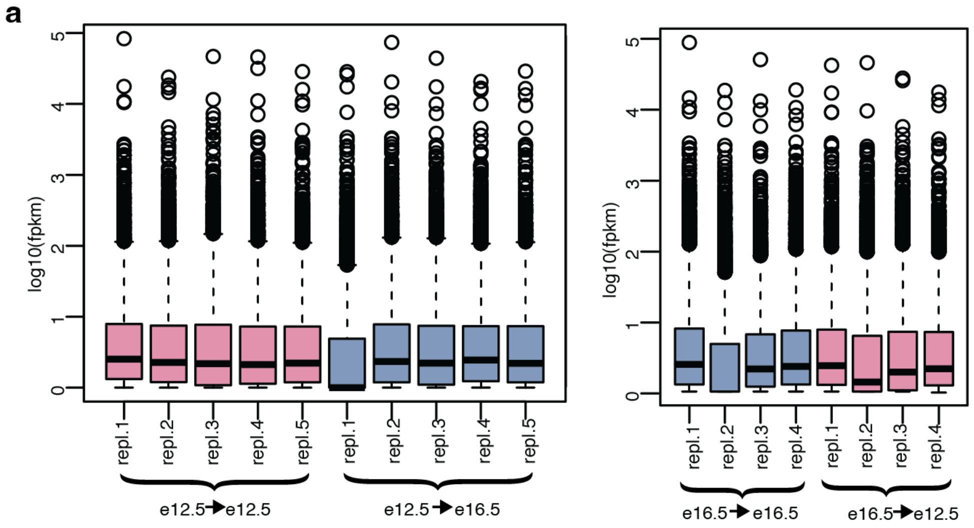


**Supplementary Fig. 13: Normalization of RNA-seq in transplantationexperiments a**, FPKM normalization of count matrices from bulk RNA sequencing.

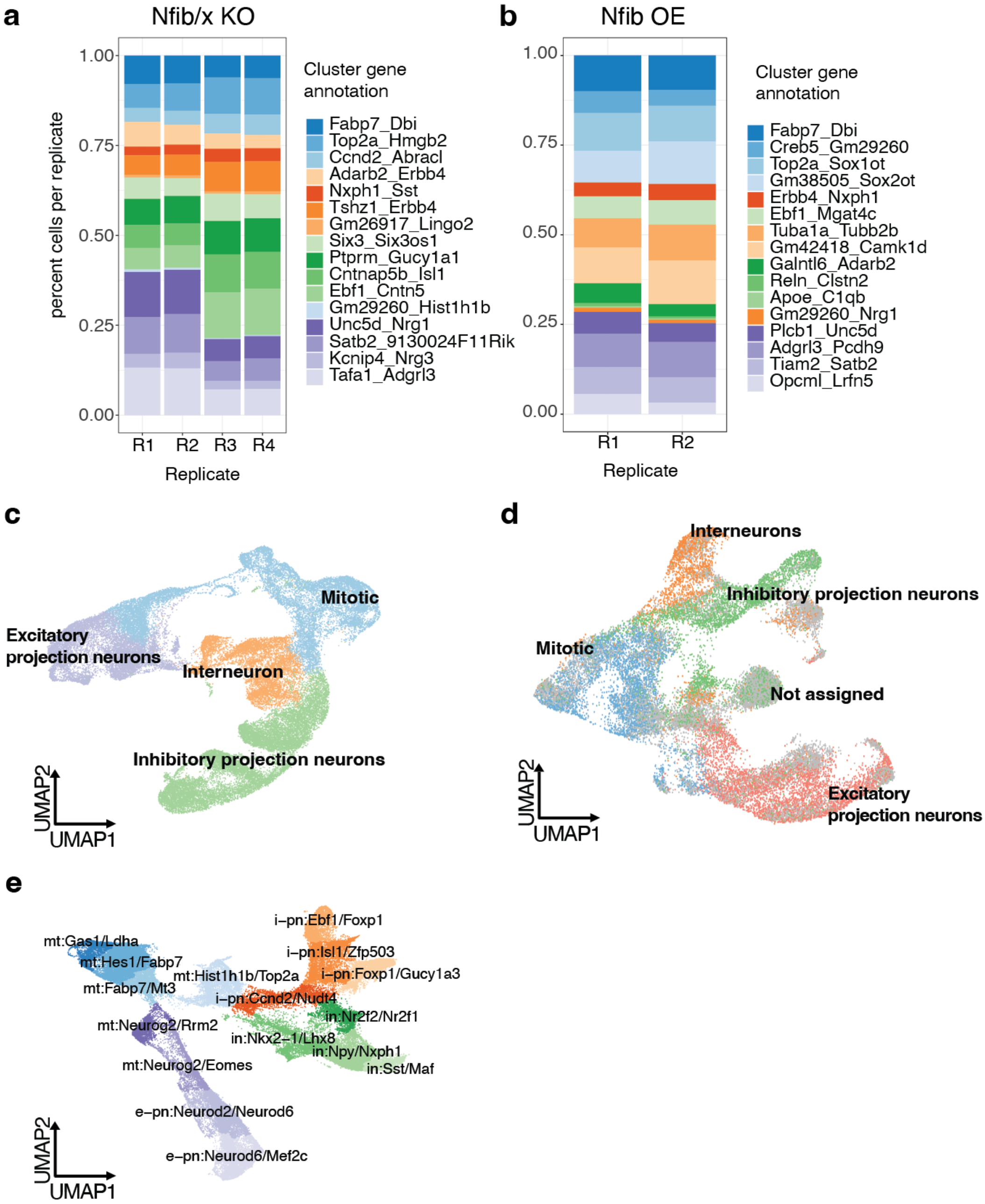


**Supplementary Fig. 14: Cell states in Nfib/x KO and Nfib OE.** **a**, Relative fraction of clusters across replicates in Nfib/x KO. Replicates R1/R2 and R3/R4 are technical replicates of two biological replicates. **b**, Relative fraction of clusters across replicates in Nfib OE. **c**, Broad cell state annotation in Nfib/x KO. Broad cell states were generated by aggregating cluster annotations. **d**, Broad cell state annotation in Nfib OE. Broad cell states were generated by aggregating predicted label annotation. **e**, UMAP-embedding of integrated wild-type datasets of inhibitory and excitatory lineages that was used as a reference for label transfer. Cells are annotated by cluster.

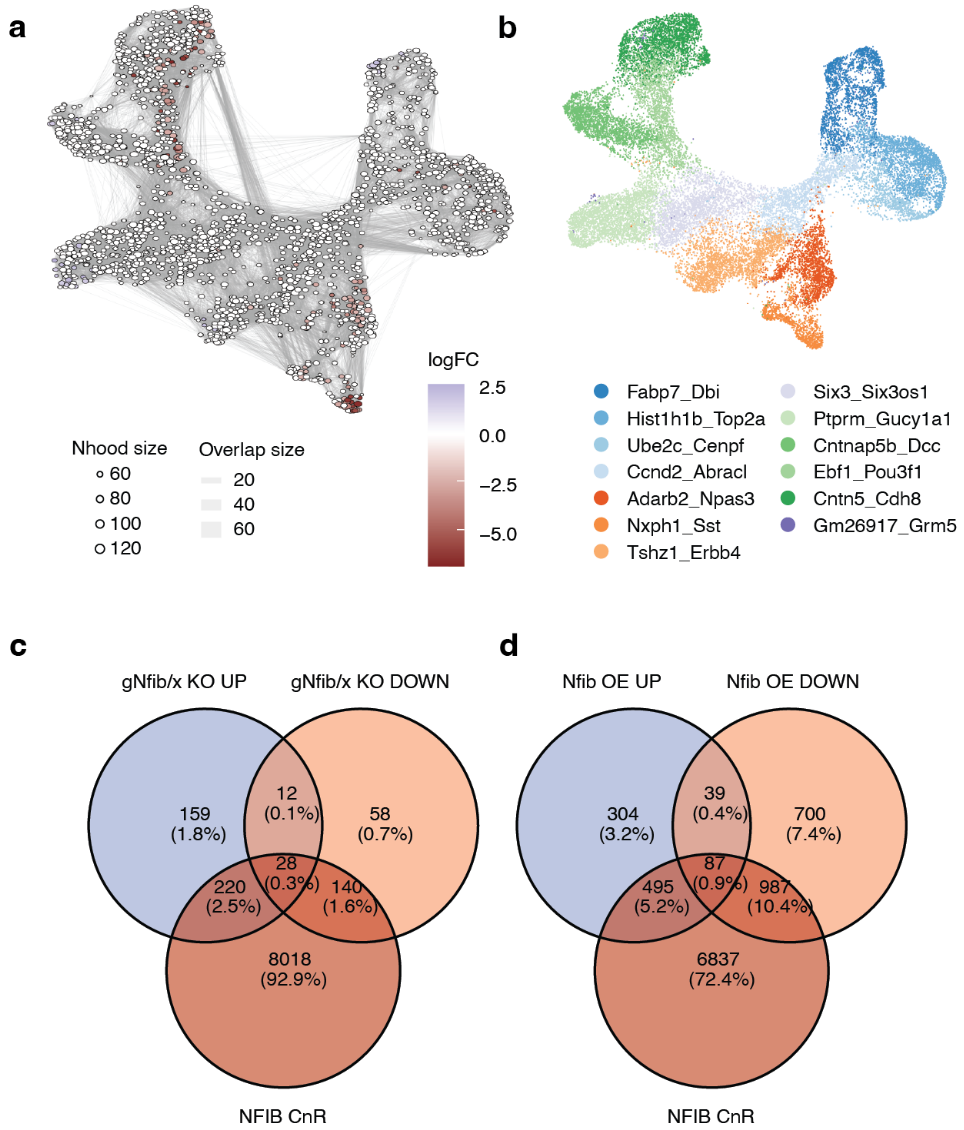


**Supplementary Fig. 15: Differential abundance in cell-neighbourhoods of the ventral lineage upon Nfib/x KO a**, UMAP embedding of cell neighbourhoods in the inhibitory subset of Nfib/x KO. Each dot represents a neighbourhood of cells and lines between neighbourhoods indicate the amount of shared cells. Differential abundance upon Nfib/ KO is indicated by color, with positive FC indicating increased abundance when knocking-out and vice-versa. **b**, UMAP embedding of cells in the ventral lineage that contain either gNfib/x or gLacZ. Cells are annotated by clusters. **c**, Overlap between DE-genes in Nfib/x KO and genes that are directly bound by NFIB at their promoter (according to NFIB CUT&RUN). **d**, Overlap between DE-genes in Nfib OE and genes that are directly bound by NFIB at their promoter (according to NFIB CUT&RUN).

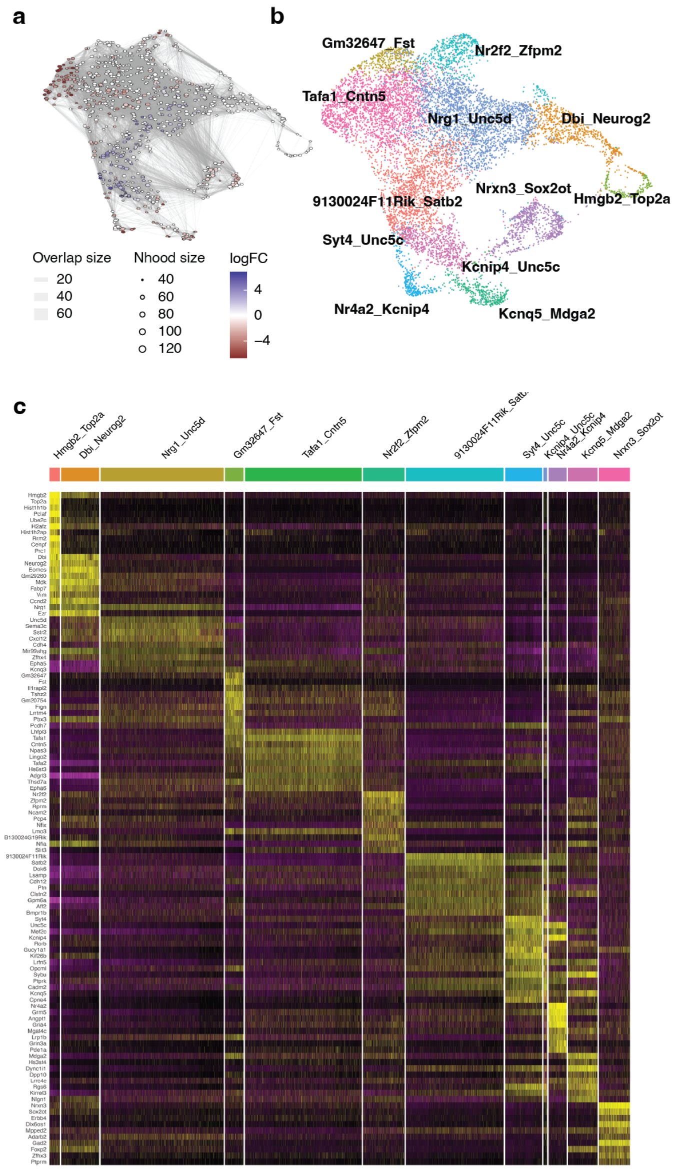


**Supplementary Fig. 16: Differential abundance in cell-neighbourhoods of the dorsal lineage upon Nfib/x KO a**, UMAP embedding of cell neighbourhoods in the excitatory subset of Nfib/x KO. Each dot represents a neighbourhood of cells and lines between neighbourhoods indicate the amount of shared cells. Differential abundance upon Nfib/ KO is indicated by color, with positive FC indicating increased abundance when knocking-out and vice-versa. **b**, UMAP embedding of cells in the dorsal lineage that contain either gNfib/x or gLacZ. Cells are annotated by clusters. **c**, Heatmap showing top 10 marker genes per cluster of the dorsal subset of Nfib/x KO.

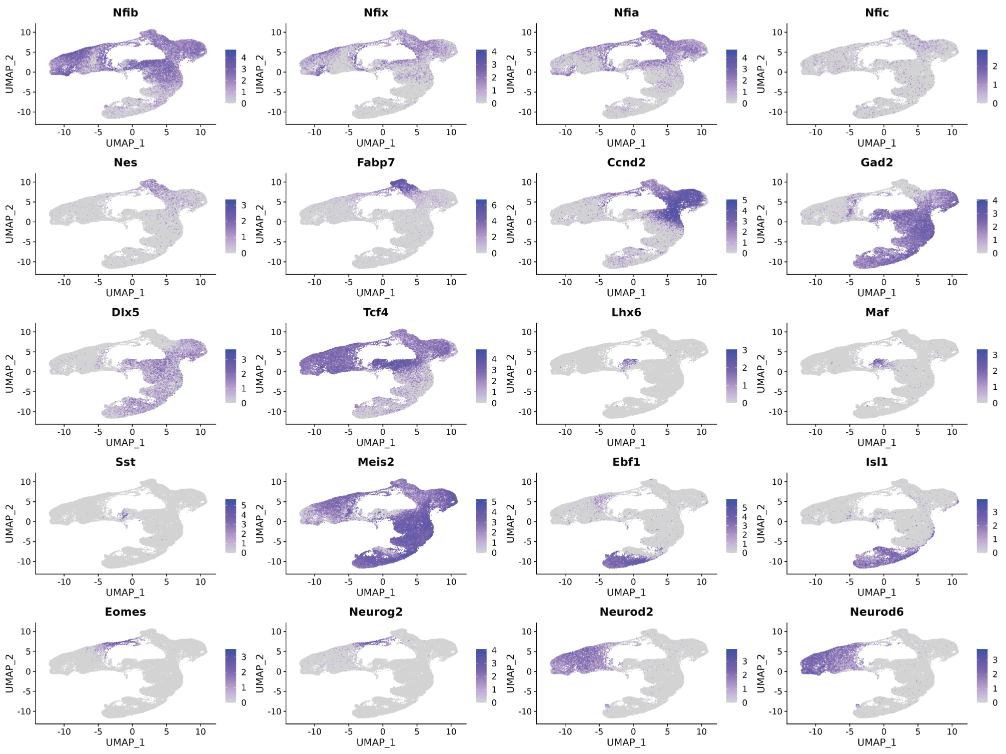


**Supplementary Fig. 17: Expression patterns of marker genes in *Nfib/x* KO.** Log-normalized expression of cells is shown according to cell's position in UMAP-embedding: Members of the NFI family of TFs (*Nfib, Nfix, Nfia, Nfic*), ventral mitotic markers (*Fabp7, Nes, Ccnd2*), post-mitotic precursors of inhibitory neurons (*Gad2, Dlx5*), interneuron precursors (*Tcf4, Lhx6, Maf, Sst*), inhibitory projection neuron precursor (*Meis2, Ebf1, Isl1*), dorsal intermediate progenitors (*Eomes, Neurog2*) and excitatory projection neuron precursors (*Neurod2, Neurod6*).

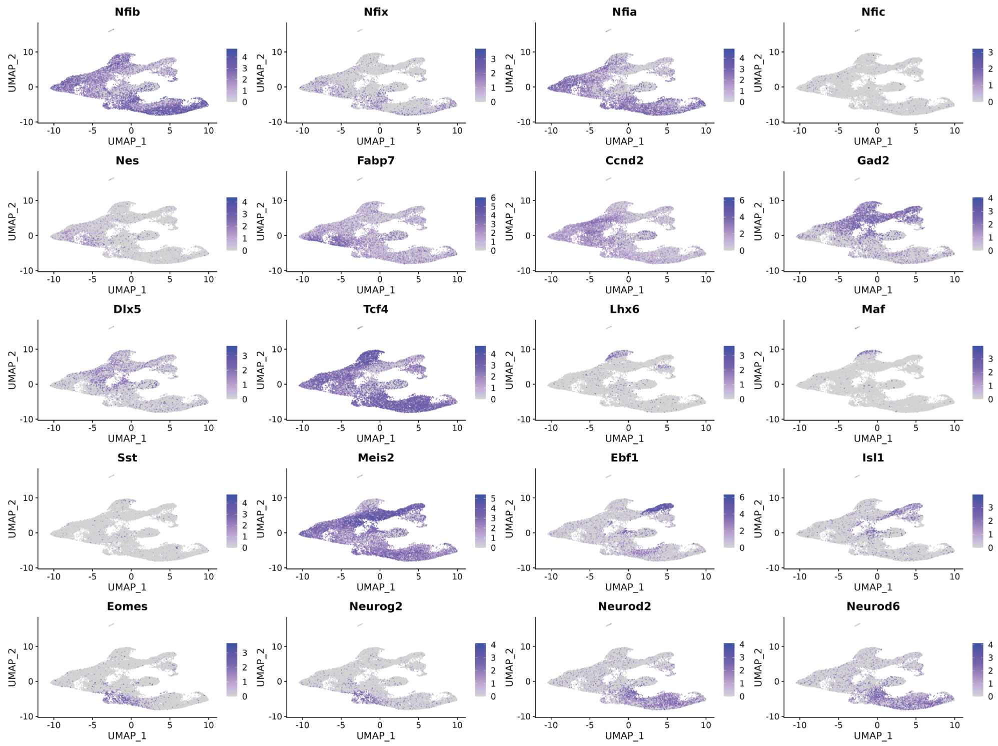


**Supplementary Fig. 18: Expression patterns of marker genes in Nfib OE.** Log-normalized expression of cells is shown according to cell's position in UMAP-embedding: Members of the NFI family of TFs (*Nfib, Nfix, Nfia, Nfic*), ventral mitotic markers (*Fabp7, Nes, Ccnd2*), post-mitotic precursors of inhibitory neurons (Gad2, Dlx5), interneuron precursors (*Tcf4, Lhx6, Maf, Sst*), inhibitory projection neuron precursor (*Meis2, Ebf1, Isl1*), dorsal intermediate progenitors (*Eomes, Neurog2*) and excitatory projection neuron precursors (*Neurod2, Neurod6*).

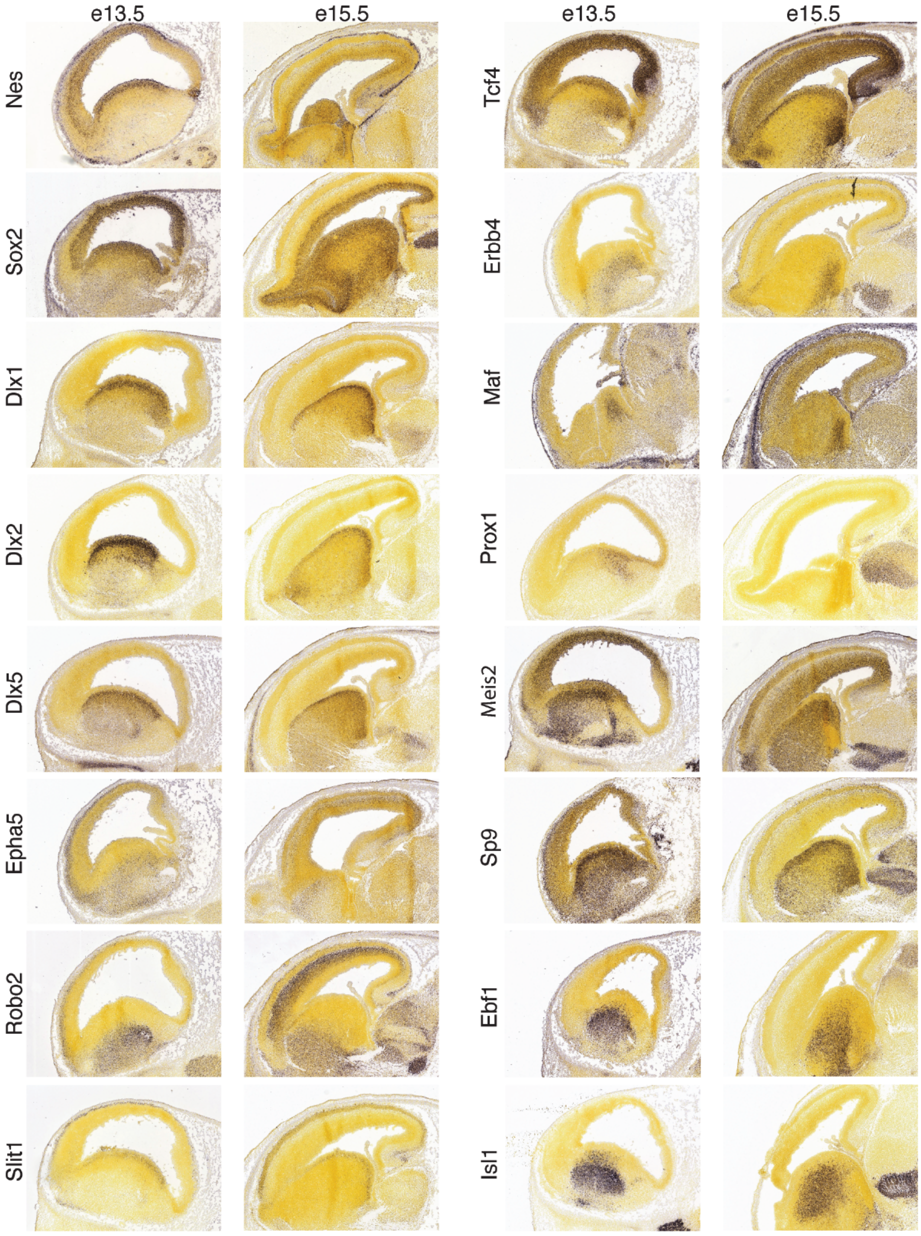


**Supplementary Fig. 19: In-situ Hybridization (ISH).** ISH-pictures of Allen Brain Institute's Developing Mouse Brain Atlas at e13.5 and e15.5 showing coronal sections of genes contained within Fig. 4k.

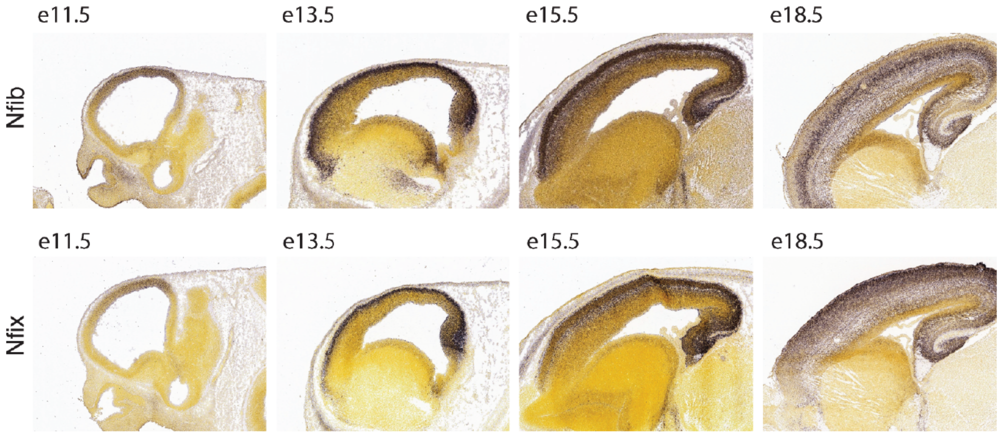


**Supplementary Fig. 20: ISH of Nfib and Nfix.** ISH-pictures from Allen Brain Institute's Developing Mouse Brain Atlas at e13.5 and e15.5 of Nfib and Nfix.

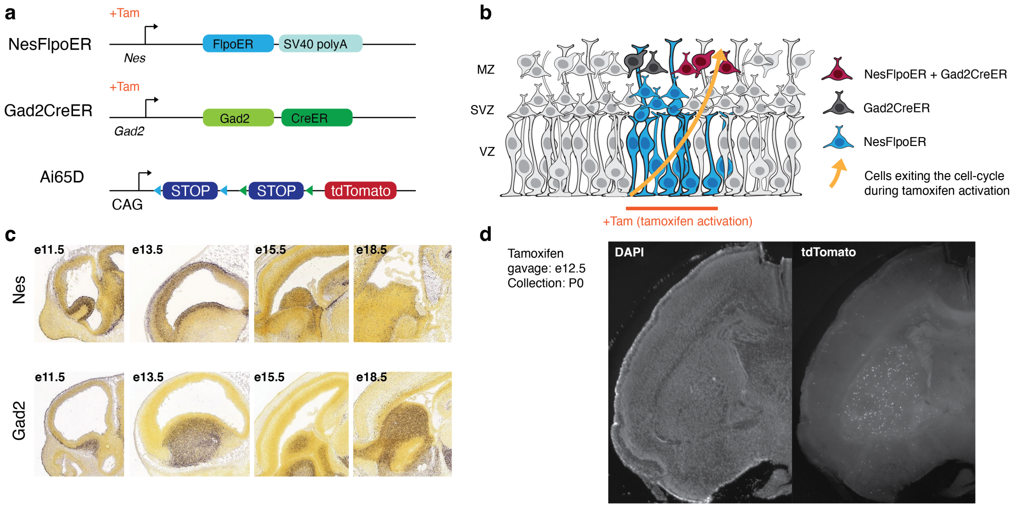


**Supplementary Fig. 21: An intersectional approach for GABAergic neuron birthdating.** **a**, Schematic illustrating the crossing of alleles: NesFlpoER allele, Gad2CreER allele, and Ai65D intersectional reporter allele. **b**, Schematic illustrating the rationale behind the birthdating method: administration of tamoxifen inducing the expression of tdTomato in a cohort of cells that was transitioning from a mitotic (Nes$^{+}$) to a postmitotic (Gad2$^{+}$) cell state. **c**, *In situ* hybridization images from the Allen Brain Atlas; sagittal sections. **d**, Coronal sections of brains with DAPI$^{+}$ (right) and tdTomato$^{+}$ (left) cells; tamoxifen gavage at e12.5, collection at P0.

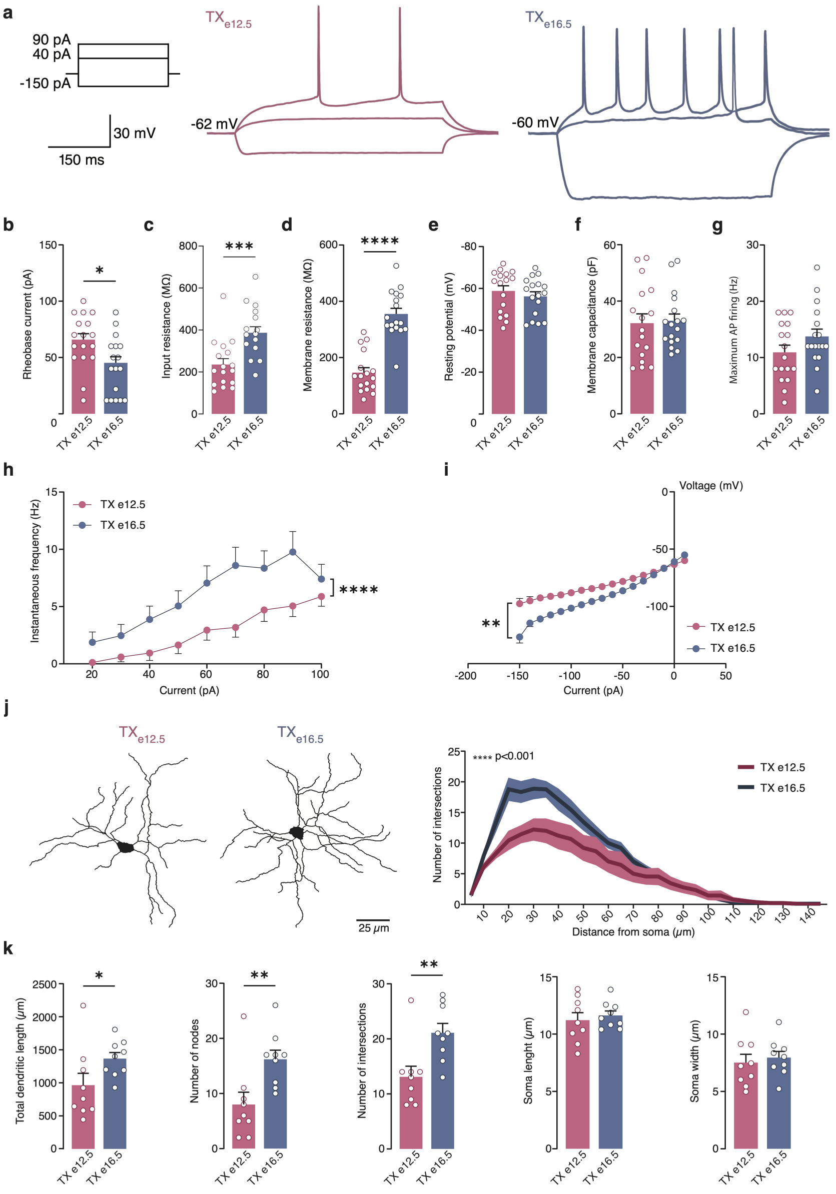


**Supplementary Fig. 22: Electrophysiological and morphological characterisation of TX$_{e12.5}$ and TX$_{e16.5}$ cohorts.** **a**, Representative voltage responses to hyperpolarising (–150 pA) and depolarising (40 and 90 pA) current pulses in TX$_{e12.5}$ and TX$_{e16.5}$ GABAergic neurons. **b–d**, Summary bar graphs of rheobase current (b), input resistance (c) and membrane resistance (d) in TX$_{e12.5}$ and TX$_{e16.5}$ GABAergic neurons. **e–g**, Summary bar graphs of resting potential (e), membrane capacitance (f) and maximum action potential (AP) firing (g) in TX$_{e12.5}$ and TX$_{e16.5}$ GABAergic neurons. **h**, Action potential firing frequency (Hz) plotted as a function of injected current steps (I-F curve). **i**, Current-voltage (I-V) plots recorded from TX$_{e12.5}$ and TX$_{e16.5}$ GABAergic neurons. **j**, Sholl analysis of reconstructed biocytin-filled GABAergic neurons recorded in the dorsal striatum region. Representative images of TX$_{e12.5}$ and TX$_{e16.5}$ neurons are shown on the left. The right panel shows the number of dendritic branches (arbor complexity) plotted against distance from the cell body (soma).

**Supplementary Fig. 22:**

**k**, (From left to right) Summary of the total dendritic length, number of nodes, maximum number of intersections, soma length and soma width in $TX_{e12.5}$ and $TX_{e16.5}$ neurons. Plots represent mean ± s.e.m. and raw data points. Mann–Whitney test for all panels except for panels h,i and j (Two-way ANOVA with multiple comparisons). * $P$<0.05, ** $P$<0.01, *** $P$<0.001, **** $P$<0.0001.

### Supplementary Tables

| Gene symbol | $log_2FC$ | Function during neurogenesis | Reference (PMID) |
|---|---|---|---|
| Tubb2a | 2,06 | Microtubule protein involved in neuronal migration and proliferation; mutation linked to brain malformation and epilepsy. | 24702957 |
| Rtn1 | 1,33 | Reticulon protein localized in ER and neuronal dendrites, regulates ER activity and cytosolic calcium dynamics. | 23454728, 23559015 |
| Nfix | 2,10 | Transcription factor essential for normal brain development, regulates progenitor cell differentiation in the hippocampus. | 23042739, 18477394 |
| Mapt | 2,08 | Microtubule-associated protein with role in brain development and neurogenesis as well as detrimental role in neurodegeneration at later age. | 29202785 |
| Nfib | 2,79 | Transcription factor involved in neuronal radial glia differentiation and cortical development regulation. | 23749646, 32166136 |
| Thra | 1,40 | Nuclear hormone receptor required for normal neural progenitor cell proliferation in human cerebral cortical development. | 31628250 |
| Ly6h | 1,42 | Endogenous neurotoxin-like protein, inhibits alpha7 nicotinic acetylcholine receptor currents at the plasma membrane. | 25716842, 32686737 |
| Zfp704 | 1,51 | C2H2 zinc finger protein expressed in several NC-derived lineages. | 17693064 |
| Myt1l | 1,23 | CCHC zinc finger TF involved in neuronal identity and maturation. | 21617644, 34614421 |
| Ncam1 | 1,48 | Neural cell adhesion molecule important for neuronal migration, neurite developent, synaptogenesis and neural stem cells regulation. | 19788570, 20038681, 17682066 |
| Stmn2 | 1,14 | Microtubule regulator, necessary for normal axonal outgrowth , regeneration and protection. | 30643292 |
| Nrxn3 | 1,01 | Presynaptic transmembrane protein with role in synapse development and function. | 34879268, 2847265 |
| PISD | 1,01 | Mitocondrial-localized enzyme, associated with impaired mitochondrial protein homeostasis. | 30858161 |
| Smarca2 | 1,08 | Subunit of chromatin remodelling complexes, involved in neural progenitor differentiation. | 31375262 |
| Ina | 1,16 | Neuronal intermediate filament protein that influence morphology and physiology of axonx. | 11739575 |
| Fxyd6 | 1,00 | Transmembrane protein involved in Na/K pump function modulation. | 33231612 |
| Zbtb20 | 1,11 | Transcription factor with role in neurogenesis and astrocytogenesis modulation. | 27282384, 27000654 |
| Gria2 | 1,15 | Subunit of AMPA receptor, LOF leads to neurodevelopmental disorders. | 31300657 |
| Socs2 | 1,19 | Intracellular protein that regulates neuron embryonic development and the neurotrophin signaling. | 12368809, 24860421 |
| Chl1 | 1,24 | Neural recognition molecule that begatively regulates neuronal proliferation and differentiation. | 20933598 |
| Pde4dip | 1,03 | Protein anchoring components of the cAMP-dependent pathway in the ER to Golgi trafficking. | 35346821 |
| Gria1 | 1,18 | Subunit of AMPA receptor, LOF leads to neurodevelopmental disorders. | 35675825 |
| Mef2c | 1,95 | Transcripton factor involved in interneurons fate and maturation and is linked to various neuropsychiatric and neurodevelopmental disorders. | 27779093, 18579729, 32452758 |
| Gucy1a3 | 1,41 | Subunit of guanylate cyclase enzyme regulating MGE neurons migration and marker of indirect striatal medium spiny neurons. | 24155296, 36081908 |
| Erbb4 | 1,00 | Receptor tyrosine kinase required for interneuron migration and inhibitory synapse formation. | 15473965, 34226493 |
| Flrt2 | 1,02 | cell adhesion molecules with role in Tangential Migratory Streams of Cortical Interneurons. | 34301831 |
| Shtn1 | 1,02 | Protein coding gene involved in neuronal polarization and axonogenesis. | 17030985, 30733148 |
| Lmo4 | 1,36 | Transcriptional regulator essential for normal patterns of proliferation ans survival of neuroepithelial cells. | 15691703 |
| Epha5 | 1,91 | Ephrin receptor protein, involved in axon guidance and synaptogenesis during development. | 19326470, 20824214 |
| Tcf4 | 1,32 | Transcription factor regulating cortical interneuron neurogenesis. | 35965434 |
| Maf | 1,34 | Transcription factor regulating cortical interneuorn fate and maturation. | 30699346, 32452758 |

**Table 1:** Functional summary of e16.5 enriched genes differentially expressed between $FT_{e12.5 + 6h}$ and $FT_{e16.5 + 6h}$.

| sgRNA name | Sequence | FW Primer | RV Primer | In vitro TIDE Knockout score (%) |
|---|---|---|---|---|
| gLacZ | GTGCGAATACGCCCACGCGAT | NA | NA | NA |
| gNfib | CTGGCGTCTGGATCTAGTCA | ACATGACCGTGGACCTGGA | GCTCAGTGAGAAGCCCGAAAT | 63 |
| gNfix | TTGGAAAGCACGCAGCAGGG | CTACCGGTGAGGCCCAGAAG | CAGGATGAGTTCCACCCGTT | 46 |

**Table 2:** Selected sgRNAs list with primers used to analyze CRISPR interference efficiency.
